# *C. elegans* “reads” bacterial non-coding RNAs to learn pathogenic avoidance

**DOI:** 10.1101/2020.01.26.920322

**Authors:** Rachel Kaletsky, Rebecca S. Moore, Geoffrey D. Vrla, Lance L. Parsons, Zemer Gitai, Coleen T. Murphy

**Affiliations:** Department of Molecular Biology; LSI Genomics, Princeton University, Princeton NJ 08544

## Abstract

C. elegans is exposed to many different bacteria in its environment, and must distinguish pathogenic from nutritious bacterial food sources. Here, we show that a single exposure to purified small RNAs isolated from pathogenic *Pseudomonas aeruginosa* (PA14) is sufficient to induce pathogen avoidance, both in the treated animals and in four subsequent generations of progeny. The RNA interference and piRNA pathways, the germline, and the ASI neuron are required for bacterial small RNA-induced avoidance behavior and transgenerational inheritance. A single non-coding RNA, P11, is both necessary and sufficient to convey learned avoidance of PA14, and its *C. elegans* target, *maco-1*, is required for avoidance. A natural microbiome *Pseudomonas* isolate, GRb0427, can induce avoidance via its small RNAs, and the wild *C. elegans* strain JU1580 responds similarly to bacterial sRNA. Our results suggest that this ncRNA-dependent mechanism evolved to survey the worm’s microbial environment, use this information to make appropriate behavioral decisions, and pass this information on to its progeny.

## Small RNAs from *P. aeruginosa* induce species-specific avoidance

*C. elegans* is surrounded by and consumes bacteria as its primary nutrient source. Its natural habitat contains many different bacterial species; about a third of these are in the *Pseudomonas* family (Samuel et al., 2016), which can be either beneficial or detrimental to the worms. Despite their natural attraction to pathogenic *Pseudomonas aeruginosa* (PA14), *C. elegans* can learn to avoid this pathogen after becoming ill (Zhang et al., 2005). Recently, we discovered that worms epigenetically pass on to their progeny this learned avoidance of PA14 (Moore et al., 2019), but the nature of the signal that conveys the identity of the pathogen was unknown. Therefore, we sought to determine the macromolecular nature of the bacterial cue to which they respond.

Although bacterial metabolites can serve as cues that alter worm behavior (Meisel et al., 2014), training worms with PA14 culture supernatant for 24h did not induce avoidance learning (Fig 1A). Next, we isolated nucleic acid components from pathogenic (25°C, plate-grown) PA14, added the purified samples to spots of OP50 *E. coli* to train the worms for 24h, and subsequently tested for their preference to OP50 *E. coli* vs PA14 (Fig 1B). We tested total DNA (Fig S1A), total RNA (Fig 1C), large RNA (>200nt) (Fig S1B), small RNA (<200nt), and RNase- and DNase-treated fractions (Fig 1D). We found that training on PA14 total RNA and small RNA, as well as DNase-treated small RNA samples (Fig 1C-D), induced avoidance of PA14, while DNA and large RNAs did not (Fig S1A-C). To determine whether bacterial metabolism is required for this effect, we trained worms on heat-killed OP50 *E. coli* bacteria supplemented with purified PA14 small RNAs, and still observed learned avoidance (Fig 1E), while small RNAs isolated from the less virulent *ΔlasR* mutant did not induce PA14 avoidance (Fig S1D). These results suggest that bacterial small RNAs are at least partially responsible for learned pathogenic avoidance.

**Figure 1:**
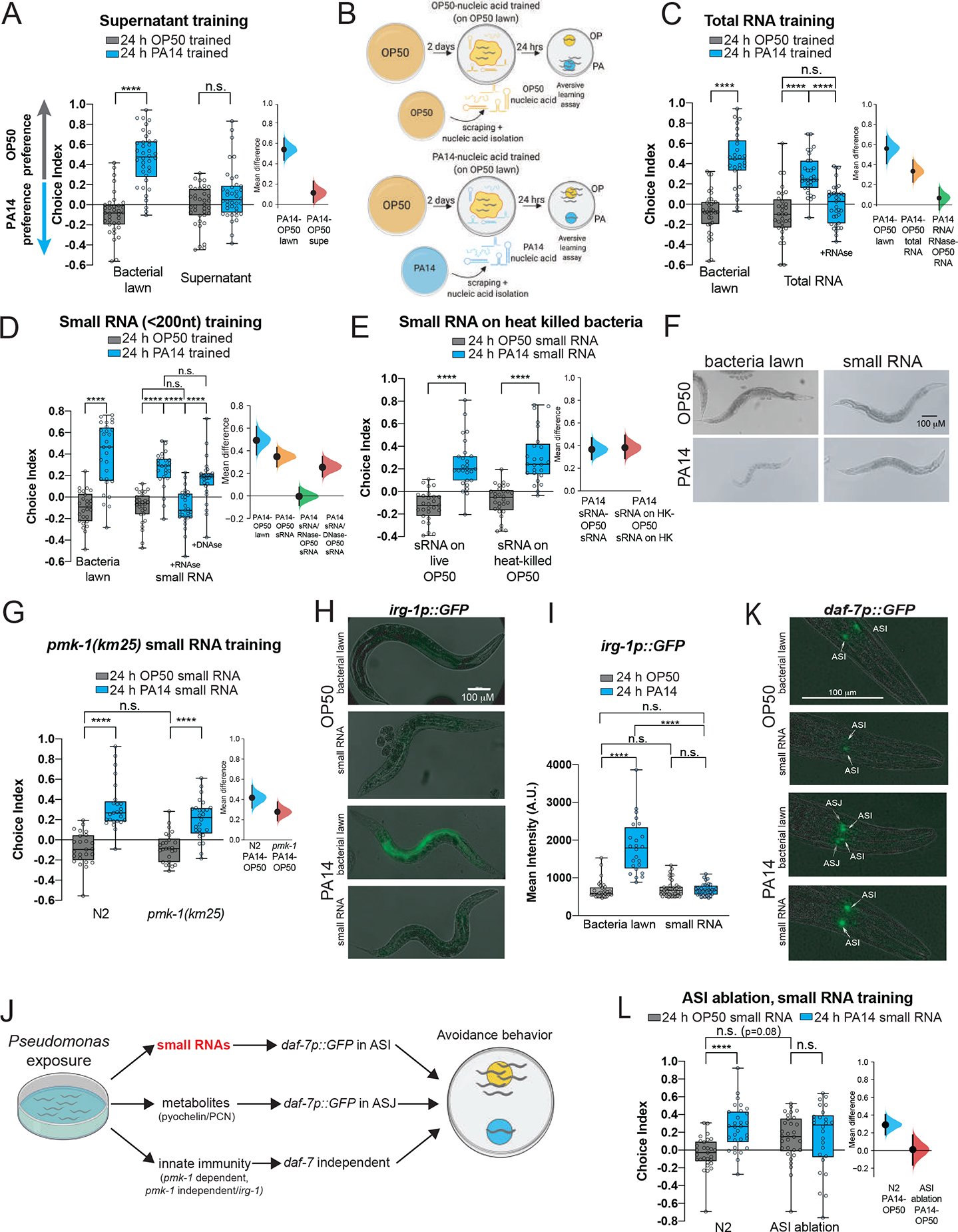
*Pseudomonas aeruginosa* (PA14) small RNAs are sufficient to induce *C. elegans* pathogen avoidance. (A) Worms exposed to a PA14 bacterial lawn for 24h learn to avoid PA14 in a choice assay, while training with PA14 supernatant is not sufficient to elicit PA14 avoidance. (B) Small RNA training protocol and aversive learning assay. Choice index = (# of worms on OP50 - # of worms on PA14)/(total # of worms). (C) Training with purified PA14 total RNA confers avoidance of PA14 similar to PA14 lawn exposure. RNA treatment with RNase prior to training abolishes this effect. (D) Purified PA14 sRNAs (<200 nt) induce learned avoidance of PA14. RNA treatment with RNase abolishes this effect, while treatment with DNase does not. (E) PA14 sRNA induces learning independent of bacterial metabolism. (F) *C. elegans* exposed to PA14 small RNAs for 24h are healthy compared to those exposed to PA14 bacterial lawns. (G) *pmk-1(km25)* is not required for PA14 sRNA-induced pathogenic learning. (H) *irg-1p::GFP* expression is induced by PA14 bacterial lawn exposure, but not by PA14 sRNAs alone. (I) GFP intensity from (H) was quantified, n ≥ 26 worms per group. (J) Worms learn to avoid *Pseudomonas* using several independent mechanisms. (K) *daf-7p*::*GFP* expression increases in the ASI neuron upon PA14 bacterial lawn or PA14 sRNA exposure (white arrows). *daf-7p::GFP* ASJ expression is induced only in PA14 bacteria-trained animals (grey arrowheads). (L) Worms with the ASI neurons genetically ablated fail to learn upon PA14 sRNA training. For all learning assays, each dot represents an individual choice assay plate (average of 115 worms/plate) with all data shown from at least 3 independent replicates. The box extends from the 25^th^ to 75^th^ percentiles, with whiskers from the minimum to the maximum values. One-Way (C-D) and Two-Way ANOVA (A, E, G, I, L), Tukey’s multiple comparison test. *p ≤ 0.05, **p ≤ 0.01, ***p ≤ 0.001, ****p < 0.0001, n.s. = not significant. Mean differences are shown using Cumming estimation plots (Ho et al., 2019), with each graphed as a bootstrap sampling distribution. Mean differences are depicted as dots; 95% confidence intervals are indicated by the ends of the vertical bars. Additional estimation plots for individual and pooled replicates are provided in Supplemental File 4. Imaging experiments (H-K) are representative results from 3 independent replicates.

### Small RNA avoidance does not require illness or the innate immune pathway

The discovery that purified small RNAs are sufficient to induce *P. aeruginosa* avoidance in the absence of the intact pathogen indicates that the avoidance induced by small RNAs does not require virulence per se, and the worms treated with small RNAs are healthy and their offspring develop normally (Fig 1F, S1E). Moreover, the innate immune pathway regulator *pmk-1* (Kim et al., 2002) is not required for the avoidance effect (Fig 1G, Fig S1F), nor is the canonical *pmk-1*-independent innate immune response (*irg-1p::gfp*) (Estes et al., 2010) induced by PA14 sRNA (Fig 1H-I). Therefore, small RNAs isolated from pathogenic PA14 replicate the parental avoidance behavior induced by the living pathogen without exposure to the bacteria themselves, illness, or induction of the innate immune response, suggesting that a distinct genetic pathway is required for the bacterial sRNA learned avoidance response (Fig 1J).

### *daf-7* expression in the ASI is induced by PA14 small RNA

The DAF-7 TGF-β ligand is induced in the ASI and ASJ neurons upon direct exposure to PA14 (Meisel et al., 2014), and *daf-7* is expressed at higher levels in the ASI neuron of F1 progeny of PA14-trained mothers (Moore et al., 2019). Surprisingly, PA14 sRNA exposure induced expression of *daf-7* solely in the ASI neurons, resembling the expression pattern of F1-F4 progeny of trained mothers (Moore et al., 2019), despite being the parental generation (Fig 1K, Fig S1G). Exposure to sRNA from a less virulent *Pseudomonas aeruginosa* mutant, *Δ*lasR, did not increase ASI *daf-7* levels compared to wild type PA14 (Fig S1H).

To assess the role of the ASI neuron in this process, we tested worms whose ASI neurons have been genetically ablated. While wild-type worms trained on a bacterial lawn learn to avoid PA14 (Moore et al., 2019), exposure of ASI-ablated worms to PA14 sRNAs does not induce avoidance (Fig 1L). These results suggest that the ASI neuron is required for bacterial sRNA-mediated PA14 avoidance (Fig 1J), just as it is required in the F1 generation for transgenerational inheritance of avoidance (Moore et al., 2019).

### The endogenous RNA interference pathway is required for small RNA-induced avoidance

We next wondered whether the components of the RNA interference (RNAi) pathway (Ghildiyal and Zamore, 2009) are required for pathogenic avoidance induced by PA14 sRNAs. RNAi was discovered in *C. elegans* (Fire et al., 1998) and is used widely to knock down gene expression via double-stranded RNAs homologous to endogenous sequences. Worms take up these artificially-expressed dsRNAs, then process and spread them across tissues, utilizing the endogenous RNAi system to silence genes for several generations. While none of these genes are required for avoidance induced by direct bacterial lawn exposure (Fig S2), sRNA-induced avoidance requires the SID-1 (Shih and Hunter, 2011) and SID-2 (Winston et al., 2007) dsRNA transporters (Fig 2A, Q; Fig S2A-B), as well as the RNAse III nuclease Dicer/DCR-1 that cleaves dsRNA (Bernstein et al., 2001; Ketting, 2001) (Fig 2B, Q; Fig S2C). *sid-1* mutants have high naïve PA14 avoidance relative to wild type, but still increase avoidance after lawn training (Fig S2A), showing that their innate immunity/metabolite responses are intact. The primary siRNA Argonaute RDE-1 (Tabara et al., 1999), the germline 22G siRNA biogenesis MUTator complex RDE-2/MUT-8 (Tops et al., 2005), and RDE-4, which binds dsRNA and mediates exo-RNAi (Tabara et al., 2002), have an abnormal response to control (OP50) sRNAs and show no further avoidance upon PA14 sRNA training (Fig S2D-F, 2Q), but have normal PA14 lawn learning and naïve behavior (Fig S2D-F). These results are consistent with *rde-4*’s reported roles in chemotaxis and response to *E. coli* (Liu et al., 2012; Posner et al., 2019). Together, these results suggest that components of the RNAi pathway are required for proper function of the bacterial small RNA response pathway.

**Figure 2:**
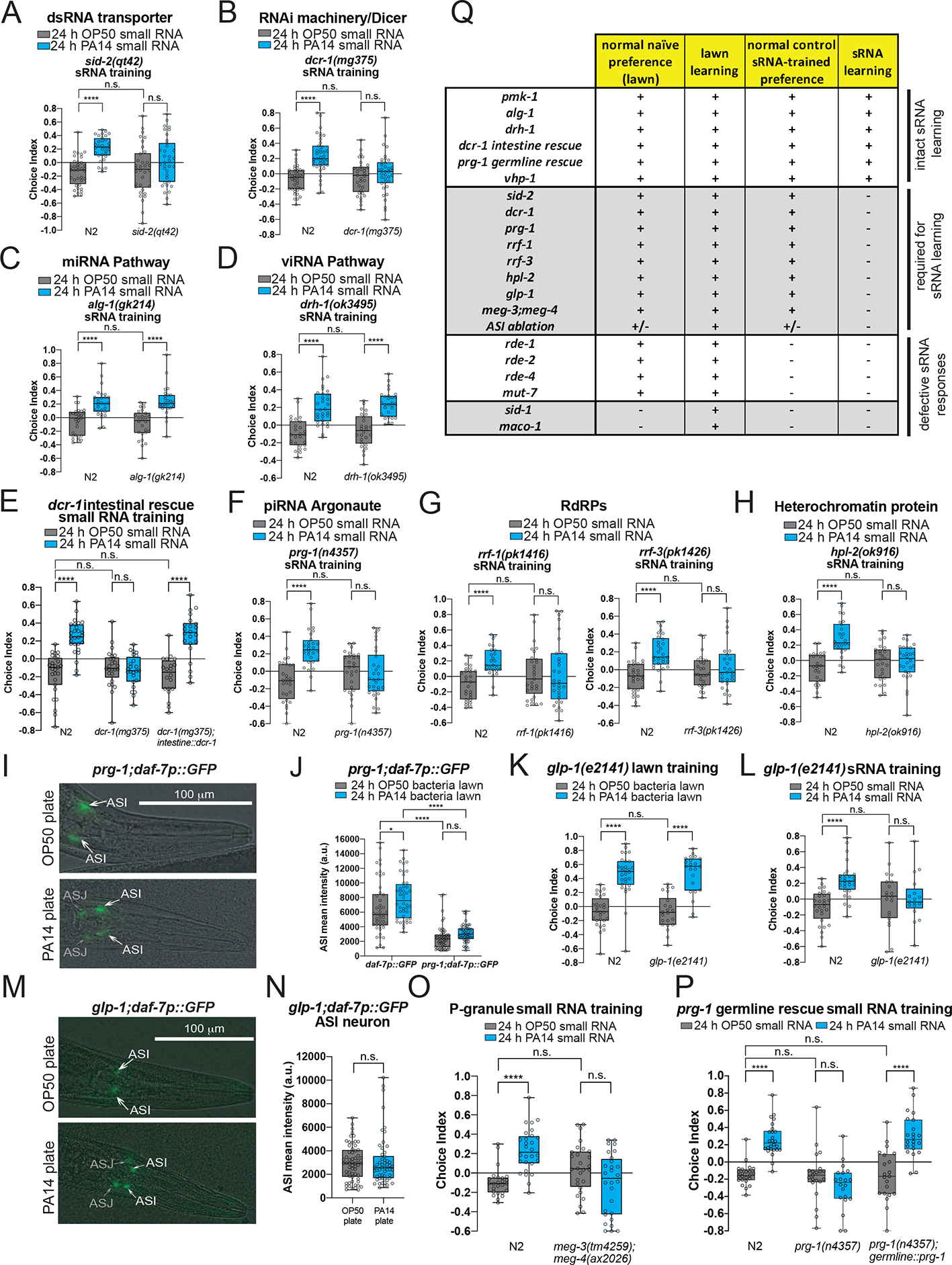
Worm dsRNA transport, processing machinery, piRNA Piwi/PRG-1 Argonaute pathway, and the germline are required for bacterial sRNA-induced pathogen avoidance Genes required. *(A) sid-2(qt42)*, (B) *dcr-1(mg375*), and not required (C) *alg-1(gk214*), (D) *drh-1(ok3495)* for sRNA-induced aversive learning. (E) Intestinal rescue of *dcr-1* (*vha-6p::dcr-1::dcr-1 3’UTR*) restores sRNA-mediated learning in *dcr-1* mutants. (F) *prg-1(n4357*), (G) *rrf-1(pk1417*), *rrf-3(pk1426*), and (H) *hpl-2(ok916*) mutants are deficient in PA14 sRNA-induced learned avoidance. (I-J) *prg-1* mutants induce *daf-7p::GFP* expression in the ASJ neuron, but not the ASI neuron, after PA14 bacterial lawn exposure (n ≥ 52 neurons per group). (K-L) *glp-1(e2141)* mutants lacking a germline can learn to avoid PA14 after training on a bacterial lawn, but are defective in sRNA-induced pathogenic learning. (M) Germline-less *glp-1* mutants induce *daf-7p::GFP* expression in the ASJ neuron, but not the ASI neuron (N), after PA14 lawn exposure. (n ≥ 52 neurons per group, Student’s t-test). (O) Germline P granule mutants exhibit defective sRNA-induced learning. (P) Germline *prg-1* rescue is sufficient to restore sRNA-induced learning. (Q) Summary of genes involved in naïve preference and small RNA learning. ((+) indicates statistically significant naïve preference or learning. (-) indicates defective naïve preference or learning. (+/-) indicates statistically borderline behavior). For choice assays and *prg-1;daf-7p::GFP* (J): Two-Way ANOVA, Tukey’s multiple comparison test. For all learning assays, each dot represents an individual choice assay plate (average of 115 worms/plate) with all data shown from at least 3 independent replicates. At least three biological replicates were performed for all experiments. *p ≤ 0.05, ****p < 0.0001, n.s. = not significant.

*C. elegans* uses similar machineries for responding to both exogenous siRNAs and *C. elegans-*produced microRNAs, but these processes utilize different Argonaute homologs: RDE-1 is specific for siRNAs, while *alg-1/*AGO-1 is specific for miRNAs. Whereas *rde-1* mutants disrupted sRNA-induced avoidance, mutants of the microRNA-specific Argonaute *alg-1*/AGO-1 (Grishok et al., 2001) were not defective for isolated sRNA-induced learned avoidance (Fig 2C, Q; Fig S2G). Similarly, mutants of the RIG-I-like RNA helicase *drh-1* mutants are competent for bacteria sRNA learning (Fig 2D, Q; Fig S2H), suggesting that the intracellular pathogen response to viral infection is not involved in *C. elegans’* learned avoidance of bacteria through its small RNAs (Ashe et al., 2015; Gammon et al., 2017; Tabara et al., 2002). Moreover, microRNA processing is unaffected in the *dcr-1(mg375)* mutant (Welker et al., 2010), which is unresponsive to sRNA from PA14 (Fig. 2B). Together, these data suggest that the secondary siRNA pathway, rather than the microRNA or viral processing pathways, mediates PA14 sRNA-induced avoidance learning.

### RNA interference pathway function in the intestine is necessary for sRNA-induced behavior

Knowing the site of RNA uptake and processing is critical for understanding the underlying mechanism of bacterial small RNA-induced avoidance. The SID-2 transmembrane protein is expressed in the intestine and is required for the import of intestinal dsRNA (McEwan et al., 2012; Winston et al., 2007) and is required for sRNA-induced learning (Fig 2A), while other components, including *dcr-1*, are expressed more broadly (Kaletsky et al., 2018). *dcr-1* rescue exclusively in the intestine was sufficient to restore small RNA-mediated learning (Fig 2E, Q; Fig S2I), suggesting that uptake and processing of bacterial small RNAs by SID-2 and DCR-1, respectively, are initiated in the intestine.

### The piRNA pathway is required for bacterial small RNA-induced avoidance

We showed previously that the piRNA regulator Piwi/PRG-1 Argonaute and its downstream components (MUT-7/RNAse D (Ketting et al., 1999), RRF-1/RNA-dependent RNA polymerase (Aoki et al., 2007), SET-25 histone methylase (Towbin et al., 2012), and the HPL-2 chromatin reader (Couteau et al., 2002)) are all required for the inheritance of learned pathogenic avoidance (Moore et al., 2019). However, these proteins were not required for maternal pathogen avoidance learning when trained on a PA14 bacterial lawn (Moore et al., 2019) (Fig S3A-F). Therefore, we were surprised to find that *prg-1* mutant mothers were defective specifically in PA14 sRNA-induced avoidance response (Fig 2F, Q). Furthermore, *mut-7, rrf-1*, *rrf-3* (an RNA-dependent RNA polymerase ((Smardon et al., 2000)) and *hpl-2* animals phenocopied the sRNA learning defect of *prg-1* mutants (Fig 2F-H, Q; Fig S3A-F). Consistent with this behavior, *prg-1* mutants exposed to PA14 bacterial lawns failed to upregulate *daf-7* in ASI (Fig 2I-J) but still induced *daf-7* expression in the ASJ neurons (Fig 2I). These data may explain why PA14 lawn-trained *prg-1* mothers still learn to avoid PA14 (Fig S3A), but do not pass on this learned information, and cannot learn when exposed to PA14 sRNAs alone (Fig 2F, Q).

### A functional germline is required for small RNA-induced avoidance

Because PRG-1 is required for *daf-7* expression changes in the ASI neurons, we next asked whether signaling through other tissues is required to induce gene expression in the nervous system and subsequent behavioral changes. *glp-1(e2141)* mutants lack a germline when grown at the restrictive temperature (25°C) (Austin and Kimble, 1987). Like wild type, these germline-less animals learn to avoid PA14 after training on a PA14 bacterial lawn (Fig 2K), demonstrating that learning via sRNA-independent innate immune pathways does not require a functional germline. However, *glp-1* mutants fail to exhibit sRNA-induced avoidance of PA14 (Fig 2L, Q). Consistent with the requirement of the germline for sRNA-mediated learning, germline-less *glp-1* worms were able to upregulate *daf-7* expression in the ASJ neurons (Fig 2M, Fig S3G) but not in the ASI neurons (Fig 2M, N) upon PA14 plate training. Furthermore, worms with defective germline granules (*meg-3;meg-4* mutants) (Ouyang et al., 2019) are also unable to induce avoidance in response to PA14 sRNA (Fig 2O, Fig S3H), despite having normal naïve preferences and lawn learning. Finally, germline-specific expression of *prg-1* fully rescued *prg-1’s* sRNA-induced PA14 avoidance (Fig 2P, Fig S3I). These results suggest that, while not needed for innate immunity-induced avoidance, a functional germline is required for sRNA induction of this neuronally-directed avoidance behavior, and that the piRNA pathway acts in the germline to do so. Thus, bacteria-derived small RNAs do not act directly in neurons, but rather through an indirect mechanism that first requires uptake by the intestine and then piRNA processing and P granule function in an intact germline to then communicate to the ASI neurons.

### Small RNA-induced behavior is transgenerationally inherited

We previously showed that PA14 lawn exposure induces heritable avoidance behavior that persists through the F4 generation (Moore et al., 2019). In light of the finding that PA14-derived sRNAs alone are sufficient for maternal learning, we examined the progeny of sRNA-trained mothers. Remarkably, we found that a single 24hr exposure of *C. elegans* to purified PA14 sRNA was sufficient to induce avoidance of PA14 not only in mothers, but also in the subsequent four generations (Fig 3A-B), completely mimicking the transgenerationally-inherited learned avoidance of training on pathogenic bacteria lawns that we previously identified (Moore et al., 2019). This avoidance effect persisted despite the fact that neither the mothers nor the progeny had ever directly encountered PA14. Transgenerational inheritance of avoidance for both bacterial lawn and sRNA exposure required the SID-1 and SID-2 dsRNA transporters (Fig S4A-B) and components of the RNAi machinery, including DCR-1 (Fig S4C). Together, these data suggest that the mechanism by which *C. elegans* transmits its learned avoidance does not depend on an innate immune response, or on molecules canonically sensed by Pattern Recognition Receptors (e.g., metabolites, proteins, lipopolysaccharides, etc.), but rather on *C. elegans’* ability to identify specific pathogens via bacterially-expressed sRNAs. Moreover, the similar magnitude of the behavior in progeny of live bacteria- and sRNA-trained animals suggests that the sRNA response is sufficient to fully explain the transgenerational change in behavior (Fig 3B).

**Figure 3:**
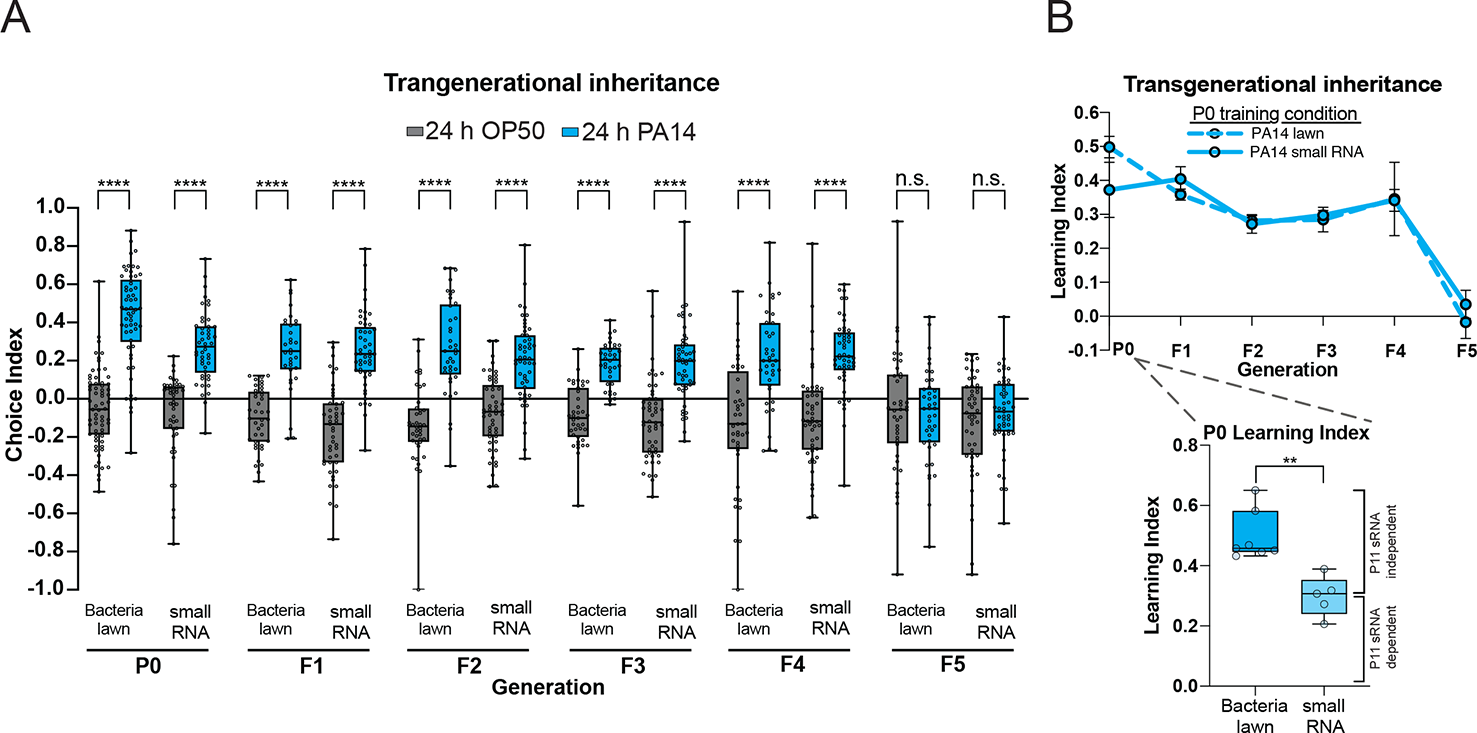
Small RNA-induced behavior is transgenerationally inherited. (A) Mothers exposed to PA14 small RNAs learn to avoid PA14 compared to OP50 sRNA-treated controls. Untrained (naïve) progeny of PA14 small RNA-trained mothers continue to avoid PA14 from generation F1-F4. 5^th^ generation progeny return to a state of PA14 attraction. Each dot represents an individual choice assay plate (average of 115 worms per plate) with all data shown from at least four biological replicates. Two-Way ANOVA, Tukey’s multiple comparison test. The learning index for each generation is shown in (B), mean ± SEM. (B, inset) The magnitude of small RNA learning is less than learning that occurs from PA14 lawn exposure. (Learning Index=Average naive choice index–Average trained choice index). Each data point is the learning index from an independent experiment containing ∼7-10 choice assay plates with an average of 115 worms per choice assay plate. Student’s t-test. For (A-B), each generation represents data pooled from at least 4 independent replicates. **p ≤ 0.01, ****p < 0.0001, n.s. = not significant.

Our data support a model in which pathogen avoidance is induced by at least two pathways: mothers worms respond acutely through the innate immune response (Troemel et al., 2006) to pathogen exposure and metabolites, which induces *daf-7* expression in the ASJ neuron (Meisel et al., 2014), resulting in about half of the avoidance behavior exhibited by trained worms (Fig 3B inset, P0). In a separate mechanism that we have identified here, pathogenic bacterial sRNAs are taken up and processed through the canonical RNA interference pathway via Dicer activity in the intestine. Subsequent germline signaling involving the piRNA pathway and epigenetic changes via HPL-2 (Moore et al., 2019) both induce *daf-7* expression in the ASI neuron, ultimately contributing to avoidance of PA14, and lead to transgenerational inheritance of the avoidance behavior signal.

### A single bacterial small RNA, P11, induces learned avoidance of PA14

*C. elegans* utilizes the information from pathogenic PA14 sRNA to induce avoidance, despite never having been sick. To determine whether specific sRNAs convey this information, first we tested the avoidance-inducing ability of sRNAs isolated from less-pathogenic growth conditions that do not induce pathogen avoidance, i.e., 15°C PA14 grown on plates (Moore et al., 2019), and PA14 grown under planktonic conditions (liquid-grown; Fig S5A). Learned avoidance was specific for sRNAs isolated from pathogenic conditions, that is, bacterial lawns of 25°C-plate grown PA14, as sRNAs isolated from less virulent 15°C-grown PA14 and liquid-grown PA14 did not induce subsequent avoidance of PA14 (Fig 4A-B).

**Figure 4:**
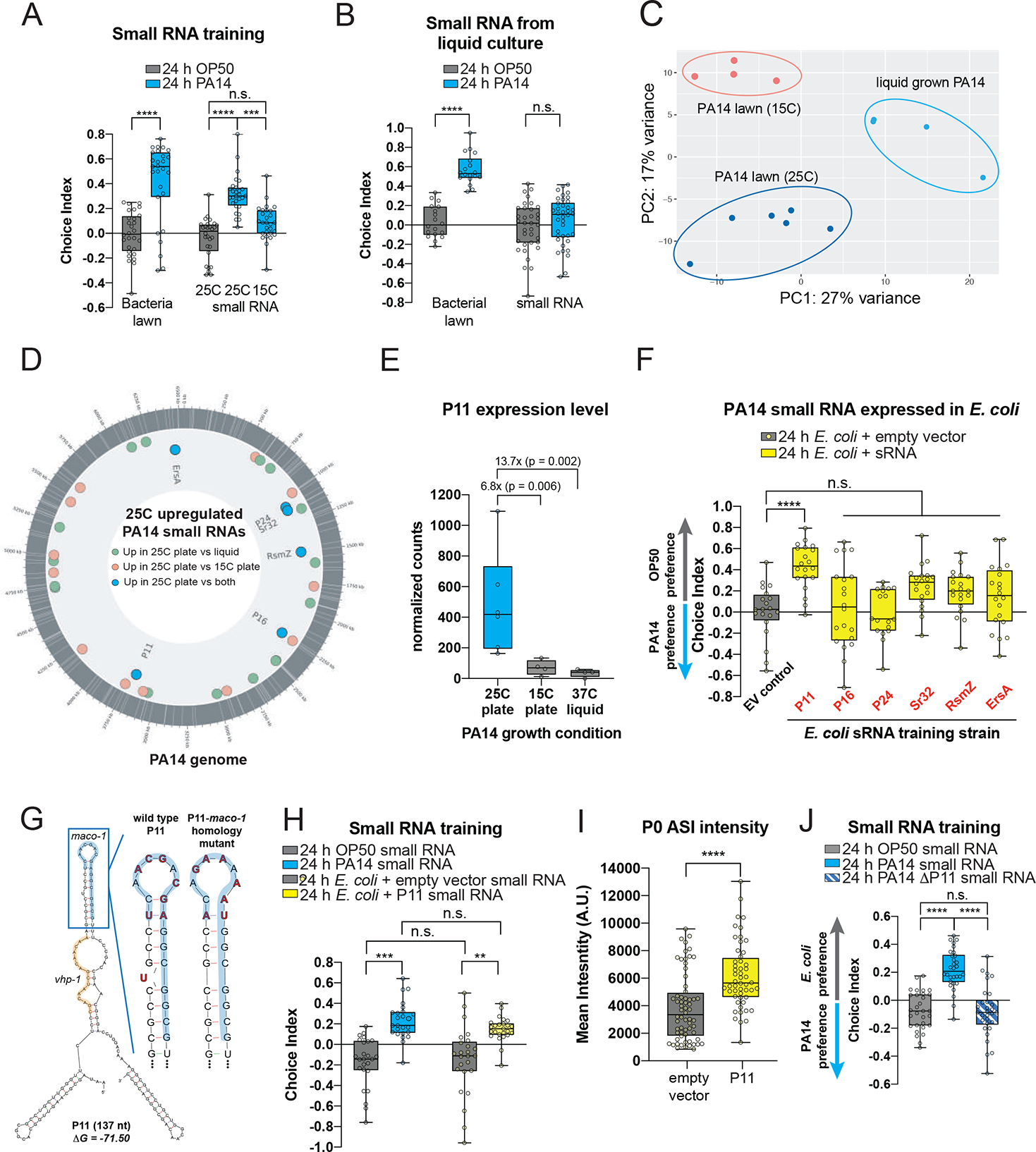
The PA14 P11 sRNA is required and sufficient for learned avoidance behavior (A-B) Small RNAs isolated from PA14 grown at 15°C. (A), or planktonic (liquid grown) PA14 (B) do not induce PA14 avoidance compared to worms trained on PA14 lawns or sRNAs from 25°C plate grown PA14. (C) Principal component analysis of RNA-seq samples from PA14 small RNA groups. (D) 25°C upregulated PA14 sRNAs compared to 15°C-grown and liquid-grown PA14 sRNAs. The outermost gray circle represents the PA14 genome, with previously identified sRNAs (Wurtzel et al., 2012) annotated with white lines. sRNAs that were upregulated (DESeq2, adjusted p-value ≤ 0.05) in the 25°C plate condition relative to 15°C plate grown PA14 (pink) or liquid grown PA14 (green) are indicated. Overlapping sRNAs upregulated in the 25°C plate condition relative to both other conditions are noted (blue dots). (E) DESeq2 normalized counts are shown for PA14 P11 expression. DESeq2 p-adj values are shown with average fold changes. (F) Worms were trained on *E. coli* (MG1655) containing empty vector (EV control) or *E. coli* (MG1655) expressing PA14 sRNAs (red). Following training, a standard OP50 vs PA14 choice assay was performed. Worms trained on PA14 or *E. coli* expressing P11 exhibited PA14 avoidance behavior. (G) Predicted secondary structure of wild-type P11 (G = −71.50) using mfold (Zuker, 2003). (G, inset) Wild-type bases in bold were mutated to disrupt *maco-1* homology but retain predicted P11 structure (G = −71.30). (H) Worms trained with sRNAs isolated from PA14 (blue) or *E. coli* expressing P11 (yellow) exhibit learned PA14 avoidance. (I) Training with *E. coli* expressing P11 induces *daf-7p::GFP* expression in the ASI neurons compared to *E. coli* empty vector-treated controls. (J) sRNA isolated from PA14-ΔP11 do not induce PA14 learned avoidance compared to PA14 sRNA trained worms. (For all learning assays, each dot represents an individual choice assay plate (average of 115 worms/plate) with all data shown from at least 3 independent replicates. Two-Way ANOVA, Tukey’s multiple comparison test. **p ≤ 0.01, ***p ≤ 0.001, ****p < 0.0001, ns = not significant.

We then sequenced the small RNAs isolated from PA14 under the different growth conditions and identified sRNAs that were differentially expressed (Fig 4C-D). Of the annotated PA14 sRNAs (Wurtzel et al., 2012), 18 and 22 were significantly more highly expressed in 25°C-grown PA14 compared to 15°C-grown and liquid-grown PA14, respectively (Tables 1-2). Of these, six sRNAs were upregulated in the 25°C-grown samples compared to both conditions: P11, P16, P24, PA14sr-032, ErsA, and PrrB/RsmZ (Fig 4D-E, Tables 1-2).

To test candidate sRNAs for effects on worm behavior, worms were trained on *E. coli* individually expressing each of the six PA14 sRNAs. While the other sRNAs had no significant effect on avoidance, exposure to *E. coli* expressing PA14 P11 was sufficient to induce PA14 avoidance behavior similar to animals treated with pathogenic PA14 bacteria (Fig 4F, Fig S5B-C). (Worms do not avoid the P11-expressing *E. coli,* eliminating P11 itself as the detected component (Fig S5D).) The function of P11 (Fig 4G), a 137-nt non-coding RNA specific to the *Pseudomona*s family (Livny, 2006), has not been studied in *P. aeruginosa*, but its *P. stutzeri* ortholog *nfiR* is required for nitrogen fixation and growth under stress conditions (Zhan et al., 2019). Worms treated with sRNAs isolated from P11-expressing *E. coli* also displayed PA14 avoidance (Fig 4H). Feeding worms P11-expressing *E. coli* induced *daf-7* expression in the ASI neuron (Fig 4I), similar to treatment with PA14 sRNAs (Fig 1J). To test the requirement of P11 in PA14 sRNA-induced learning, worms were treated with sRNAs isolated from a mutant PA14 strain in which P11 was deleted (PA14-*P11*) (Fig S5E). Worms exposed to PA14*-ΔP11* sRNAs did not acquire PA14 avoidance (Fig 4J). Together, these results demonstrate that P11 is both necessary and sufficient for PA14 avoidance learning, despite the fact that exposure to P11 ncRNA does not make worms ill, or affect their germlines or reproductive capacity (Fig S5F-G). Thus, *C. elegans* may have evolved detection of P11, which we find is both involved in PA14 pathogenesis (Fig S5H) and upregulated in *P. aeruginosa* grown in conditions that induce virulence (Fig 4E), as a biomarker of pathogenic PA14.

### P11 is required for the transgenerational inheritance of learned avoidance

Exposing mothers to *E. coli* expressing the PA14 P11 ncRNA caused transgenerational inheritance through the F4 generation, similar to the effect observed with PA14 and PA14 sRNA (Fig 5A-B). Inherited learned avoidance required P11, as a PA14*-ΔP11* lawn maintained the ability to induce learning in mothers, but failed to transmit learning to progeny (Fig 5C). Consistent with sRNAs being required for transgenerational avoidance, PA14 sRNA (Fig 5D-E) and PA14 P11 (Fig 5D, F) exposure to mothers induced *daf-7* expression in progeny, similar to PA14 lawn exposure (Meisel et al., 2014) (Fig 5D).

**Figure 5:**
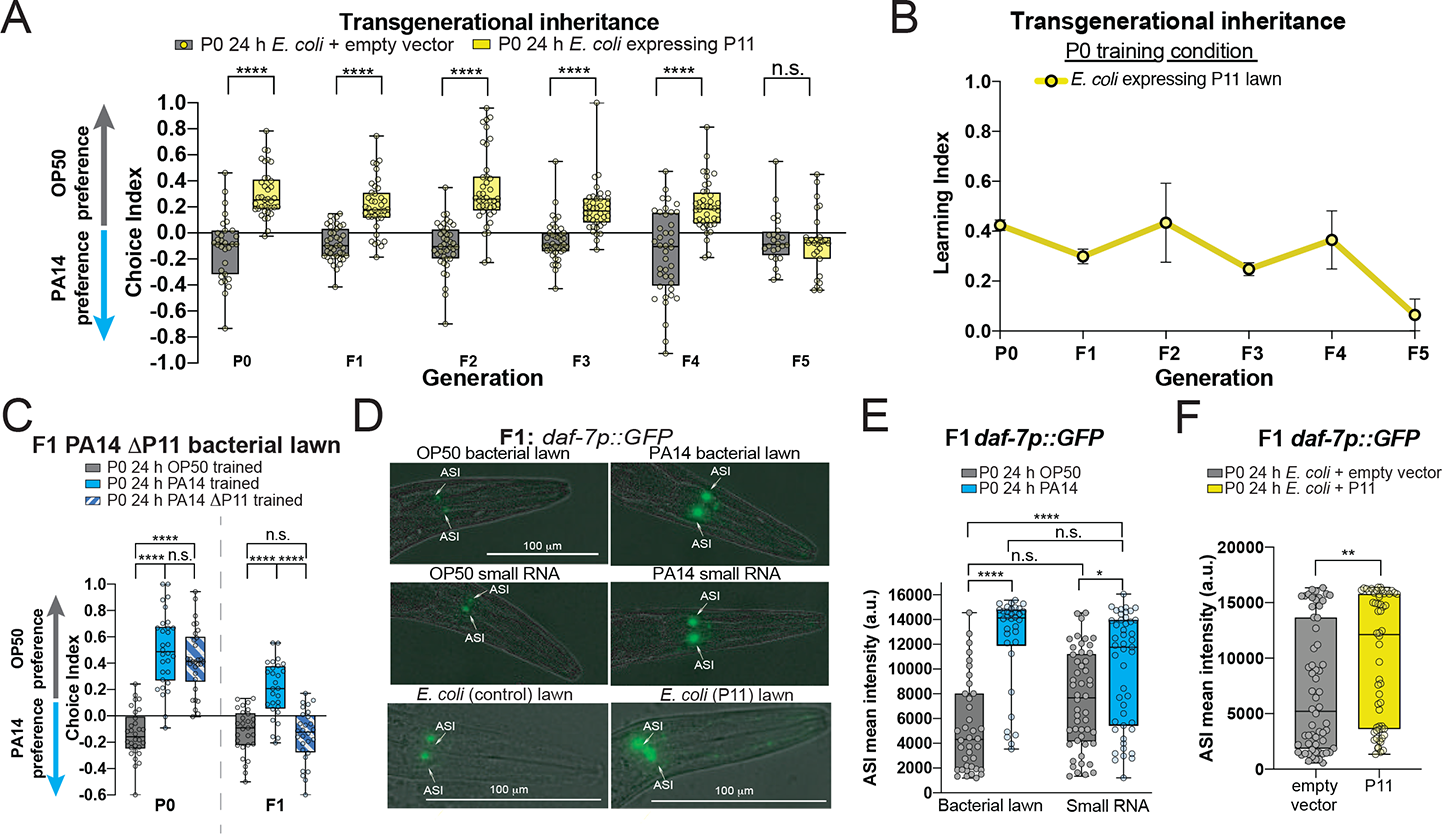
PA14 P11 small RNAs are sufficient to induce transgenerational pathogen avoidance. (A, B) Mothers exposed to lawns of *E. coli* expressing P11 learn to avoid PA14 compared to empty vector controls. Untrained (naïve) progeny of P11-trained mothers continue to avoid PA14 from generation F1-F4. 5^th^ generation progeny return to a state of PA14 attraction. The learning index for each generation is shown in (B), mean ± SEM. (C) P11 is required for inherited learned avoidance. Mothers trained on PA14-ΔP11 bacteria can learn to avoid PA14, but fail to transmit the learned avoidance to the next generation. (D) *daf-7p::GFP* expression remains elevated in the F1 progeny of PA14 bacterial lawn, PA14 sRNA-trained (E), and P11-trained (F, Student’s t-test) mothers. (For all choice assays and, each dot represents an individual choice assay plate (average of 115 worms/plate) with all data shown from at least 3 independent replicates. Two-Way ANOVA, Tukey’s multiple comparison test. *p ≤ 0.05, **p ≤ 0.01, ****p < 0.0001, n.s. = not significant.

### The macoilin MACO-1 is required for the small RNA-induced avoidance response

The presence of a behavioral response to a sequence-specific bacterial signal implies the existence of a *C. elegans* sequence that matches the bacterial sRNA. To find P11’s target(s), we analyzed the *C. elegans* genome for potential P11 homology, including mRNAs and non-coding RNAs (Table S3). The longest perfect match to P11 is a 17mer in *maco-1,* the *C. elegans* homolog of the mammalian neuronal gene Macoilin (Kuvbachieva et al., 2004) (Fig S6A). *maco-1* plays a role in chemotaxis, thermotaxis (Miyara et al., 2011), oxygen sensing, neuronal excitability (Arellano-Carbajal et al., 2011), and dauer formation, and is expressed in neurons, including the ASI (Neal et al., 2016). Upon PA14 exposure, *maco-1* expression is decreased in mothers and progeny (Fig. 6A), as one might expect for a target of P11. Therefore, we tested a *maco-1* mutant for its response to PA14 and sRNA. *maco-1* mutants exhibit high naïve preference for PA14 (Fig. 6B), and do not increase their PA14 avoidance upon sRNA treatment (Fig 6C). In contrast to wild-type worms (Fig 1K), expression of *daf-7p::gfp* in the ASI of *maco-1* mutants is not increased upon sRNA treatment (Fig 6D), suggesting that MACO-1 regulates *daf-7* expression. *maco-1’s* innate immune pathways are intact, as they increase avoidance to PA14 when exposed to PA14 bacteria (Fig 6E). Finally, training on *E. coli* expressing a multiple base substitution mutant of P11 that conserves P11 secondary structure but disrupts the perfect match to *maco-1* (Fig 4G) exhibited no learned avoidance (Fig 6F). By contrast, *vhp-1*, which has a 16nt match to P11 (Fig 4G), increases its expression in worms exposed to PA14 (Fig S6B), and loss of *vhp-1* had no effect on PA14 or sRNA-induced avoidance (Fig S6B-D, Fig 2Q). Additionally, progeny of PA14 lawn-exposed mothers inherited pathogen avoidance normally (Fig S6C) and expression of *vhp-1* is unchanged compared to progeny of OP50 treated mothers (Fig S6B). Therefore, *maco-1* is specifically required for response to PA14 sRNAs, likely through its sequence matching P11. *maco-1* mutants resemble some RNAi mutants (Fig 2Q) in that mothers exposed to PA14 lawns can increase avoidance, but there is no sRNA-mediated increased avoidance, and their naïve avoidance is already high (Fig 6B; 2Q).

**Figure 6:**
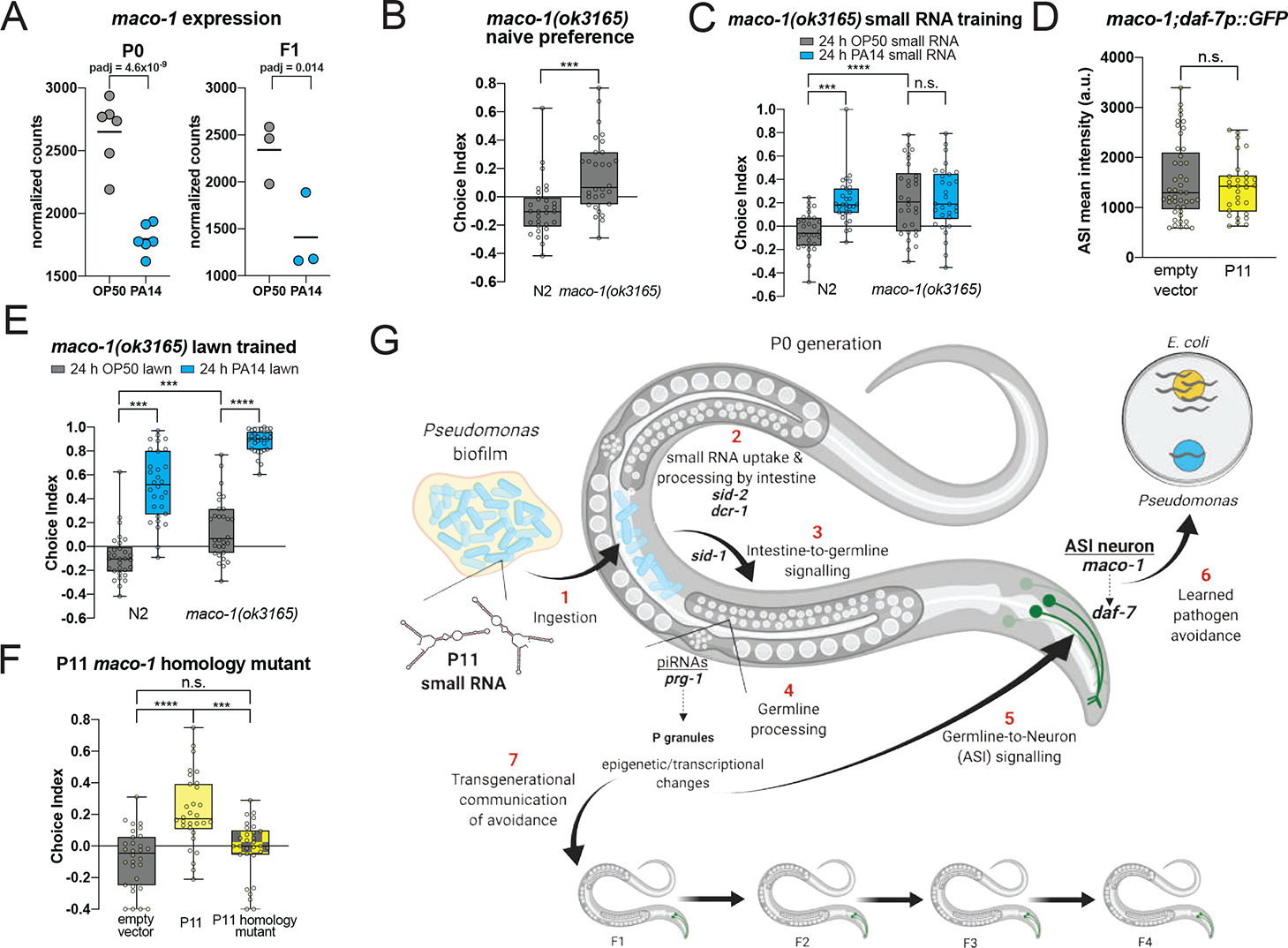
PA14 P11 small RNAs induces avoidance through its target, *maco-1*. (A) *maco-1* expression is reduced in PA14-exposed mothers and their naïve F1 progeny(Moore et al., 2019). DESeq2 p-adj values are shown. (B) Naïve choice preference of *maco-1(ok3165)* mutants compared to wild-type controls. (C) *maco-1* mutants do not exhibit increased avoidance of PA14 when trained with PA14 small RNAs. (D) *maco-1* mutants do not increase ASI *daf-7* expression upon *E. coli*-P11 training. (E) PA14 avoidance learning is intact in *maco-1* mutants trained on PA14 lawns. (F) Wild-type worms do not learn to avoid PA14 when trained with a P11 mutant in which the *maco-1* homology site is altered (Fig. 4G). One-way ANOVA, Tukey’s multiple comparison test. (G) Model of PA14 P11 sRNA-induced transgenerational learned avoidance. For choice assays, each dot represents an individual choice assay plate (average of 115 worms per plate) with all data shown from at least 3 independent replicates. Two-Way ANOVA, Tukey’s multiple comparison test. ***p ≤ 0.001, ****p < 0.0001, ns = not significant.

Together, our data suggest that the ingestion of PA14 and subsequent uptake and Dicer-mediated processing of the PA14 ncRNA P11 in the intestine, followed by further processing and piRNA regulation in the germline, leads to downregulation of *maco-1*, which in turn is required for upregulation of *daf-7* expression in the ASI and subsequent maternal avoidance behavior (Fig 6G).

### Small RNA-induced avoidance is species-specific

Like *Pseudomonas aeruginosa*, *Serratia marcescens* exposure also induces avoidance (Zhang et al., 2005); however, this avoidance is not passed on to the next generation, and PA14-induced transgenerational avoidance is species-specific (Moore et al., 2019). Therefore, we wondered whether small RNAs from *S. marcescens* could induce avoidance, and whether sRNA-induced avoidance is species-specific. We found that treatment with sRNA isolated from *S. marcescens* does not induce avoidance of *S. marcescens* or PA14, and that sRNA from PA14 does not induce avoidance of *S. marcescens* (Fig 7A-B). Therefore, like transgenerational inheritance of avoidance, sRNA-induced avoidance is not a generic response induced by all bacteria.

**Figure 7:**
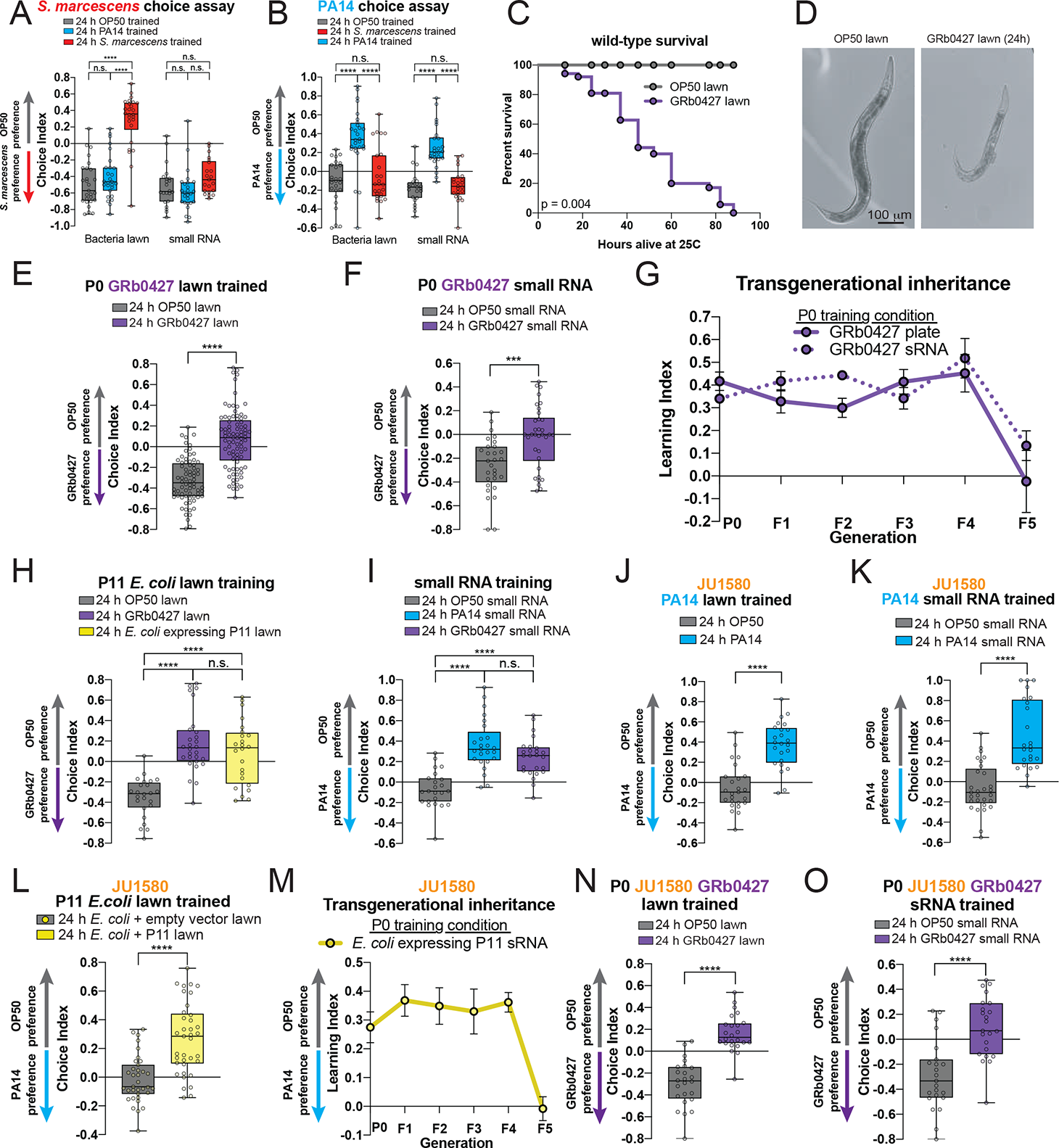
A wild *C. elegans* strain and natural bacterial isolate exhibit sRNA learning and transgenerational inheritance. (A) Worms trained on a lawn of *S. marcescens* (Sm) learn to avoid Sm, but exposure to Sm sRNAs does not induce Sm avoidance. (B) PA14 lawn and sRNA training, and not Sm lawn or Sm sRNA training, induce PA14 avoidance. (C) A pathogenic microbiome isolate, GRb0427 (Samuel et al., 2016), reduces *C. elegans*’ lifespan. Log-rank (Mantel-Cox test), n=120 animals/group. (D) DIC images of Day 1 worms treated with OP50 or GRb0427 from the L4 stage for 24h. (E) GRb0427 plate training and (F) GRb0427 sRNA training induces learned avoidance of GRb0427. (F) Untrained (naïve) progeny of GRb0427 plate- and sRNA-trained mothers continue to avoid GRb0427 from generation F1-F4. 5^th^ generation progeny return to a state of GRb0427 attraction. The learning index for each generation is shown, mean ± SEM. (H) P11-trained worms avoid GRb0427 similar to GRb0427 plated-trained worms. (I) GRb0427 sRNA-training induces PA14 avoidance similar to PA14 sRNA training. (J) PA14 lawn and (K) PA14 sRNA training induce PA14 avoidance in the wild *C. elegans* isolate JU1580 (Andersen et al., 2012). (L) P11-trained JU1580 worms avoid PA14, and untrained (naïve) progeny of P11-trained mothers continue to avoid PA14 from generation F1-F4. 5^th^ generation progeny return to a state of PA14 attraction. The learning index for each generation is shown. (N-O) JU1580 worms can learn to avoid GRb0427 when trained on GRb0427 lawns (N) or GRb0427 sRNA (O). For all learning assays, each dot represents an individual choice assay plate (average of 115 worms per plate) with all data shown from at least 3 independent replicates. Two-Way ANOVA, Tukey’s multiple comparison test. ***p ≤ 0.001, ****p < 0.0001, ns = not significant.

### A natural bacterial isolate from the worm’s microbiome induces avoidance learning

Our data suggest that *C. elegans* might have evolved a response to pathogenic bacteria that anticipates illness through its “reading” of the P11 ncRNA sequence and subsequent regulation of avoidance. If true, one might expect to find evidence of a similar phenomenon in wild isolates of *C. elegans* and in response to bacteria found in the natural microbiome of *C. elegans.* Recent 16s rRNA sequencing of the worm’s habitat revealed that *Pseudomonas* species make up about a third of *C. elegans’* natural ecosystem (Samuel et al., 2016). Like PA14, the *Pseudomonas* GRb0427 (Samuel et al., 2016) is first attractive to *C. elegans*, but after 24hr of exposure, which makes the worms sick and is eventually lethal (Fig 7C-D), *C. elegans* avoid GRb0427 (Fig 7E). Exposure to sRNA isolated from GRb0427 also induces avoidance (Fig 7F); furthermore, like PA14, both GRb0427 plate and sRNA training induces avoidance behavior that is passed on for four generations (Fig 7G). Training on P11-expressing *E. coli* also induces subsequent avoidance of GRb0427 (Fig 7H), and GRb0427 bacteria training induces avoidance of PA14 (Fig 7I), despite having never encountered PA14, suggesting that a P11-like sRNA from the currently unsequenced GRb0427 sRNA may mediate learned avoidance. sRNA-induced avoidance behavior and its transgenerational inheritance is caused by bacteria that are abundant in *C. elegans’* natural ecosystem, suggesting we have uncovered a behavior that is likely also functional in the wild.

### A wild *C. elegans* strain also learns to avoid *Pseudomonas* through small RNAs

To further test whether sRNA-induced bacterial avoidance learning occurs in nature, we subjected a wild strain of *C. elegans*, JU1580 (Andersen et al., 2012), to PA14 bacteria, PA14 sRNAs, and P11-expressing *E. coli*. JU1580 exhibits PA14 avoidance after training with each (Fig 7J-L), and JU1580 transgenerationally transmits P11-induced avoidance for four generations (Fig 7M). These findings suggest that the sRNA “reading” pathway is functional in this wild strain of *C. elegans*. This is particularly interesting because this strain has been shown to be defective for long viral dsRNA processing via *drh-1* (Ashe et al., 2013), which agrees with our finding that DRH-1 is dispensable for sRNA-induced learning (Fig 2D). Moreover, JU1580 learns to avoid the natural microbiome isolate GRb0427 (Fig 7N), and can do so when exposed to GRb0427’s isolated sRNA (Fig 7O). Therefore, all the aspects of the sRNA-based bacterial avoidance learning that we have described here are conserved in a wild strain of *C. elegans* and a wild bacterial isolates from its natural environment. The fact that natural isolates of both *C. elegans* and its microbiome *Pseudomonas* mimic the phenomenon we have characterized suggests that worms have evolved the ability to “read” ncRNAs that are biomarkers of virulence, enabling the worms to predict infection and enable a large fraction of the population to avoid the pathogen for several generations.

## DISCUSSION

### *C. elegans* can identify bacteria by “reading” their small RNAs

Cellular responses to exogenous RNA were discovered over 20 years ago in *C. elegans* (Fire et al., 1998), but the natural context in which worms evolved the ability to detect and respond to non-viral RNA remains unclear (Braukmann et al., 2019). Here we have identified a trans-kingdom signaling system that uses components of the RNAi pathway to “read” bacterial small RNAs, particularly P11, which is required for surface-growth of PA14 (Chuang et al., 2019) and virulence (our results). This small RNA-sensing pathway depends on processing through the germline and subsequent communication to neurons, and is independent of the pathways induced by innate immune systems and secreted molecules (Meisel et al., 2014; Papenfort and Bassler, 2016). Whether other animals utilize their RNA interference pathways for similar purposes, such as mammalian intestinal cells exposed to the microbiome, will be interesting to explore.

### The physiological relevance of small RNA-induced pathogen avoidance

Previously, trans-kingdom signaling has been reported in which small RNAs of a pathogen hijack the host immune system to avoid detection (Li et al., 2014; Weiberg et al., 2013). By contrast, we have found that *C. elegans* uses detection of specific bacterial small RNAs to mount a species-specific avoidance response that is propagated for several generations. While other previously-described trans-kingdom signaling systems are beneficial to the pathogen, *C. elegans* identifies small RNAs of pathogens in order to protect itself and to induce a search for less-pathogenic food sources. The phenomenon described here may explain why both systemic and TEI responses to exogenous RNA exist: to modify the worm’s behavior in response to encounters with naturally abundant and pathogenic bacterial species, which are identified by the worm through unique small RNA signatures. The fact that this avoidance endures for several generations suggests that an acute response is not sufficient to protect against some abundant pathogens, but must be transient enough to be reset in order to prevent account for changes in available bacterial food quality.

The process we have uncovered here is also distinct from responses to prolonged, multi-generational bacterial exposure that induce dauer formation (Palominos et al., 2017; Moore and Murphy, unpublished data) and infection adaptation (Burton et al., 2019). For example, when worms are forced to encounter pathogenic *Pseudomonas* continuously (>6 days) over multiple generations, eventually their progeny halt development at the alternative, pre-reproductive L3 larval stage known as dauer, undergoing extensive remodeling of their cuticle, mouths, and other tissues. Dauer remodeling is extremely energy-intensive and reproductively unfavorable - a last-ditch effort to survive under extreme conditions - which may be why it requires such a prolonged exposure to pathogens to induce this extreme response. By contrast, 24h of exposure to pathogenic *Pseudomonas* is sufficient to cause worms to switch their behavior from attraction to avoidance, and to induce the transgenerational inheritance of this learned avoidance. Rapid and lasting behavioral changes may be a more efficient and reproductively advantageous mechanism to force the worms to escape and thus explore a new and potentially safe environment. By passing this avoidance behavior on to several generations of their progeny, they may spare them from ever experiencing a prolonged exposure to the same pathogen, despite its potential abundance in the environment. Such a species-specific and plastic response may provide worms with a powerful survival mechanism that is fast-acting and potentially rapidly reversible (Burton et al., 2019), a first line of defense against pathogens.

## Conclusion

Here we have found that small, non-coding RNAs expressed in bacteria are sufficient to induce avoidance behavior in *C. elegans*. This was surprising, as the mechanism does not require sensing of bacterial metabolites, or the induction of a pathogen response. Instead, *C. elegans* appear to be able “read” ncRNAs that are specific to the bacterium in its pathogenic state. Bacterial small RNAs act as a decipherable code that allows the worms and their progeny to identify pathogens and respond appropriately when confronted with the pathogen in the future, prior to actually becoming ill. That is, the progeny already know to avoid these pathogens not because they have become sick, but because they have been “vaccinated” against the pathogen. This nascent adaptive immune system may enable *C. elegans* to identify and avoid pathogenic bacteria that exist in high abundance in their microbiome, and to pass on this avoidance to several generations of progeny. This trans-kingdom communication paradigm may represent a form of *C. elegans*’ adaptive immune memory that prepares future generations for encounters with harmful environmental conditions, allowing them to properly respond to a pathogenic threat. Our results suggest that a physiological role for the RNA interference pathway may in fact be to allow worms to survey their microbial environment, providing them with information that they not only utilize themselves, but also pass on to several generations of progeny.

## Supporting information

Supplemental File 4 Cumming Estimation Statistics

Table S1 DESeq_sRNA_25plate_vs_15_plate

Table S2 DESeq_sRNA_25plate_vs_liquid

Table S3 P11 homology to C. elegans coding and ncRNA

## Materials and Methods

### C. elegans and bacterial strains cultivation

Worm strains were provided by the *C. elegans* Genetics Center (CGC). KU25: *pmk-1(km25)*, AU133: *agls17[Pmyo-2::mCherry + Pirg-1::GFP]*, PY7505: oyIs84 [*gpa-4p*::TU#813 + *gcy-27p*::TU#814 + *gcy-27p*::GFP + *unc-122p*::DsRed], NL3321: *sid-1(pk3321),* HC271: ccIs4251[(pSAK2) myo-3p::GFP::LacZ::NLS + (pSAK4) myo-3p::mitochondrial GFP + dpy-20(+)] I. qtIs3[myo-2p::GFP dsRNA hairpin]. mIs11 [myo-2p::GFP + pes-10p::GFP + gut-promoter::GFP], YY470: *dcr-1(mg375)*, WM27: *rde-1(ne219),* NL3531: *rde-2(pk1657),* WM49: *rde-4(ne301),* COP2012: *alg-1(knu867),* RB2519: *drh-1(ok3495),* CQ636 *dcr-1(mg375); vha-6p::dcr-1::dcr-13’UTR* + *[Pmyo-2p::mCherry],* MAH23: *rrf-1(p1417)*, NL2099: *rrf-3(ok1426)*, NL917: *mut-7(pk204)*, SX922: *prg-1(n4357)*, RB995: *hpl-2(ok916),* CQ605 *prg-1(n4357); ksIs2 [Pdaf-7p::GFP + rol-6(su1006)],* CF1903: *glp-1(e2141)*, CQ640: *glp-1(e2141); ksIs2 [Pdaf-7p::GFP + rol-6(su1006)]* (strain was made by mating CF1903 with FK181), YL243: *unc-119(ed) III; vrls79 [pie-1p::GFP::prg-1 + unc-119(+)],* CQ655: *prg-1(n4357); unc-119(ed) III; vrls79 [pie-1p::GFP::prg-1 + unc-119(+)]* (strain was made by mating SX922 with YL243), JH3225: *meg-3(tm4259); meg-4(ax2026)*, FK181: *ksIs2 [Pdaf-7p::GFP + rol-6(su1006)]*, RB2329: *maco-1(ok3165)*, CQ654:*ksIs2[Pdaf-7p::GFP + rol-6(su1006)]*; *maco-1(ok3165)* (this strain was made by mating FK181 with RB2329), JT366: *vhp-1(sa366)*.

### Bacterial strains

*P. aeruginosa* PA14, and *P. aeruginosa* Δ*lasR* were gifts from Z. Gitai. GRb0427 was a gift from B. Samuel. OP50 was provided by the CGC. *S. marcescens* (ATCC 274) was purchased from ATCC.

### General worm maintenance

Worm strains were maintained at 15°C on High Growth Media (HG) plates (3 g/L NaCl, 20 g/L Bacto-peptone, 30 g/L Bacto-agar in distilled water, with 4 mL/L cholesterol (5 mg/mL in ethanol), 1 mL/L 1M CaCl2, 1 mL/L 1M MgSO4, and 25 mL/L 1M potassium phosphate buffer (pH 6.0) added to molten agar after autoclaving) on *E. coli* OP50 using standard methods.

### General bacterial cultivation

OP50, *P. aeruginosa* PA14, *P. aeruginosa* Δ*lasR, S. marcescens,* and GRb0427 were cultured overnight in autoclaved and cooled Luria Broth (10 g/L typtone, 5 g/L yeast extract, 10 g/L NaCl in distilled water) shaking (250 rpm) at 37°C. *E. coli* strains expressing PA14 small RNAs were cultured overnight in Luria Broth supplemented with 0.02% arabinose w/v and 100 μg/mL carbenicillin.

## Training plate/worm preparation

### Worm preparation

Eggs from young adult hermaphrodites were obtained by bleaching and subsequently placed onto HG plates and incubated at 20°C for 2 days. Synchronized L4 worms were used in all training experiments. For experiments involving CF1903 (*glp-1(e2141))*, eggs from mutant and wild-type adult hermaphrodites were obtained by bleaching and placed onto High Growth (HG) plates and left at 25°C for 2 days. Germline loss was confirmed in adult *glp-1(e2141)* worms raised at 25°C.

### Bacteria lawn (25°C) training plate preparation

Overnight cultures of bacteria (prepared as described above) were diluted in LB to an Optical Density (OD600) = 1 and used to fully cover Nematode Growth Media (NGM) ((3 g/L NaCl, 2.5 g/L Bacto-peptone, 17 g/L Bacto-agar in distilled water, with 1 mL/L cholesterol (5 mg/mL in ethanol), 1 mL/L 1M CaCl2, 1 mL/L 1M MgSO4, and 25 mL/L 1M potassium phosphate buffer (pH 6.0) added to molten agar after autoclaving) plates. For preparation of *E. coli* expressing PA14 small RNAs, bacteria were seeded on NGM plates supplemented with 0.02% arabinose and 100 μg/mL carbenicillin. All plates were incubated for 2 days at 25°C unless specified otherwise (in separate incubators for control/pathogen seeded plates). On the day of training (i.e., 2 days post bleaching), plates were left to cool on a benchtop for 1 hr to equilibrate to room temperature before the addition of worms. Additionally, for *E. coli* strains expressing PA14 sRNAs, 200 μL of 0.01% arabinose was spotted onto seeded training plates 1 hr prior to use.

### Bacteria lawn (15°C) training plate preparation

PA14 was prepared by centrifuging 5 mL overnight cultures for 10 minutes at 5000 x g. The supernatant was removed, and the remaining pellet was resuspended in 5 mL of fresh LB. Washed bacteria were used to inoculate (1:500) fresh LB to grow at 15°C for 2 days. Cultures were diluted in LB to an OD600 = 1 and used to seed NGM plates. Plates were incubated at 15°C for 2 days.

### DNA/Supernatant/Small RNA training plate preparation

200 μL of OP50 was spotted in the center of a 10 cm NGM. Plates were incubated at 25°C for 2 days.

### Heat-killed bacteria training plate preparation

One day before plate use, overnight bacteria cultures of OP50 were centrifuged at 5,000 x g for 10 minutes. Following centrifugation, pellets were resuspended in 1/10 volume of LB. Resuspended bacteria pellets were heat shocked at 95°C for 1 hr. Heat shocked bacterial suspensions were left to cool at room temperature for 1 hr, and 200 μL of heat killed bacteria was spotted in the middle of a 10 cm NGM plate supplemented with 100 μg/mL carbenicillin. Plates were incubated at 25°C for 1 day, prior to use. No bacterial growth was observed on heat-killed bacteria plates both before worm training and 24h after worm training.

### DNA preparation and training

Overnight cultures were pelleted at 5,000 x g for 10 minutes at room temperature. DNA was prepared from pelleted bacteria according using the Qiagen DNeasy Blood and Tissue kit and subsequently used fresh. 10 ng of bacterial DNA was placed onto *E. coli* spots and left to completely dry at room temperature for approximately 1 hr before the addition of worms for training.

### Supernatant

Overnight bacterial cultures (undiluted) were pelleted at 5,000 x g for 10 minutes at room temperature. Supernatant was collected and filtered using a 0.22 μm syringe filter. For worm training plates, 1 mL of filtered supernatant was put onto OP50 spots and left to completely dry at room temperature for approximately 1 hr before the addition of worms for training.

## Preparation of bacteria for RNA isolation

Bacteria for RNA collection were prepared as described for training plates (i.e. 2 days on plates at 25°C). Bacterial lawns were collected from the surface of NGM plates using a cell scraper. Briefly, 1 mL of M9 buffer was applied to the surface of the bacterial lawn, and the bacterial suspension following scraping was transferred to a 15 mL conical tube. PA14, Δ*lasR*, *S. marcescens*, GRb0427, or *E. coli* expressing PA14 sRNA strains from 10 plates or OP50 from 15 plates was pooled in each tube and pelleted at 5,000 x g for 10 minutes at 4°C. The supernatant was discarded and the pellet was resuspended in 1 mL of Trizol LS for every 100 mL of bacterial pellet recovered. The pellet was resuspended by vortexing and subsequently frozen at −80°C until RNA isolation.

## Bacteria RNA isolation

To isolate RNA from bacterial pellets, Trizol lysates were incubated at 65°C for 10 min with occasional vortexing. Debris was pelleted at 7000 x g for 5 min at 4°C. The supernatant was transferred to new tubes containing 1/5 the volume of chloroform. Samples were mixed thoroughly by inverting and centrifuged at 12000 x g for 10 min at 4°C. The aqueous phase was used at input for RNA purification using the mirVana miRNA isolation kit according to the manufacturer’s instructions for total RNA, large RNA (>200 nt), or small RNA (<200 nt) isolation. Purified RNA was used immediately or frozen at −80°C until further use.

### RNase treated samples

For RNase treatment of purified RNA, samples containing 100 μg of RNA were treated with 2.5 μL of an RNAse A (500 U/mL) and RNase T (20,000 U/mL) cocktail for every 50 μL of RNA (RNase Cocktail Enzyme Mix, Ambion). Samples were incubated at room temperature for 20 minutes before adding to worm training plates seeded with OP50. RNase degradation was confirmed using an Agilent 2100 Bioanalyzer. 100 μg of purified small RNA (measured prior to RNAse degradation) treated with RNase was spotted onto plates prior to use for worm training.

### DNase treated samples

For DNase treatment, samples containing 100 μg of purified RNA were treated with 2U of DNase I per 10 mg of RNA using the Invitrogen DNA-free kit according to the manufacturer’s instructions. 100 μg of purified small RNA treated with DNase was spotted onto plates prior to use for worm training.

### Total/small RNA/small RNA on heat killed bacteria

240 μg of total RNA, or 100 μg of small or large, or RNase/DNase-treated small RNA was placed directly onto OP50 spots and left to completely dry at room temperature (∼ 1 hr) before use on day of experiment for worm training.

## Worm preparation for training

Synchronized L4 worms were washed off plates using M9 and left to pellet on the bench top for approximately 5 minutes. 5 μL of worms were placed onto small RNA-spotted training plates, while 10 μL or 40 μL of worms were plated onto OP50 or *E. coli* expressing PA14 small RNAs, or pathogen-seeded training plates, respectively. Worms were incubated on training plates at 20°C in separate containers for 24 hr. After 24 hr, worms were washed off plates using M9 and washed an additional 3 times to remove excess bacteria. Worms were tested in an aversive learning assay described below.

## Aversive learning assay

Overnight bacterial cultures were diluted in LB to an Optical Density (OD_600_) = 1, and 25 μL of each bacterial suspension was spotted onto one side of a 60 mm NGM plate and incubated for 2 days at 25°C. After 2 days assay plates were left at room temperature for 1 h before use. Immediately before use, 1 μL of 1M sodium azide was spotted onto each respective bacteria spot to be used as a paralyzing agent during choice assay. To start the assay (modified from (Zhang et al., 2005)), worms were washed off training plates in M9 allowed to pellet by gravity, and washed 2 additional times in M9. 5 μL of worms were spotted at the bottom of the assay plate, using a wide orifice tip, midway between the bacterial lawns. Aversive learning assays were incubated at room temperature for 1 hr before manually counting the number of worms on each lawn. Plating a large number worms (>200) on choice assay plates was avoided, since excess worms clump at bacterial spots making it difficult to distinguish animals, and high densities of worms can alter behavior.

In experiments in which F1 and subsequent generations are used: Day 1 worms after from parental (P0) training were bleached and eggs were placed onto HG plates and left for 3 days at 20°C. All animals tested are washed off HG plates with M9 at Day 1. Some of the pooled animals were subjected to an aversive learning assay, while the majority of worms were bleached to obtain eggs, which were then placed onto HG plates left at 20°C for 3 days and used to test F2s.

## Imaging and fluorescence quantitation

All images were taken on a Nikon Eclipse Ti microscope. DIC images of whole worms following OP50, PA14, or GRb0427 lawn or small RNA training were imaged at 20X.

Z-stack multi-channel (DIC, GFP) of day 1 adult GFP transgenic worms were imaged every 1 μm at 60X magnification; Maximum Intensity Projections and 3D reconstructions of head neurons were built with Nikon *NIS-Elements*. To quantify *daf-7p*::*GFP* levels, worms were prepared and treated as described for pathogen training. Worms were mounted on agar pads and immobilized using 1 mM levamisole. GFP was imaged at 60X magnification and quantified using *NIS-Elements* software. Average pixel intensity was measured in each worm by drawing a bezier outline of the neuron cell body for 2 ASI head neurons and/or 2 ASJ head neurons.

For *irg-1p::GFP* quantification, whole worms were prepared as described above and imaged at 20x magnification. Average pixel intensity was measured in each worm by drawing a bezier outline the entire worm.

## Brood size assay

L4-stage mothers were trained for 24 h on control (empty vector) or P11-expressing *E. coli* as described above. After 24 h, 15 individual worms for each condition were transferred to NGM plates seeded with OP50. Worms were transferred to fresh plates every 24 h. Progeny containing plates were incubated at 20°C for 2 days before progeny were counted. Worms were moved daily until progeny production ceased.

## Progeny development assay

Mothers were trained on OP50 or PA14 small RNAs as described above. After 24 h of training, mothers were bleached and progeny were transferred to OP50-seeded NGM plates. Plates were incubated at 20°C for 2 days before progeny were assayed for developmental stage.

## Small RNA sequencing

Each sample of PA14 small RNA was tested for *C. elegans* behavior prior to sequencing. The size distribution of small RNA samples was examined on Bioanalyzer 2100 using RNA 6000 Pico chip (Agilent Technologies, CA). For small RNA-seq, around 300 nanograms of small RNA from each sample was first treated with RNA 5’ Pyrophosphohydrolase (New England Biolabs, MA) at 37°C for 30 minutes, then converted to Illumina sequencing libraries using the PrepX RNA-seq library preparation protocol on the automated Apollo 324^TM^ NGS Library Prep System (Takara Bio, CA). Briefly, the treated RNA samples were ligated to 2 different adapters at each end, then reverse transcribed to cDNA and amplified by PCR using different barcoded primers. The libraries were examined on Bioanalyzer DNA High Sensitivity chips (Agilent, CA) for size distribution, quantified by Qubit fluorometer (Invitrogen, CA), then pooled at equal molar amount and sequenced on Illumina NovaSeq 6000 S Prime flowcell as single-end 122 nt reads. The Pass-Filter (PF) reads were used for further analysis.

## Small RNA analysis

*Pseudomonas aeruginosa* (UCBPP-PA14) small RNA stranded reads were trimmed to remove adapters using Cutadapt (v1.16.6). Reads were mapped to the CP000438.1 genome using BWA-MEM. For small RNA analysis, count tables were generated using previously annotated intergenic non-coding RNAs (sRNA) (Wurtzel et al., 2012). Differential gene expression between 25°C plate vs 15°C plate and 25°C plate vs liquid conditions was performed using DESeq2.

## Strain construction

The *dcr-1* intestinal rescue strain was made by amplifying 1602 bases upstream of the *vha-6* translational start site, and subsequently using PCR to fuse the *vha-6* promoter to the *dcr-1* cDNA and 3’UTR. Purified PCR products were injected at 10 ng/μL with 1 ng/μL *myo-2p::mCherry* into *dcr-1(mg375)* to generate strain CQ636.

## *E. coli* expressing PA14 small RNA strain construction

For cloning the six differentially expressed PA14 sRNA species into *E. coli*, we utilized the experimentally determined sequences reported by Wurtzel et al. 2012 (Chambers and Sauer, 2013). sRNA sequences were amplified from *P. aeruginosa* PA14 genomic DNA using the primer pairs described below. The sRNAs were cloned into plasmid, pBAD18-Amp (Guzman et al., 1995) which contains an arabinose inducible promoter upstream of a multiple cloning site.

Plasmids were transformed into *E. coli* MG1655. sRNA expression was induced by growing *E. coli* on NGM plates supplemented with 0.1% arabinose. Proper sRNA production was confirmed for each of the six overexpression strains using RT-PCR.

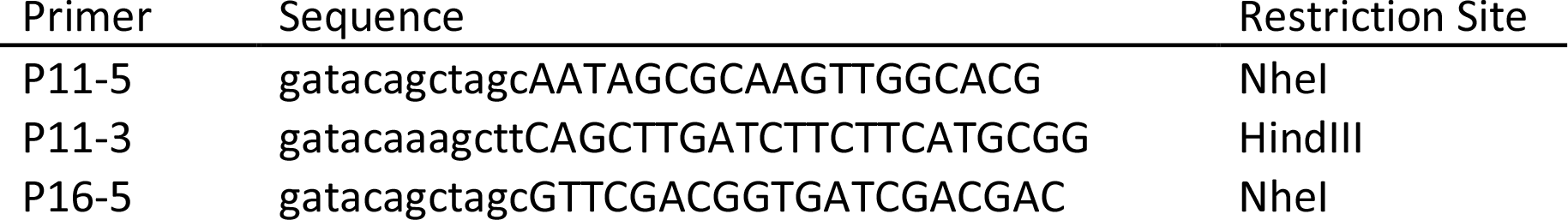
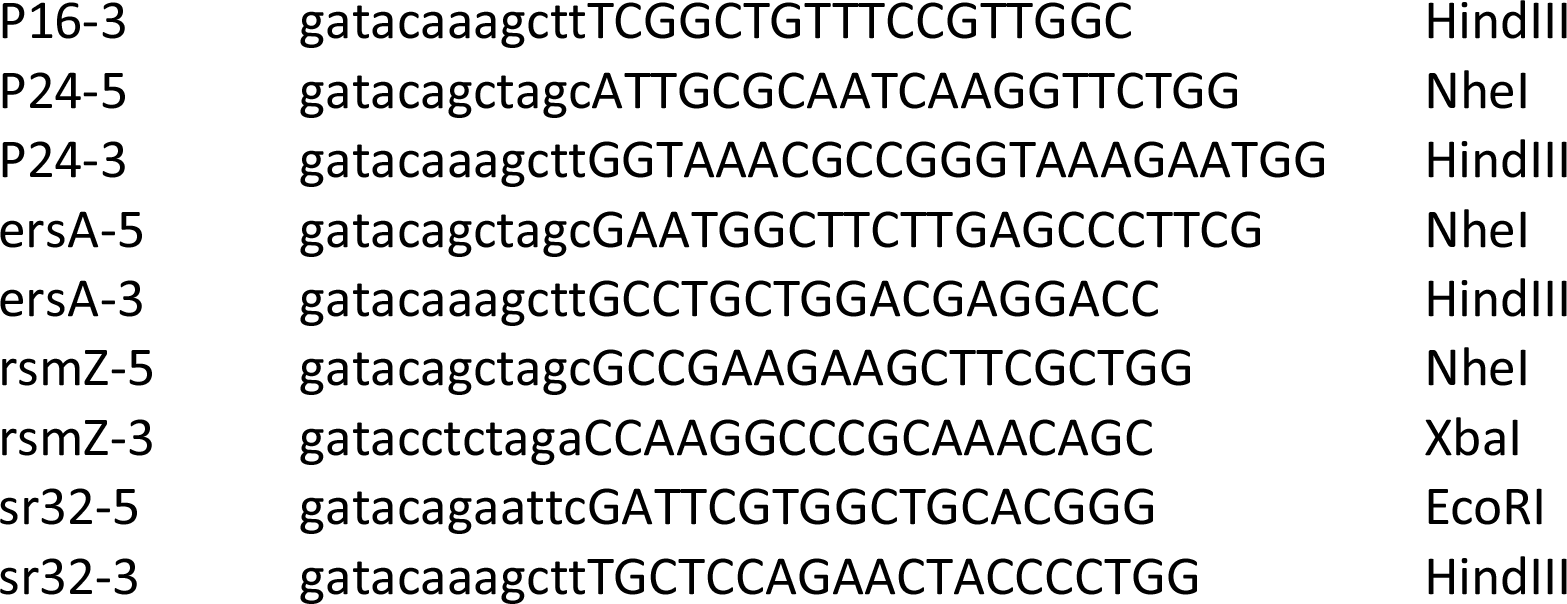

## PA14-ΔP11 mutant strain construction

The unmarked deletion of the P11 ncRNA was constructed by two-step allelic exchange using plasmid pEXG2 (Hmelo et al., 2015). Briefly, ∼400 bp fragments directly upstream and downstream of the P11 sequence were amplified from gDNA using primer pairs (P11-KO-1, P11-KO-2) and (P11-KO-3, P11-KO-4), respectively. Upstream and downstream fragments were fused together using overlap-extension PCR with primer pair (P11-KO-1, P11-KO-4), and the resulting fragment was cloned into the HindIII site of plasmid pEXG2. The pEXG2 plasmid was integrated into *P. aeruginosa* PA14 through conjugation with the donor strain *E. coli* S17.

Exconjugants were selected on 30 µg/mL gentamycin and then the mutants of interest were counterselected on 5% sucrose. Proper deletion was confirmed through PCR using primers (P11-seq-5, P11-seq-5).

P11-KO-1) GATACAAAGCTTCCTATGGCGAGATCGAAGCC

P11-KO-2) GCTCCTGCATAAGGTTGGCGCTTGCAGGGATAACGCGGG

P11-KO-3) CCCGCGTTATCCCTGCAAGCGCCAACCTTATGCAGGAGC

P11-KO-4) GATACAAAGCTTGCCGATTGCCGGGTATCG

P11-seq-5) GAAAGACCGCGGGGAGCC

P11-seq-3) CAGTTCTTCCAGGGTACGGACC

## *E. coli* expressing P11 *maco-1* homology mutant

To generate the P11 overexpression construct with mutations disrupting the 17-nt of homology to the *maco-I* gene, fragments to the left and right of the homology site were amplified from the pBAD18-P11 using primer pairs (P11-macoI-1, P11-macoI-2) and (P11-macoI-3, P11-macoI-4), respectively, and inserted into the NheI/HindIII-cut pBAD18 plasmid by Gibson assembly.

P11-maco-I-1 CTCCATACCCGTTTTTTTGGGCTAGCCATGGAGATAGGCAATAGCGC

P11-maco-I-2 CGCCGCCATTTTTCTGTGGCGGCGCTTTGTTGTCGATTGTCGGG

P11-maco-I-3 CGCCGCCACAGAAAAATGGCGGCGTTTCCCGACCGAACGGGAC

P11-maco-I-4 TCTCATCCGCCAAAACAGCCAAGCTCAGCTTGATCTTCTTCATGCGG

## PA14-ΔP11 and GRb0427 survival assay

OP50, PA14, PA14-ΔP11 GRb0427 were grown in liquid culture and diluted as described above. 200 μL of diluted bacteria was spread to completely cover a 6 cm NGM plate. Plates were incubated for 2 days at 25°C to allow bacterial growth. Plates were equilibrated to 20°C before the addition of L4 worms to plates. Assays were performed at 25°C. Assays were counted every 8-10 h until all animals on pathogenic plates died.

## Statistical analysis of choice assay data

For all learning assays, each dot represents an individual choice assay plate (∼10-300 worms/plate) with all data shown from at least 3 independent replicates (Table S4). Plates were excluded that contained less than 10 total worms per plate. The box extends from the 25^th^ to 75^th^ percentiles, with whiskers from the minimum to the maximum values. Mean differences are shown using Cumming estimation plots (Ho et al., 2019), with each graphed as a bootstrap sampling distribution. Mean differences are depicted as dots; 95% confidence intervals are indicated by the ends of the vertical bars. All figures in the main text and supplement represent pooled data from independent experiments. Results from individual experiments are provided in the statistics supplement. All estimation plots were generated using https://www.estimationstats.com/<u>#/</u>. Additional statistics were generated using Prism 8.

## Data and materials availability

Sequencing data is available: BioProject PRJNA553700.

## Acknowledgments

We thank Buck Samuel and the *C. elegans* Genetics Center for strains; the Genomics Core Facility at Princeton University; W. Wang for helping develop methods to sequence bacterial small RNAs; the Murphy lab for discussion; and Robert Clausen and Jasmine Ashraf for assistance. CTM is the Director of the Glenn Center for Aging Research at Princeton and an HHMI-Simons Faculty Scholar.

## Author contributions

RK, RSM, GDV, ZG, and CTM designed experiments. RK and RSM performed experiments and analyzed data. GDV constructed P11 *E. coli* and PA14 mutant strains. LP and RK analyzed small RNAseq data. RK, RSM, and CTM wrote the manuscript.

## Funding

This work was supported by a Pioneer Award to CTM (NIGMS DP1GM119167), the Glenn Foundation for Medical Research (GMFR CNV1001899), the HHMI-Simons Faculty Scholar Program (AWD1005048), T32GM007388 (NIGMS) support of RSM and GDV, and a Pioneer Award to ZG (DP1A1124669).

**Supplementary Information** is available for this paper.

Correspondence and requests for materials should be addressed to.

## Supplemental data figure/table legends

**Supplemental Figure 1:**
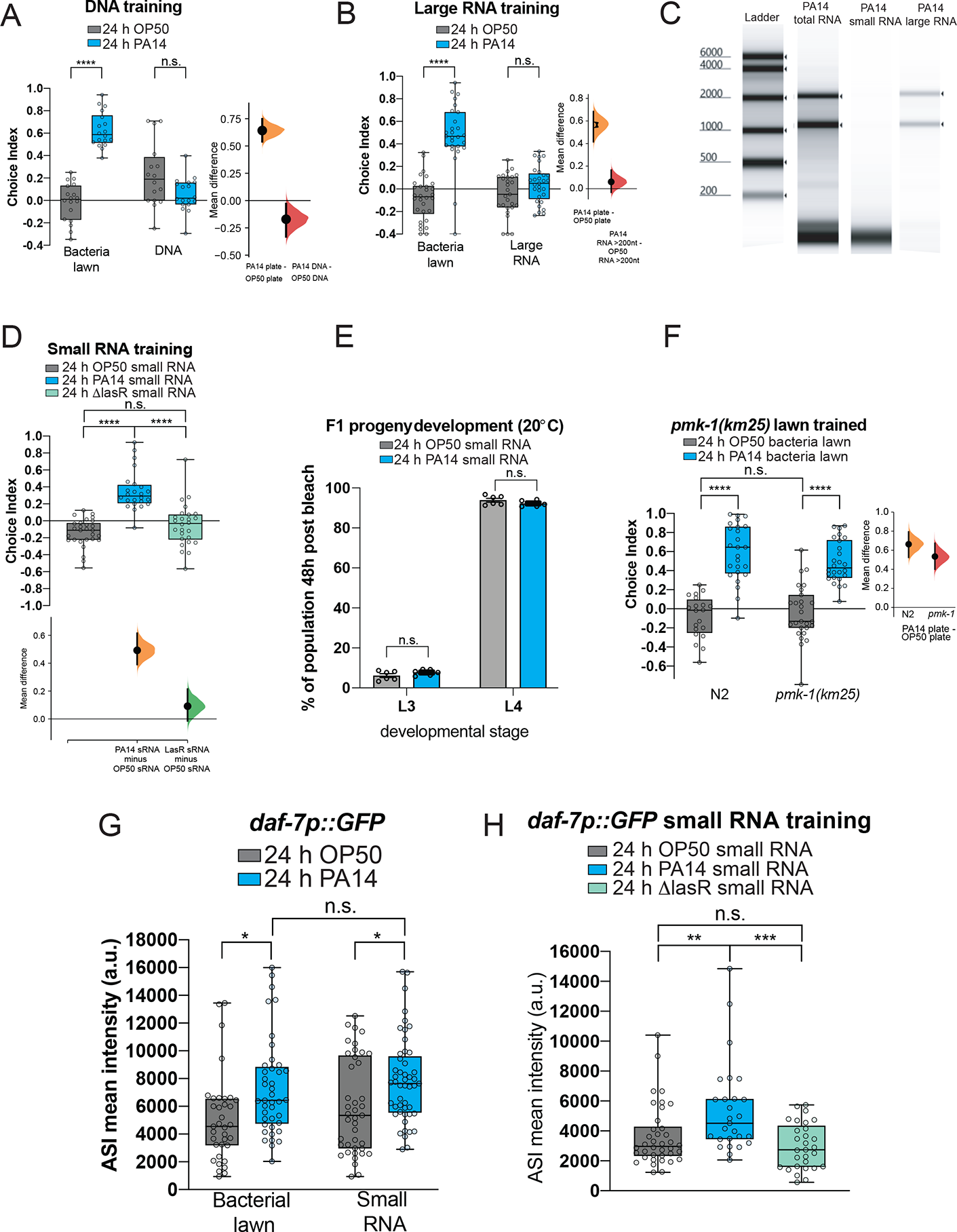
PA14 small RNAs induce maternal PA14 avoidance and increased *daf-7 expression in the ASI neurons*. (A) Worms exposed to a PA14 bacterial lawn for 24 hr learn to avoid PA14 in a choice assay, while PA14 DNA exposure alone does not induce maternal avoidance of PA14. (B) Training with large RNAs (>200 nt) isolated from bacterial lawns of PA14 is not sufficient for maternal PA14 avoidance. (C) Bioanalyzer results of isolated PA14 total RNA and fractionated small and large RNAs. RNA levels were normalized for worm training. (D) *ΔlasR* sRNA exposure does not induce PA14 learned avoidance. (E) *pmk-1(km25)* worms learn to avoid PA14 when exposed to PA14 bacteria lawns. (F) Development of progeny of PA14 small RNA-trained mothers was not delayed compared to progeny of OP50 trained mothers. n= 23-142 progeny per condition per plate. (D) One-Way or (A, B, E, F) Two-Way ANOVA, Tukey’s multiple comparison test. (A) Two or (B, D, F) three biological replicates were performed. ****p < 0.0001, n.s. = not significant. For all learning assays, each dot represents an individual choice assay plate (average 115 worms per plate) with all data shown from the indicated number of independent replicates. The box extends from the 25^th^ to 75^th^ percentiles, with whiskers from the minimum to the maximum values. Mean differences are shown using Cumming estimation plots (Ho et al., 2019), with each graphed as a bootstrap sampling distribution. Mean differences are depicted as dots; 95% confidence intervals are indicated by the ends of the vertical bars. (G) 24 h exposure of worms to a PA14 lawn or PA14 small RNAs increases *daf-7p::GFP* expression in the ASI neurons. Two-Way ANOVA, Tukey’s multiple comparison test. n ≥ 54 neurons. (H) *Δ*lasR sRNA exposure does not induces changes in *daf-7p::GFP* ASI expression. One-Way ANOVA, Tukey’s multiple comparison test. n ≥ 27 neurons. (G-H) Three biological replicates were performed. *p ≤ 0.05, **p ≤ 0.01, ***p ≤ 0.001, ****p ≤ 0.0001, n.s. = not significant.

**Supplemental Figure 2:**
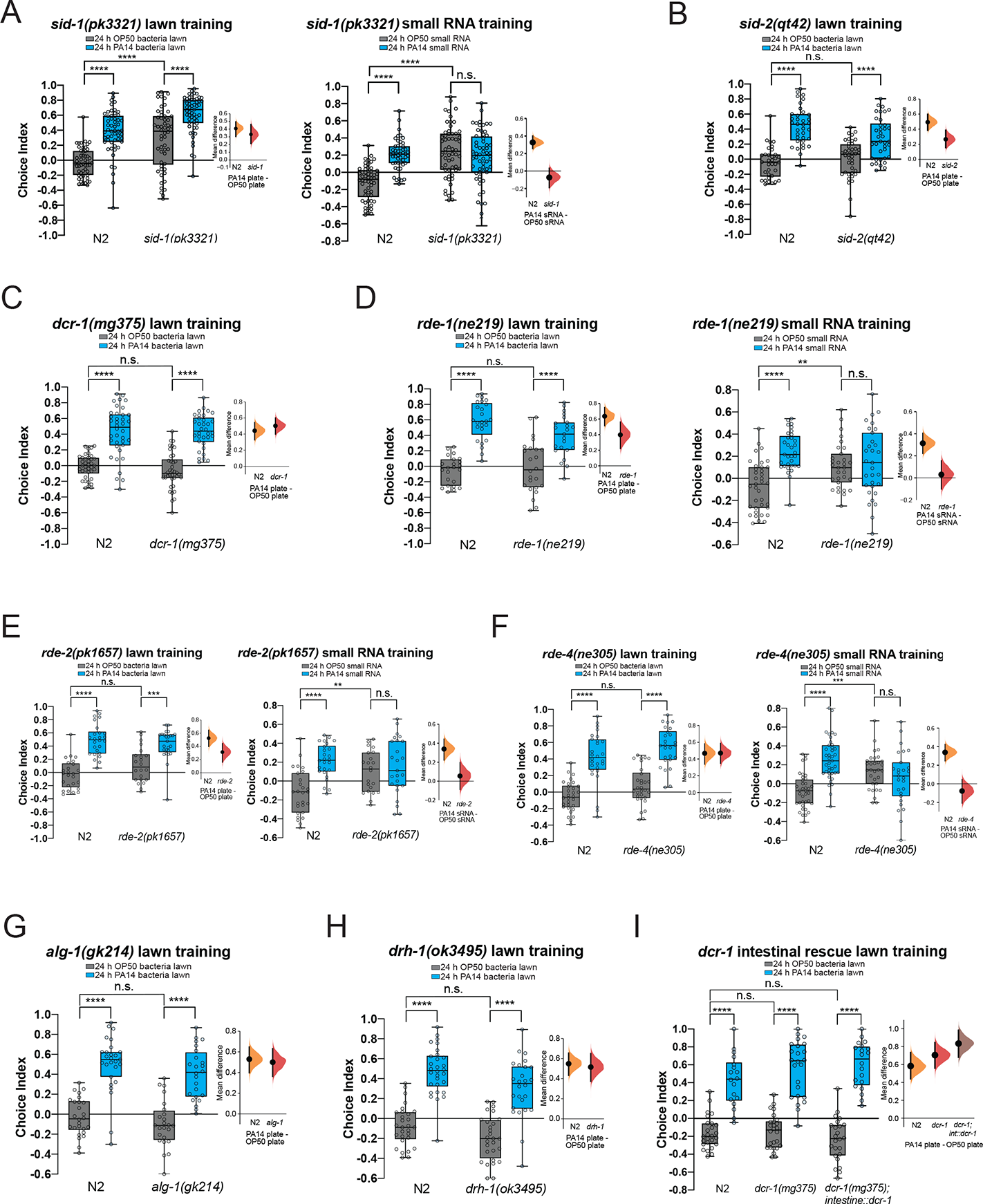
Small RNA and bacterial lawn training of *C. elegans* RNAi pathway mutants. (A) Wild-type and *sid-1(pk3321)* worms were trained on OP50 or PA14 bacterial lawns (left) or small RNA (right) for 24h and tested for learned PA14 avoidance. *sid-1* mutants have a constitutively high naïve preference, but are able to learn avoidance after training on a PA14 lawn, but not after training on small RNAs. (B-C) Wild-type and *sid-2(qt42)* (B) or *dcr-1(mg375)* (C) worms were trained on OP50 or PA14 bacterial lawns for 24h and tested for learned PA14 avoidance. (D-F) Wild-type, *rde-1(ne219)* (A), *rde-2(pk1657)* (B), or *rde-4(ne305)* (C) worms were trained on OP50 or PA14 bacterial lawns or small RNA for 24h and tested for learned PA14 avoidance. *rde-1, rde-2,* and *rde-4* mutants are able to learn avoidance following bacteria lawn training, but do not avoid PA14 following small RNA training. These mutants have high naïve preference after training on small RNAs only. (G-H) Wild-type and *alg-1(gk214)* (G) or *drh-1(ok3495)* (H) worms were trained on OP50 or PA14 bacterial lawns for 24h and tested for learned PA14 avoidance. (I) Wild-type, *dcr-1(mg375)*, and *dcr-1(mg375);vha-6p::dcr-1* worms were trained on OP50 or PA14 bacterial lawns for 24h and tested for learned PA14 avoidance. Training and choice assays for *dcr-1(mg375)* and *dcr-1(mg375);vha-6p::dcr-1* worms were performed on the same plates. For choice assays, transgenic worms expressing the rescue construct and pharyngeal mCherry were counted using a fluorescence dissecting microscope. Non-transgenic, non-fluorescent *dcr-1(mg375)* siblings from the same plates were also counted and the results are shown. For all learning assays, each dot represents an individual choice assay plate (average of 115 worms per plate) with all data shown from 3 independent replicates. The box extends from the 25^th^ to 75^th^ percentiles, with whiskers from the minimum to the maximum values. Mean differences are shown using Cumming estimation plots (Ho et al., 2019), with each graphed as a bootstrap sampling distribution. Mean differences are depicted as dots; 95% confidence intervals are indicated by the ends of the vertical bars. Two-Way ANOVA, Tukey’s multiple comparison test. **p ≤ 0.01, ***p ≤ 0.001, ****p < 0.0001, n.s. = not significant.

**Supplemental Figure 3:**
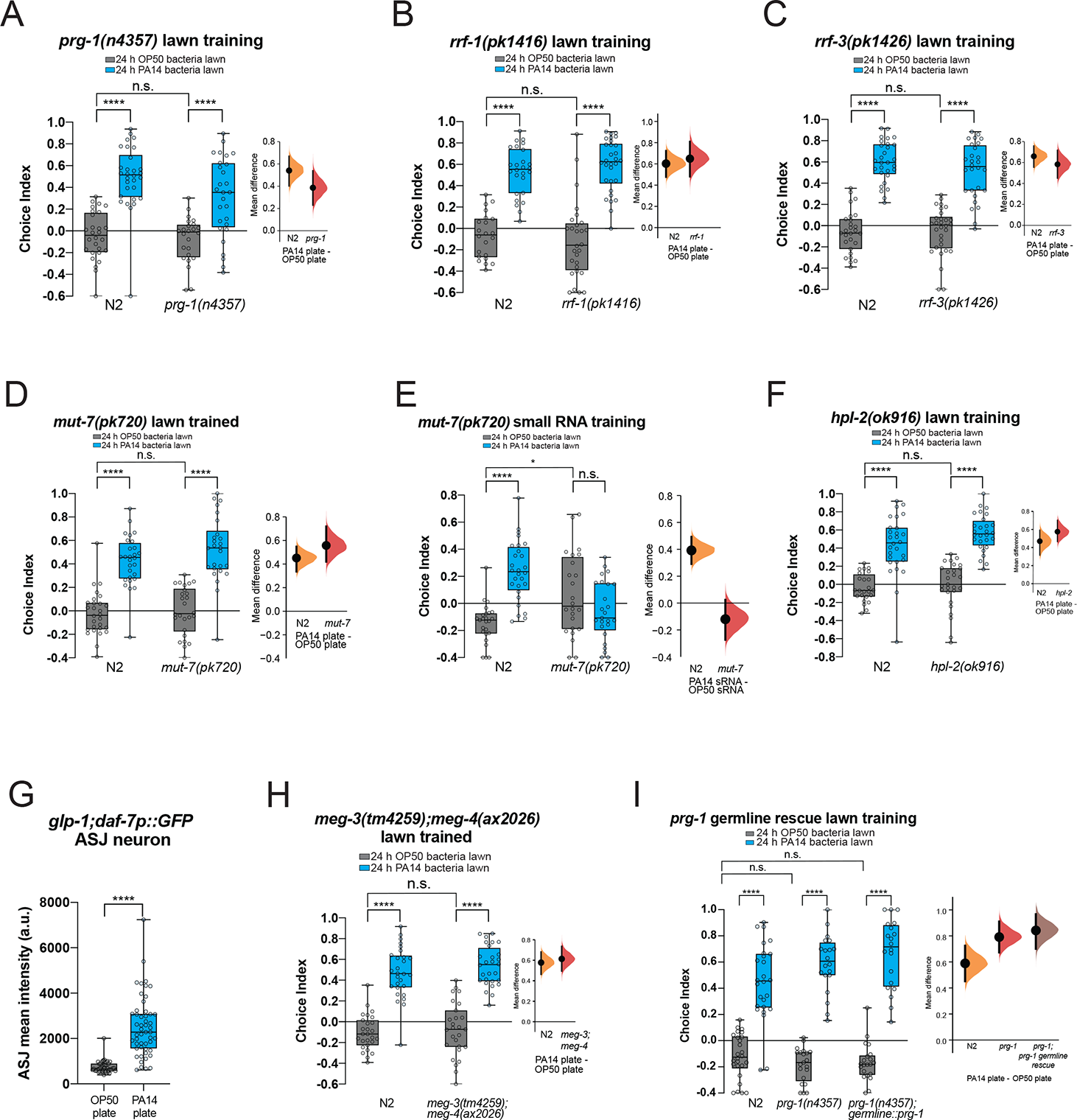
Small RNA and bacterial lawn training of *C. elegans* small RNA pathway and germline mutants. (A-D) Wild-type and *prg-1(n4357)* (A), *rrf-1(pk1416)* (B), *rrf-3(pk1426)* (C), or *mut-7(pk720)* (D) worms were trained on OP50 or PA14 bacterial lawns for 24h and tested for learned PA14 avoidance. (E) Wild-type and *mut-7(pk720)* worms were trained on OP50 or PA14 sRNA for 24h and tested for learned PA14 avoidance. (F) Wild-type and *hpl-2(ok916),* worms were trained on OP50 or PA14 bacterial lawns for 24h and tested for learned PA14 avoidance. (G) Germline-less *glp-1* mutants induce *daf-7p::GFP* expression in the ASJ neuron after PA14 lawn exposure. Student’s t-test. ****p < 0.0001. (H) Wild-type and *meg-3(tm4259);meg-4(ax2026)* worms were trained on OP50 or PA14 bacterial lawns for 24h and tested for learned PA14 avoidance. (I) Wild-type, *prg-1(n4357),* and *prg-1(n4357);pie-1p::GFP::prg-1* worms were trained on OP50 or PA14 bacterial lawns for 24h and tested for learned PA14 avoidance. Each dot represents an individual choice assay plate (average of 115 worms per plate) with all data shown from 3 independent replicates. The box extends from the 25^th^ to 75^th^ percentiles, with whiskers from the minimum to the maximum values. Mean differences are shown using Cumming estimation plots (Ho et al., 2019), with each graphed as a bootstrap sampling distribution. Mean differences are depicted as dots; 95% confidence intervals are indicated by the ends of the vertical bars. Two-Way ANOVA, Tukey’s multiple comparison test. *p ≤ 0.05, ****p < 0.0001, n.s. = not significant.

**Supplemental Figure 4:**
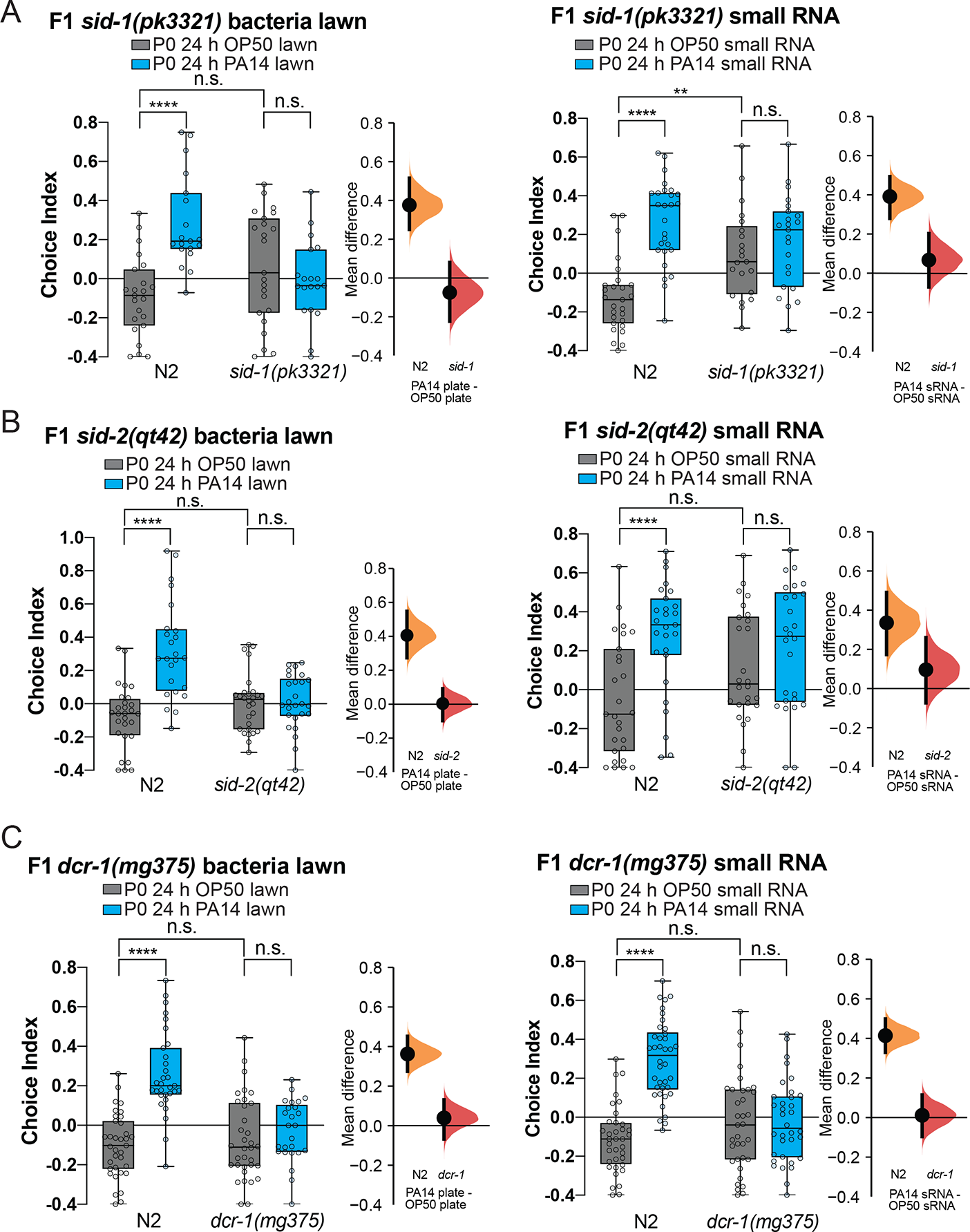
*sid-1, sid-2,* and *dcr-1* F1 progeny of plate and small RNA trained parents. (A-C) Progeny of (A) *sid-1,* (B) *sid-2,* and (C) *dcr-1* are defective in both transgenerational pathogen avoidance following maternal bacterial lawn (left) and PA14 sRNA training (right). (A-C) were independently repeated at least 3 times. Each dot represents an individual choice assay plate (average of 115 worms per plate) with all data shown from 3 independent replicates. The box extends from the 25^th^ to 75^th^ percentiles, with whiskers from the minimum to the maximum values. Mean differences are shown using Cumming estimation plots (Ho et al., 2019), with each graphed as a bootstrap sampling distribution. Mean differences are depicted as dots; 95% confidence intervals are indicated by the ends of the vertical bars. Two-Way ANOVA, Tukey’s multiple comparison test. **p ≤ 0.01, ****p < 0.0001, n.s. = not significant.

**Supplemental Figure 5:**
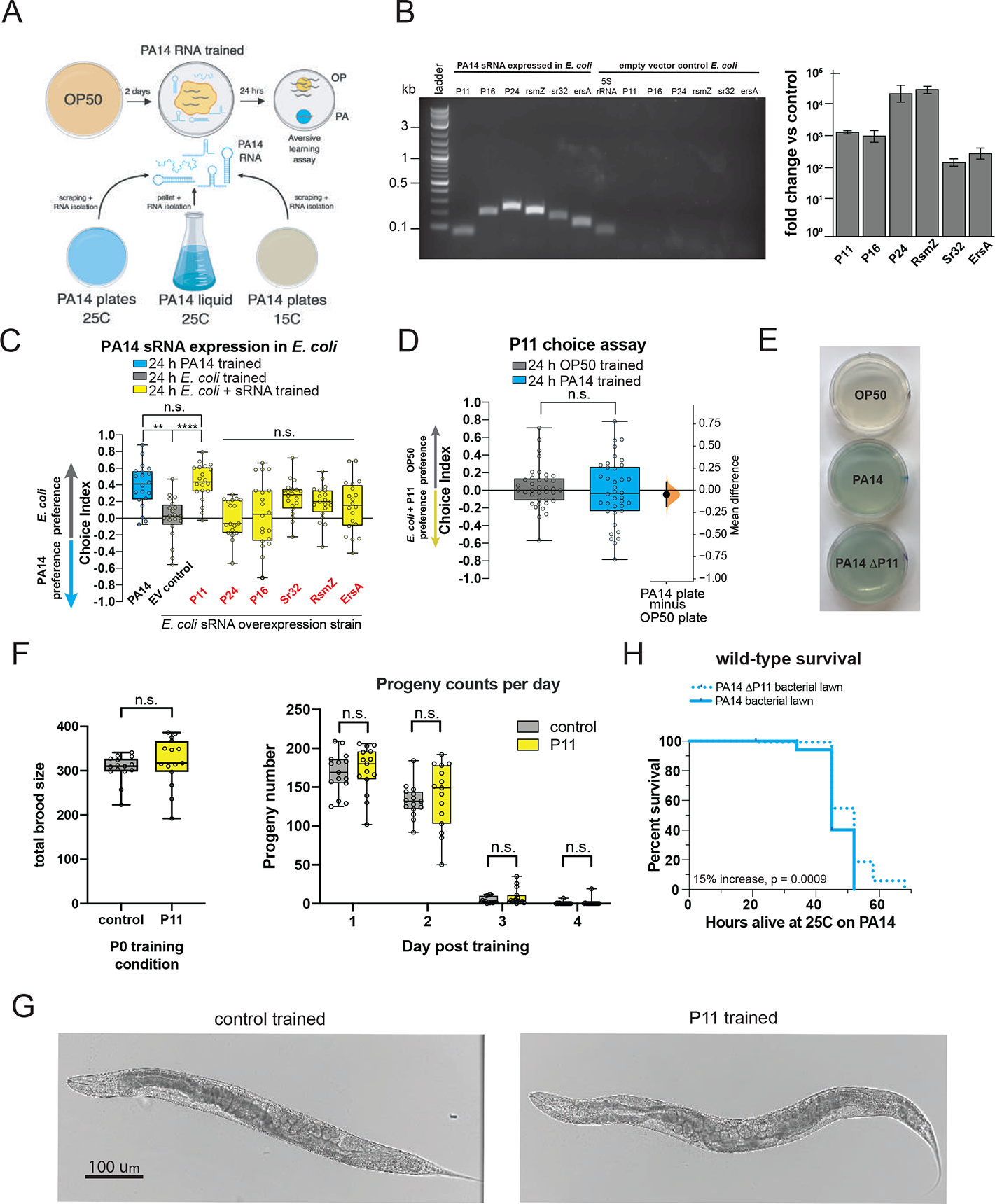
Identification and testing of the differentially regulated PA14 small RNAs (A) Small RNA training protocol using RNA isolated from PA14 cultures grown at 25°C or 15°C on plates, or in liquid culture. (B, left) Each candidate PA14 small RNA was expressed in *E. coli* strain MG1655 using an arabinose-inducible promoter. Arabinose-induced expression of each sRNA was confirmed using qRT-PCR using sRNA-specific primers (first 6 lanes). Amplification of 5S rRNA from the empty vector control strain is shown. No PCR products were observed (last 6 lanes) using the sRNA-specific primer sets with the empty vector control. (B, right) Quantification of sRNA levels in *E. coli* MG1655 strains by RT-qPCR. Expression level is the ΔCt between the sRNA producing strain and a vector control converted to fold change using the amplification factor. Values are mean +/-SD normalized by levels of a 5S rRNA control. (C) P11 expressed in *E. coli* induces PA14 avoidance in a choice assay between PA14 and *E. coli* strain MG1655. Worms were trained on OP50, PA14, *E. coli* strain MG1655 containing empty vector (EV control), or *E. coli* strain MG1655 expressing PA14 small RNAs (red). Following training, a choice assay was performed between *E. coli* strain MG1655 and PA14. Similar to Fig 4F, worms trained on PA14 or *E. coli* expressing P11 exhibited PA14 avoidance behavior. (D) PA14 plate-trained worms do not avoid *E. coli* expressing P11 compared to OP50 in a choice assay. Each dot represents an individual choice assay plate (average of 115 worms per plate) with all data shown from 2 independent replicates. The box extends from the 25^th^ to 75^th^ percentiles, with whiskers from the minimum to the maximum values. Mean differences are shown using Cumming estimation plots (Ho et al., 2019), with each graphed as a bootstrap sampling distribution. Mean differences are depicted as dots; 95% confidence intervals are indicated by the ends of the vertical bars. One-Way ANOVA, Tukey’s multiple comparison test. **p ≤ 0.01, ****p < 0.0001, n.s = not significant. (E) PA14-ΔP11 bacteria grown on NGM plates produce pyocyanin (blue pigment on plates) at levels similar to PA14. (F) Worms exposed to empty vector (top) or *E. coli* expressing P11 appear healthy after 24 of training. (G) Exposure to *E. coli* expressing P11 does not affect total brood size (left) (Student’s t-test), or the number of progeny hatched per day(right) (C, Two-Way ANOVA, Tukey’s multiple comparison test). n.s. = not significant. 15 worms were analyzed per condition. (H) PA14-ΔP11 is less pathogenic compared to PA14. Log-rank (Mantel-Cox test), n=120 animals/condition.

**Supplemental Figure 6:**
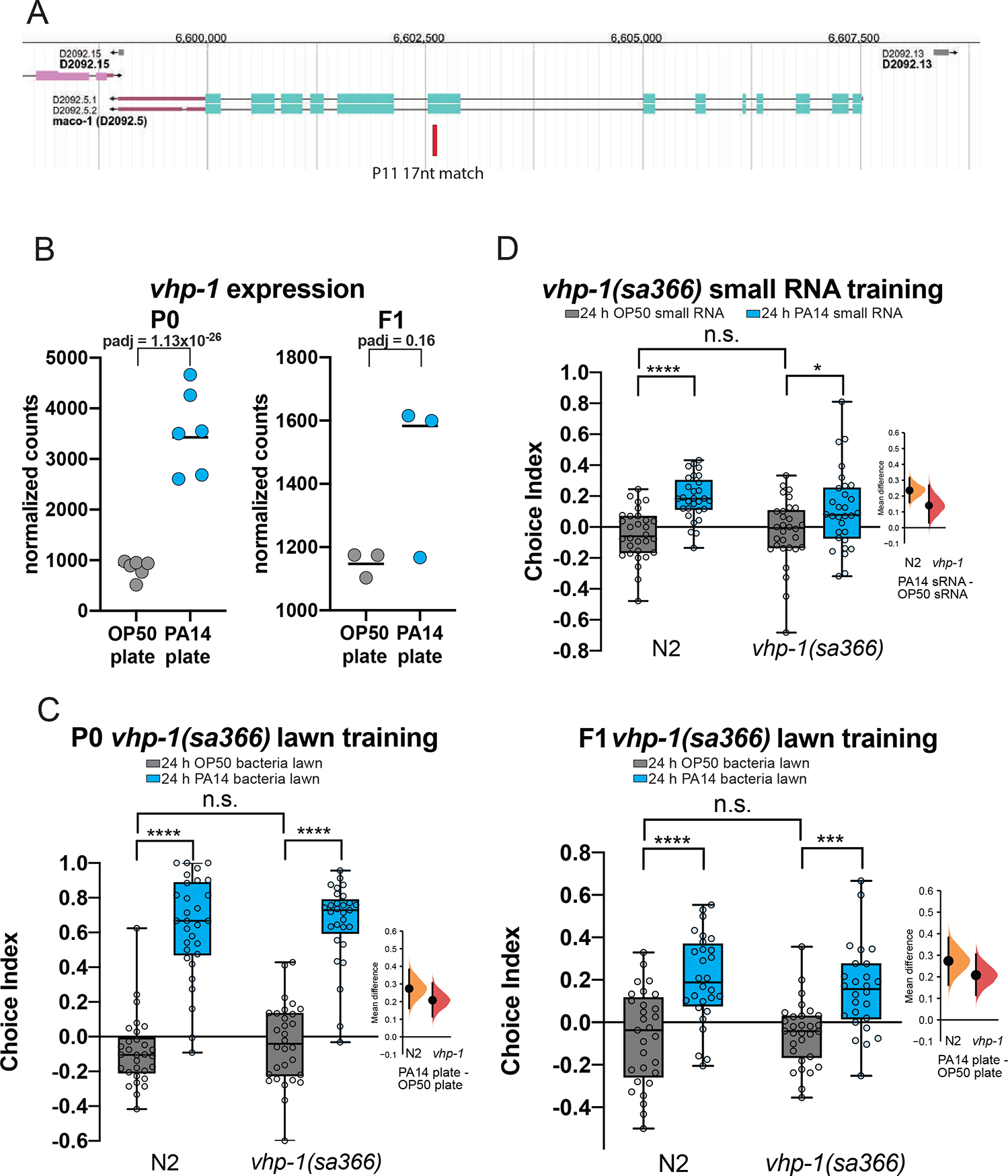
P11 region of homology with *maco-1* and behavior of *vhp-1* mutants. (A) Wormbase depiction of the *maco-1* genomic locus and the region of P11 homology (red). (B) Wild-type, and *vhp-1(sa366)* worms were trained on OP50 or PA14 bacterial lawns for 24h and tested for learned PA14 avoidance (top). Progeny of trained animals were tested for learned PA14 avoidance (bottom). (C) Wild-type, and *vhp-1(sa366)* worms were trained on OP50 or PA14 small RNA for 24h and tested for learned PA14 avoidance. Each dot represents an individual choice assay plate (average of 115 worms per plate) with all data shown from 3 independent replicates. The box extends from the 25^th^ to 75^th^ percentiles, with whiskers from the minimum to the maximum values. Mean differences are shown using Cumming estimation plots (Ho et al., 2019), with each graphed as a bootstrap sampling distribution. Mean differences are depicted as dots; 95% confidence intervals are indicated by the ends of the vertical bars. Two-Way ANOVA, Tukey’s multiple comparison test. *p ≤ 0.05, ***p ≤ 0.001, ****p < 0.0001, n.s. = not significant.

**Supplemental Table 1: DESeq2 results comparing annotated small RNAs isolated from PA14 bacteria grown at 25°C or 15°C on plates.**

**Supplemental Table 2: DESeq2 results comparing annotated small RNAs isolated from PA14 bacteria grown at 25°C on plated or in liquid.**

**Supplemental Table 3: P11 homology to C elegans coding and noncoding RNAs**

**Supplemental File 4: Estimation statistics for pooled and individual choice assay data.**

