## Supplemental File 4 Cumming Estimation Statistics for "*C. elegans* “reads” bacterial non-coding RNAs to learn pathogenic avoidance"

# 1A

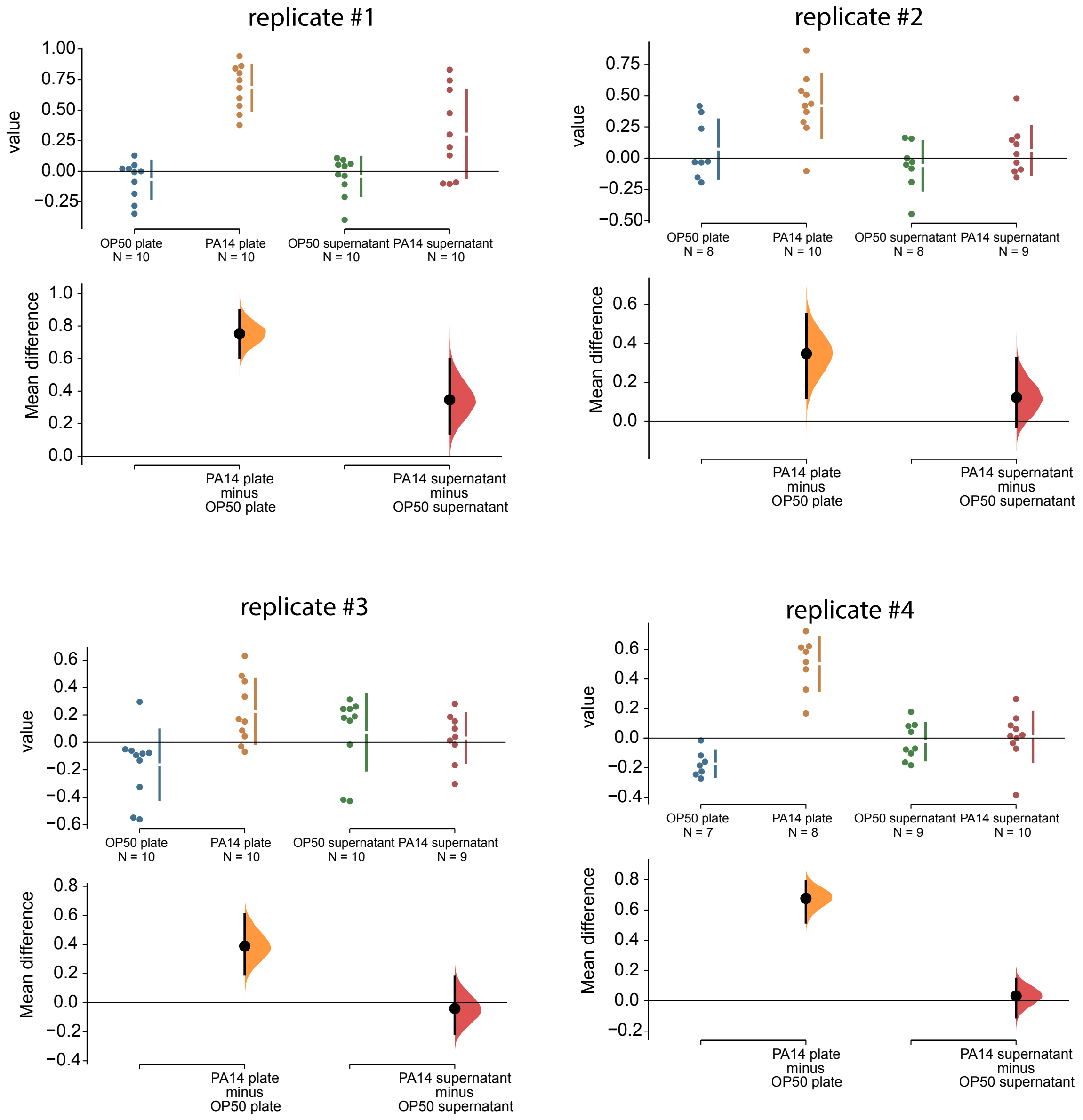
replicate #1
replicate #2
replicate #3
replicate #4
value
Mean difference
OP50 plate
N = 10
PA14 plate
N = 10
OP50 supernatant
N = 10
PA14 supernatant
N = 10
PA14 plate
minus
OP50 plate
PA14 supernatant
minus
OP50 supernatant
OP50 plate
N = 8
PA14 plate
N = 10
OP50 supernatant
N = 8
PA14 supernatant
N = 9
OP50 plate
N = 10
PA14 plate
N = 10
OP50 supernatant
N = 10
PA14 supernatant
N = 9
OP50 plate
N = 7
PA14 plate
N = 8
OP50 supernatant
N = 9
PA14 supernatant
N = 10

# 1C

replicate #1

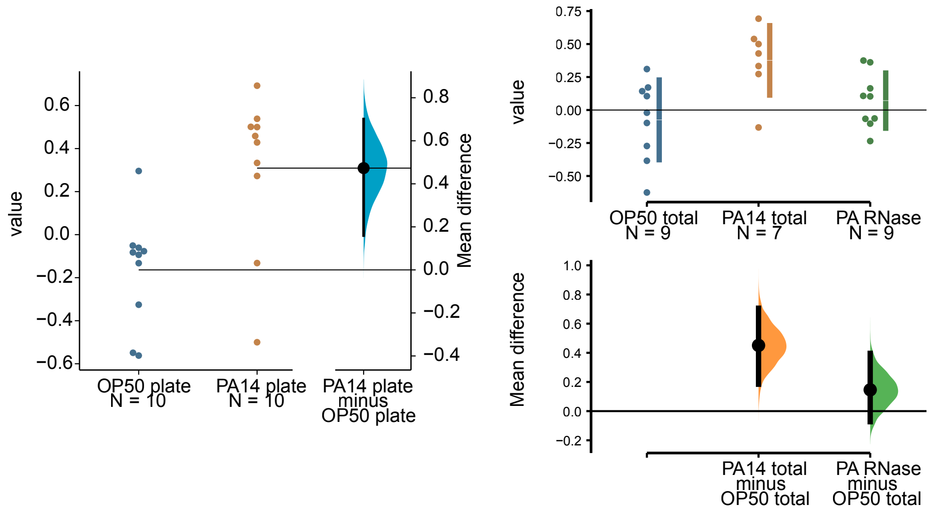


replicate #2

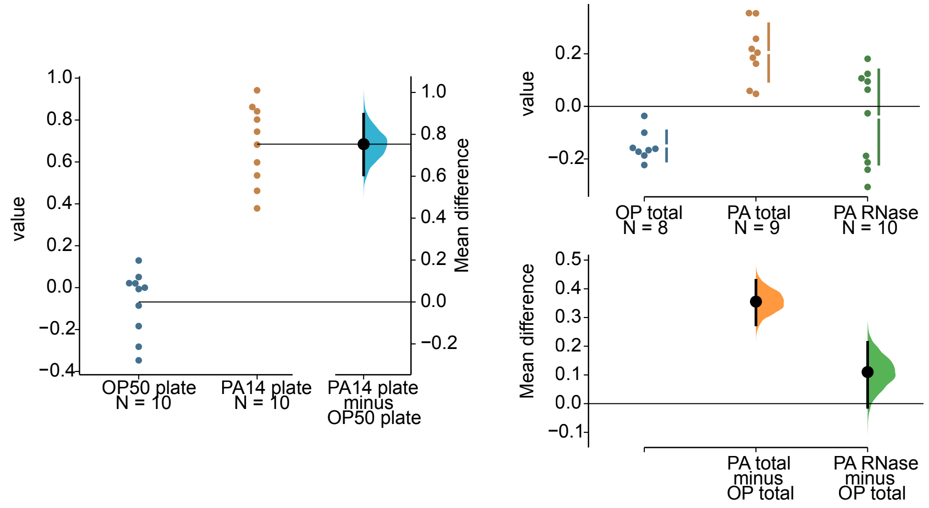


replicate #3

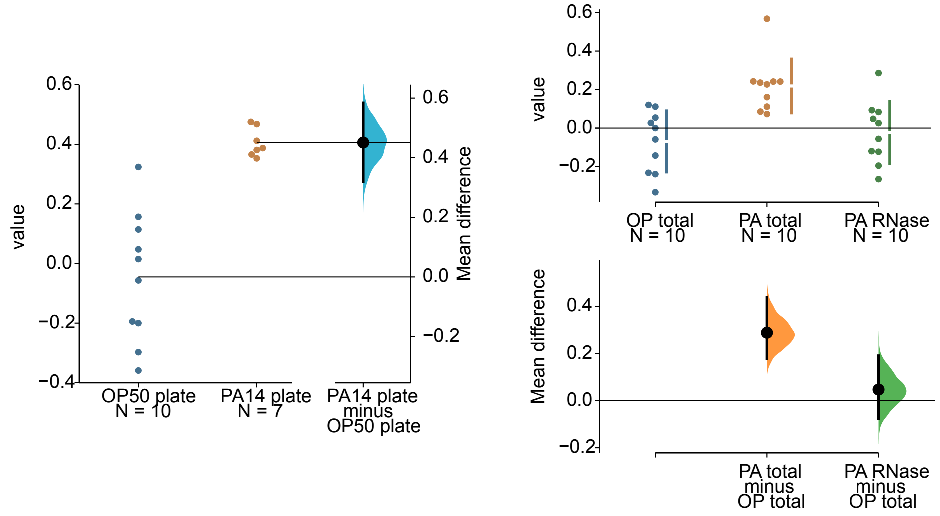

1D

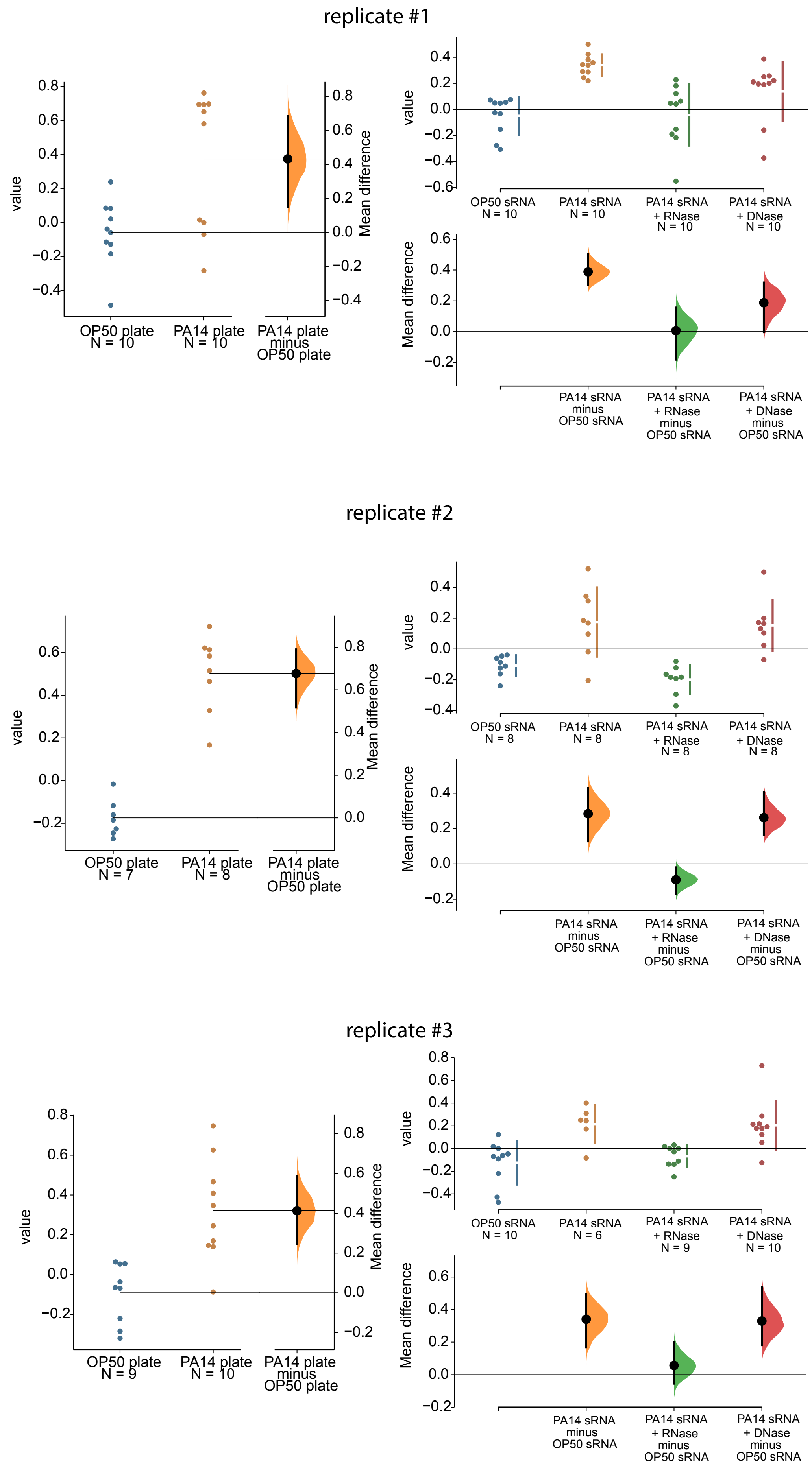

replicate #1
value
Mean difference
OP50 plate
N = 10
PA14 plate
N = 10
PA14 plate
minus
OP50 plate
OP50 sRNA
N = 10
PA14 sRNA
N = 10
PA14 sRNA
+ RNase
N = 10
PA14 sRNA
+ DNase
N = 10
PA14 sRNA
minus
OP50 sRNA
PA14 sRNA
+ RNase
minus
OP50 sRNA
PA14 sRNA
+ DNase
minus
OP50 sRNA
replicate #2
OP50 plate
N = 7
PA14 plate
N = 8
OP50 sRNA
N = 8
PA14 sRNA
N = 8
PA14 sRNA
+ RNase
N = 8
PA14 sRNA
+ DNase
N = 8
replicate #3
OP50 plate
N = 9
PA14 plate
N = 10
OP50 sRNA
N = 10
PA14 sRNA
N = 6
PA14 sRNA
+ RNase
N = 9
PA14 sRNA
+ DNase
N = 10

1E

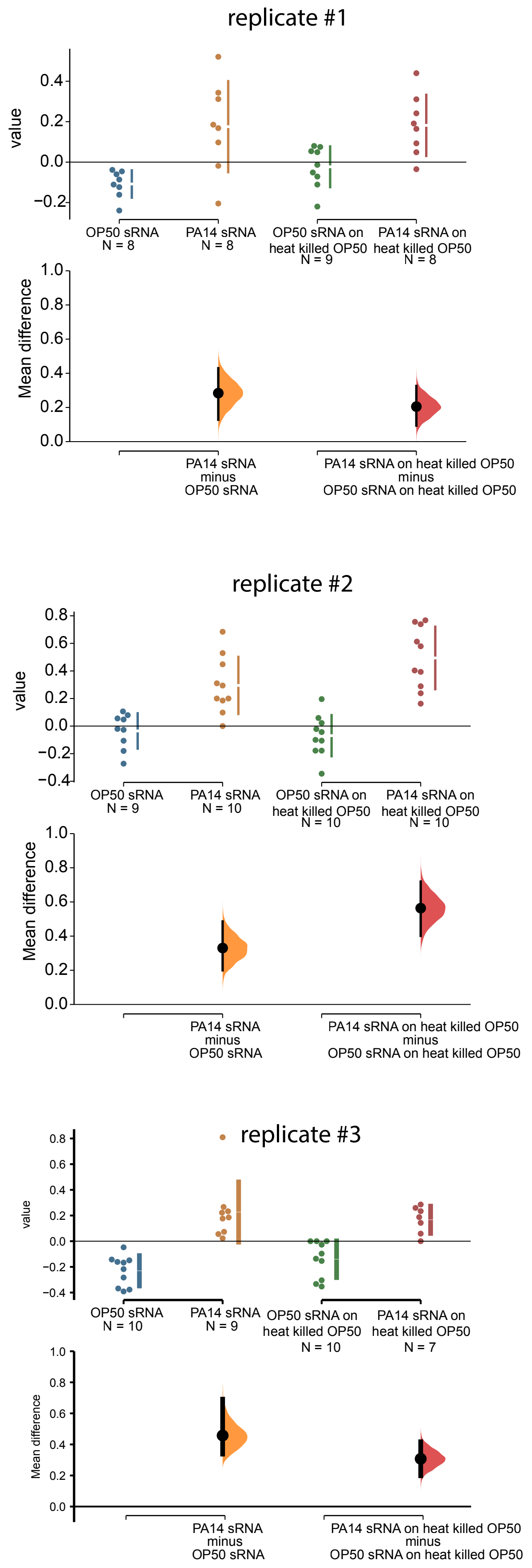
replicate #1
value
0.4
0.2
0.0
−0.2
OP50 sRNA
N = 8
PA14 sRNA
N = 8
OP50 sRNA on
heat killed OP50
N = 9
PA14 sRNA on
heat killed OP50
N = 8
Mean difference
1.0
0.8
0.6
0.4
0.2
0.0
PA14 sRNA
minus
OP50 sRNA
PA14 sRNA on heat killed OP50
minus
OP50 sRNA on heat killed OP50
replicate #2
value
0.8
0.6
0.4
0.2
0.0
−0.2
−0.4
OP50 sRNA
N = 9
PA14 sRNA
N = 10
OP50 sRNA on
heat killed OP50
N = 10
PA14 sRNA on
heat killed OP50
N = 10
Mean difference
1.0
0.8
0.6
0.4
0.2
0.0
PA14 sRNA
minus
OP50 sRNA
PA14 sRNA on heat killed OP50
minus
OP50 sRNA on heat killed OP50
replicate #3
value
0.8
0.6
0.4
0.2
0.0
−0.2
−0.4
OP50 sRNA
N = 10
PA14 sRNA
N = 9
OP50 sRNA on
heat killed OP50
N = 10
PA14 sRNA on
heat killed OP50
N = 7
Mean difference
1.0
0.8
0.6
0.4
0.2
0.0
PA14 sRNA
minus
OP50 sRNA
PA14 sRNA on heat killed OP50
minus
OP50 sRNA on heat killed OP50

1G

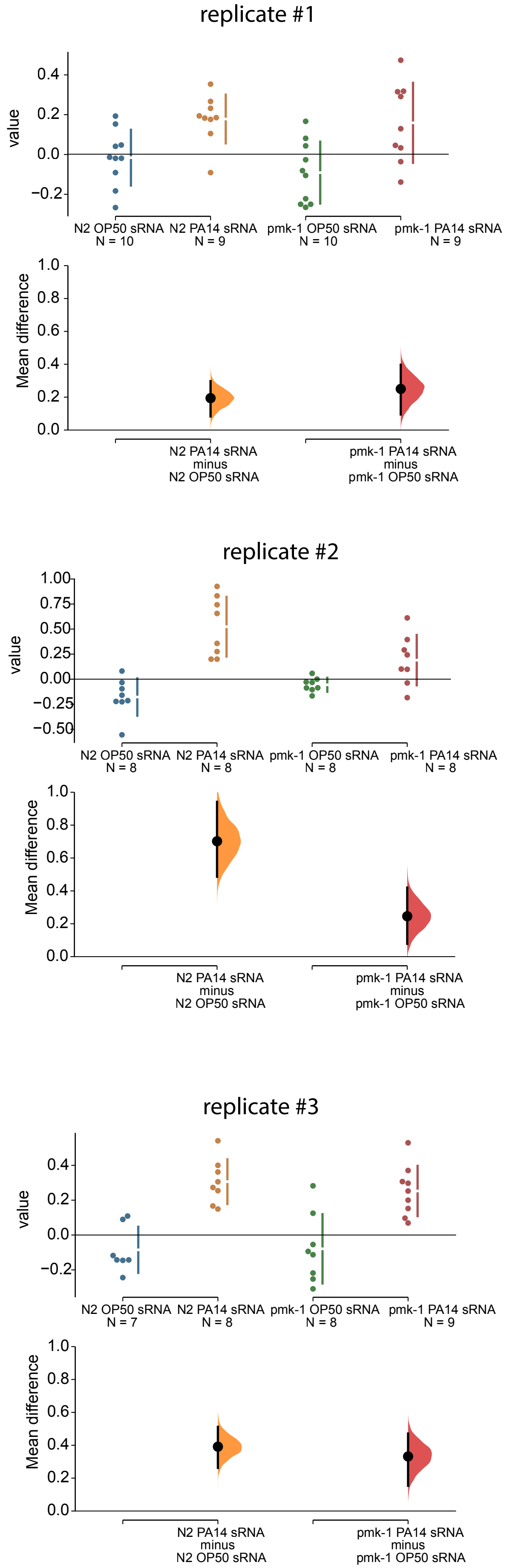

replicate #1
value
0.4
0.2
0.0
−0.2
N2 OP50 sRNA
N = 10
N2 PA14 sRNA
N = 9
pmk-1 OP50 sRNA
N = 10
pmk-1 PA14 sRNA
N = 9
Mean difference
1.0
0.8
0.6
0.4
0.2
0.0
N2 PA14 sRNA
minus
N2 OP50 sRNA
pmk-1 PA14 sRNA
minus
pmk-1 OP50 sRNA
replicate #2
value
1.00
0.75
0.50
0.25
0.00
−0.25
−0.50
N2 OP50 sRNA
N = 8
N2 PA14 sRNA
N = 8
pmk-1 OP50 sRNA
N = 8
pmk-1 PA14 sRNA
N = 8
Mean difference
1.0
0.8
0.6
0.4
0.2
0.0
N2 PA14 sRNA
minus
N2 OP50 sRNA
pmk-1 PA14 sRNA
minus
pmk-1 OP50 sRNA
replicate #3
value
0.4
0.2
0.0
−0.2
N2 OP50 sRNA
N = 7
N2 PA14 sRNA
N = 8
pmk-1 OP50 sRNA
N = 8
pmk-1 PA14 sRNA
N = 9
Mean difference
1.0
0.8
0.6
0.4
0.2
0.0
N2 PA14 sRNA
minus
N2 OP50 sRNA
pmk-1 PA14 sRNA
minus
pmk-1 OP50 sRNA

# 1L

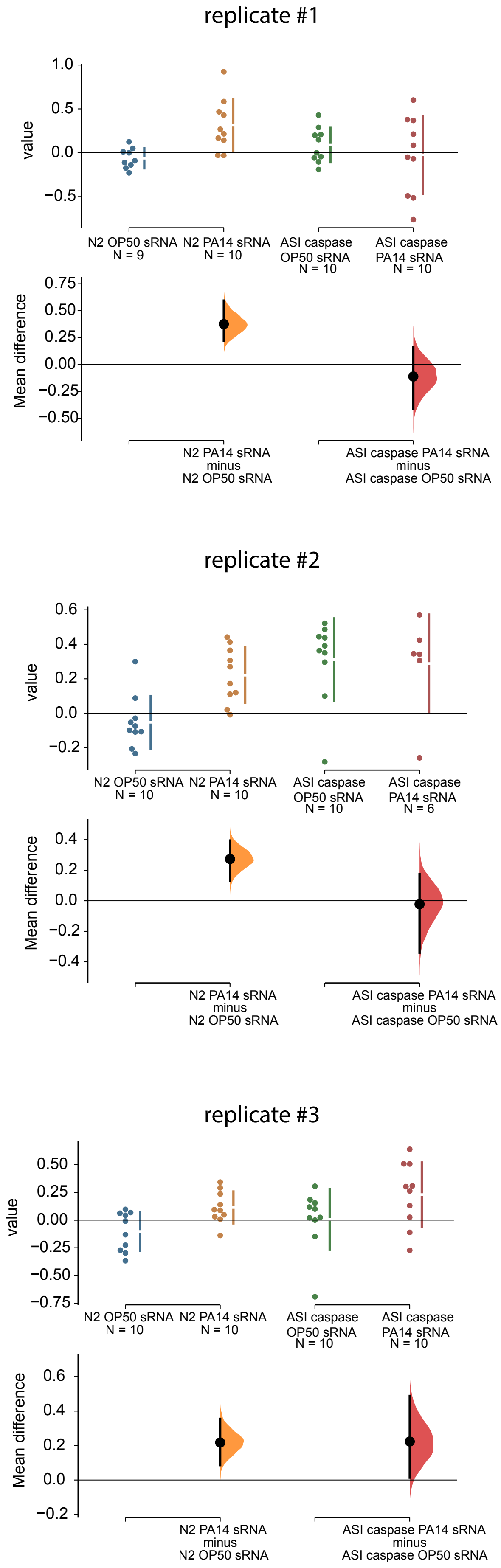
replicate #1
value
1.0
0.5
0.0
−0.5
N2 OP50 sRNA
N = 9
N2 PA14 sRNA
N = 10
ASI caspase
OP50 sRNA
N = 10
ASI caspase
PA14 sRNA
N = 10
Mean difference
0.75
0.50
0.25
0.00
−0.25
−0.50
N2 PA14 sRNA
minus
N2 OP50 sRNA
ASI caspase PA14 sRNA
minus
ASI caspase OP50 sRNA
replicate #2
value
0.6
0.4
0.2
0.0
−0.2
N2 OP50 sRNA
N = 10
N2 PA14 sRNA
N = 10
ASI caspase
OP50 sRNA
N = 10
ASI caspase
PA14 sRNA
N = 6
Mean difference
0.4
0.2
0.0
−0.2
−0.4
N2 PA14 sRNA
minus
N2 OP50 sRNA
ASI caspase PA14 sRNA
minus
ASI caspase OP50 sRNA
replicate #3
value
0.50
0.25
0.00
−0.25
−0.50
−0.75
N2 OP50 sRNA
N = 10
N2 PA14 sRNA
N = 10
ASI caspase
OP50 sRNA
N = 10
ASI caspase
PA14 sRNA
N = 10
Mean difference
0.6
0.4
0.2
0.0
−0.2
N2 PA14 sRNA
minus
N2 OP50 sRNA
ASI caspase PA14 sRNA
minus
ASI caspase OP50 sRNA

2A

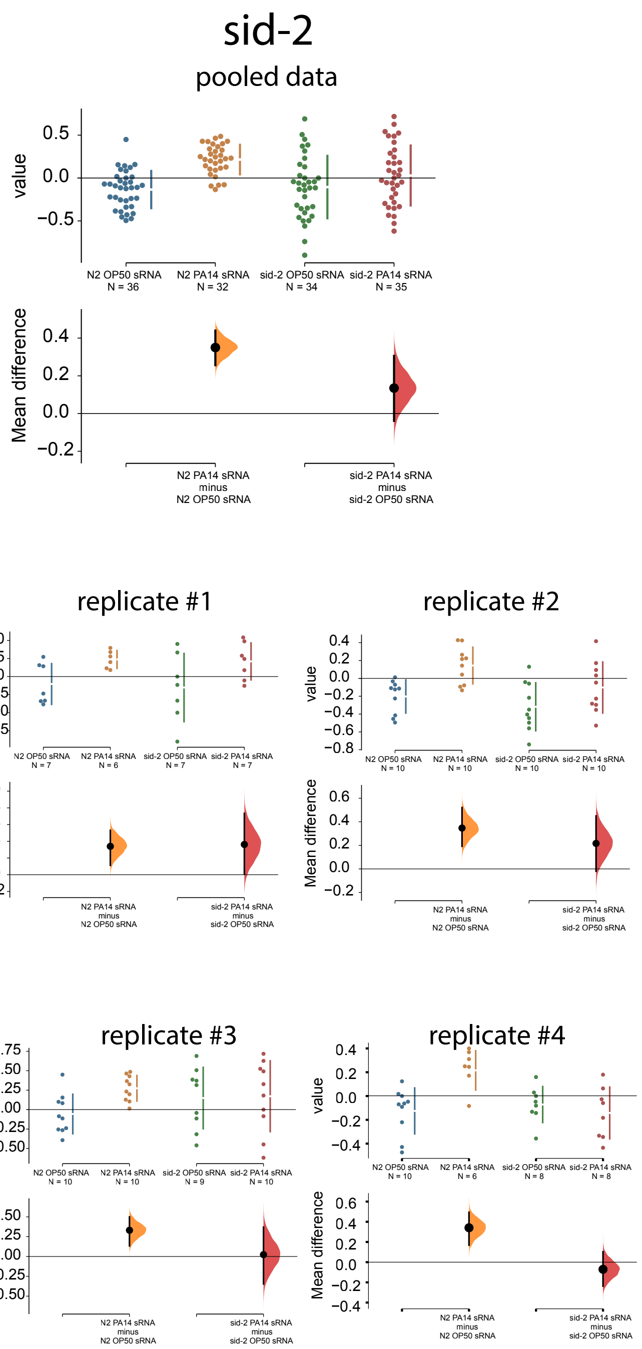
sid-2
pooled data
value
Mean difference
N2 OP50 sRNA
N = 36
N2 PA14 sRNA
N = 32
sid-2 OP50 sRNA
N = 34
sid-2 PA14 sRNA
N = 35
N2 PA14 sRNA
minus
N2 OP50 sRNA
sid-2 PA14 sRNA
minus
sid-2 OP50 sRNA
replicate #1
replicate #2
replicate #3
replicate #4

2B

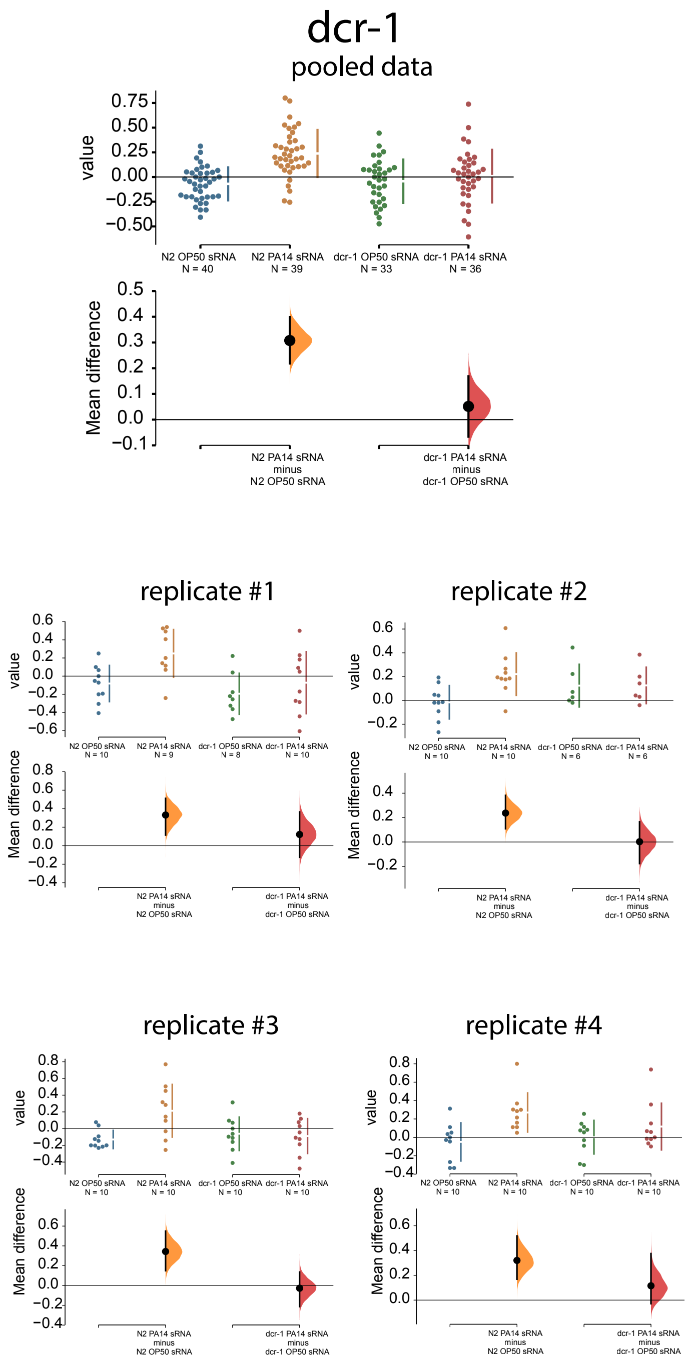
dcr-1
pooled data
value
N2 OP50 sRNA N = 40
N2 PA14 sRNA N = 39
dcr-1 OP50 sRNA N = 33
dcr-1 PA14 sRNA N = 36
Mean difference
N2 PA14 sRNA minus N2 OP50 sRNA
dcr-1 PA14 sRNA minus dcr-1 OP50 sRNA
replicate #1
replicate #2
replicate #3
replicate #4

2C

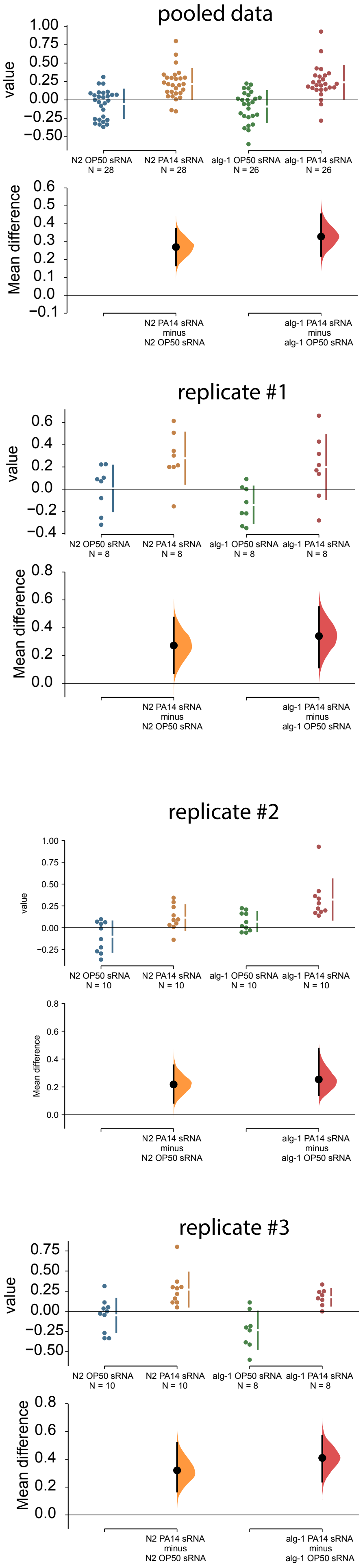

pooled data
value
1.00
0.75
0.50
0.25
0.00
−0.25
−0.50
N2 OP50 sRNA
N = 28
N2 PA14 sRNA
N = 28
alg-1 OP50 sRNA
N = 26
alg-1 PA14 sRNA
N = 26
Mean difference
0.6
0.5
0.4
0.3
0.2
0.1
0.0
−0.1
N2 PA14 sRNA
minus
N2 OP50 sRNA
alg-1 PA14 sRNA
minus
alg-1 OP50 sRNA
replicate #1
value
0.6
0.4
0.2
0.0
−0.2
−0.4
N2 OP50 sRNA
N = 8
N2 PA14 sRNA
N = 8
alg-1 OP50 sRNA
N = 8
alg-1 PA14 sRNA
N = 8
Mean difference
0.8
0.6
0.4
0.2
0.0
N2 PA14 sRNA
minus
N2 OP50 sRNA
alg-1 PA14 sRNA
minus
alg-1 OP50 sRNA
replicate #2
value
1.00
0.75
0.50
0.25
0.00
−0.25
N2 OP50 sRNA
N = 10
N2 PA14 sRNA
N = 10
alg-1 OP50 sRNA
N = 10
alg-1 PA14 sRNA
N = 10
Mean difference
0.8
0.6
0.4
0.2
0.0
N2 PA14 sRNA
minus
N2 OP50 sRNA
alg-1 PA14 sRNA
minus
alg-1 OP50 sRNA
replicate #3
value
0.75
0.50
0.25
0.00
−0.25
−0.50
N2 OP50 sRNA
N = 10
N2 PA14 sRNA
N = 10
alg-1 OP50 sRNA
N = 8
alg-1 PA14 sRNA
N = 8
Mean difference
0.8
0.6
0.4
0.2
0.0
N2 PA14 sRNA
minus
N2 OP50 sRNA
alg-1 PA14 sRNA
minus
alg-1 OP50 sRNA

2D

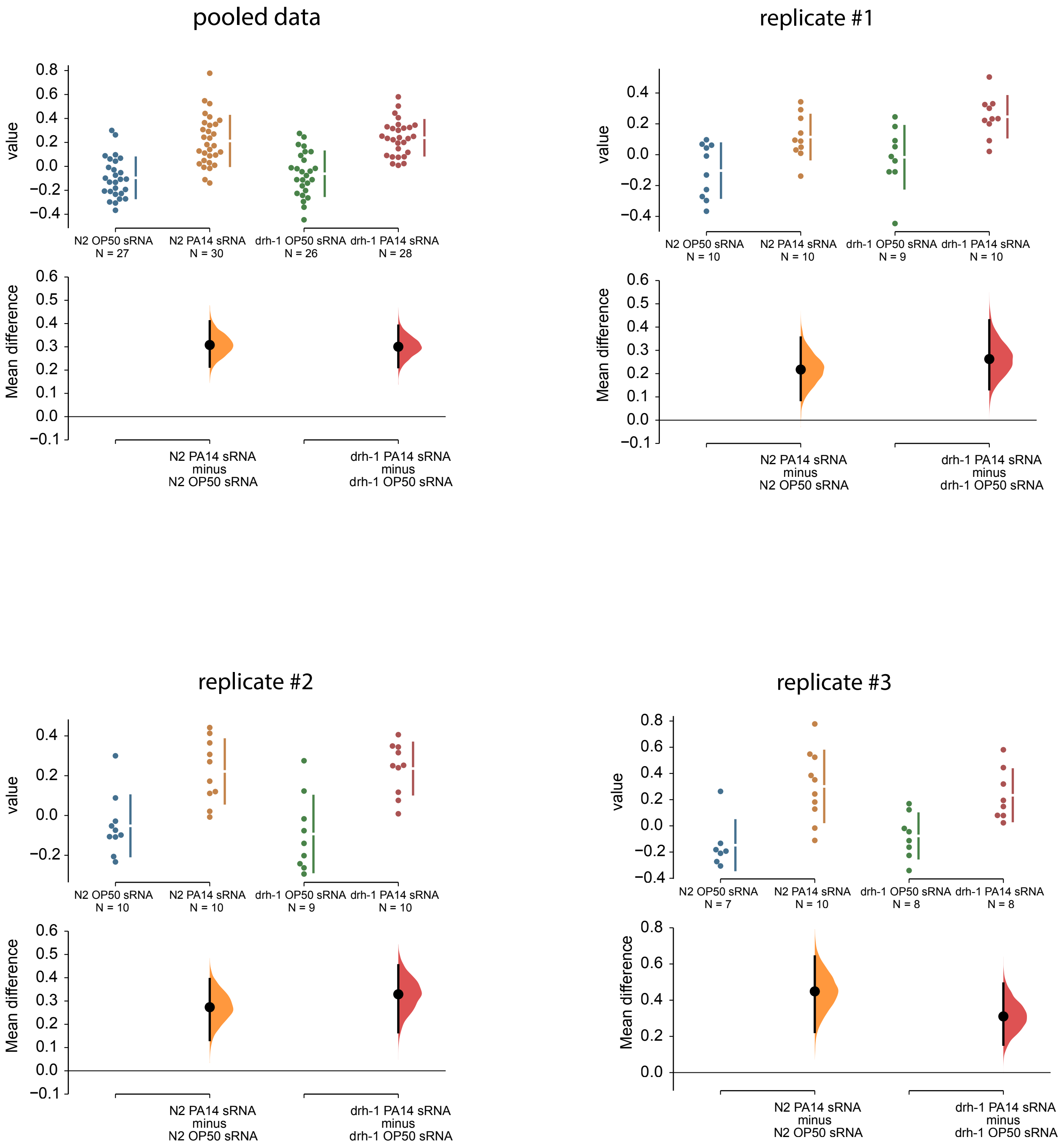
pooled data
replicate #1
replicate #2
replicate #3
value
Mean difference
N2 OP50 sRNA
N = 27
N2 PA14 sRNA
N = 30
drh-1 OP50 sRNA
N = 26
drh-1 PA14 sRNA
N = 28
N2 PA14 sRNA minus N2 OP50 sRNA
drh-1 PA14 sRNA minus drh-1 OP50 sRNA
N2 OP50 sRNA
N = 10
N2 PA14 sRNA
N = 10
drh-1 OP50 sRNA
N = 9
drh-1 PA14 sRNA
N = 10
N2 OP50 sRNA
N = 7
N2 PA14 sRNA
N = 10
drh-1 OP50 sRNA
N = 8
drh-1 PA14 sRNA
N = 8

2E

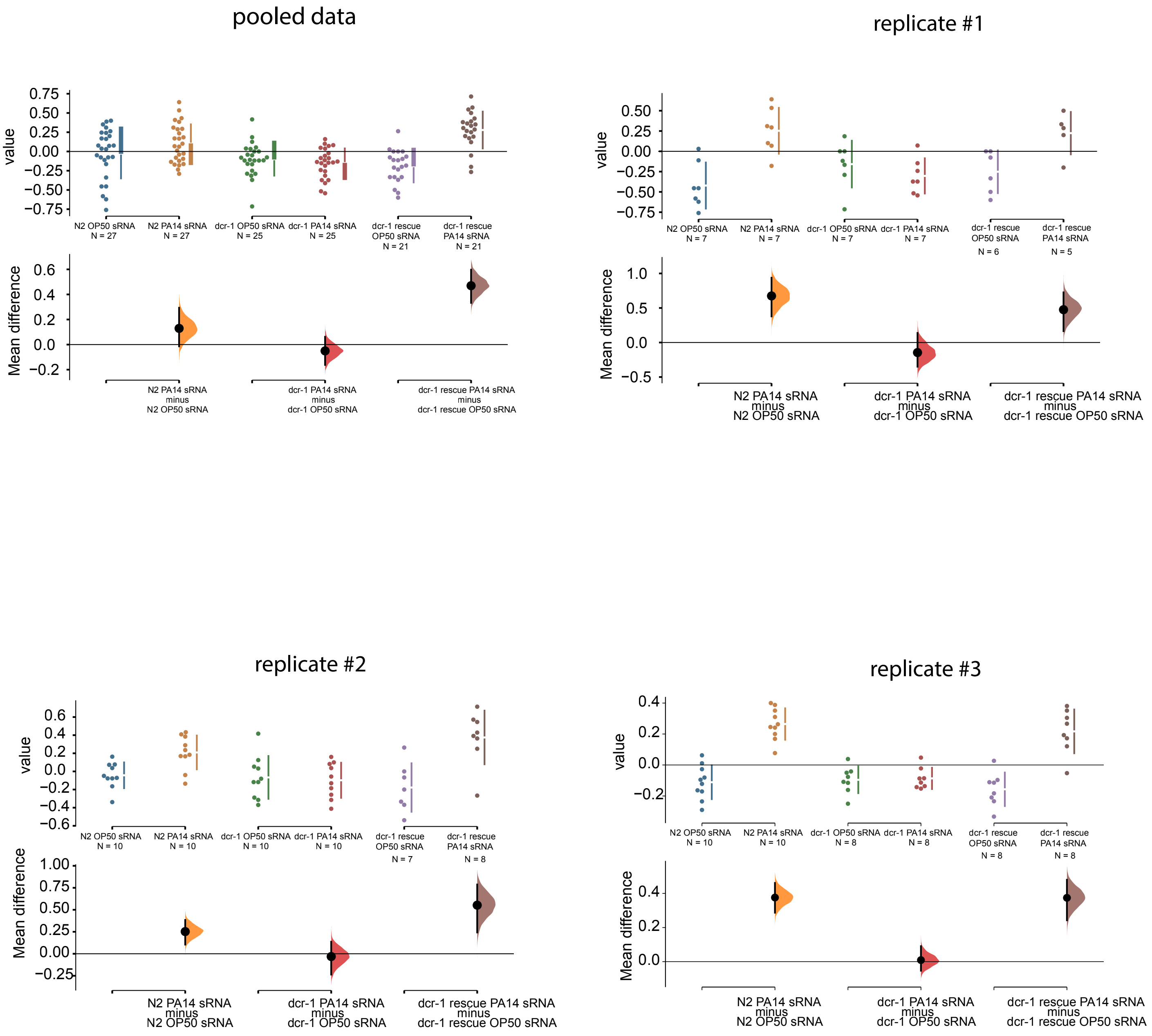
pooled data
value
Mean difference
N2 OP50 sRNA N = 27
N2 PA14 sRNA N = 27
dcr-1 OP50 sRNA N = 25
dcr-1 PA14 sRNA N = 25
dcr-1 rescue OP50 sRNA N = 21
dcr-1 rescue PA14 sRNA N = 21
N2 PA14 sRNA minus N2 OP50 sRNA
dcr-1 PA14 sRNA minus dcr-1 OP50 sRNA
dcr-1 rescue PA14 sRNA minus dcr-1 rescue OP50 sRNA
replicate #1
value
Mean difference
N2 OP50 sRNA N = 7
N2 PA14 sRNA N = 7
dcr-1 OP50 sRNA N = 7
dcr-1 PA14 sRNA N = 7
dcr-1 rescue OP50 sRNA N = 6
dcr-1 rescue PA14 sRNA N = 5
N2 PA14 sRNA minus N2 OP50 sRNA
dcr-1 PA14 sRNA minus dcr-1 OP50 sRNA
dcr-1 rescue PA14 sRNA minus dcr-1 rescue OP50 sRNA
replicate #2
value
Mean difference
N2 OP50 sRNA N = 10
N2 PA14 sRNA N = 10
dcr-1 OP50 sRNA N = 10
dcr-1 PA14 sRNA N = 10
dcr-1 rescue OP50 sRNA N = 7
dcr-1 rescue PA14 sRNA N = 8
N2 PA14 sRNA minus N2 OP50 sRNA
dcr-1 PA14 sRNA minus dcr-1 OP50 sRNA
dcr-1 rescue PA14 sRNA minus dcr-1 rescue OP50 sRNA
replicate #3
value
Mean difference
N2 OP50 sRNA N = 10
N2 PA14 sRNA N = 10
dcr-1 OP50 sRNA N = 8
dcr-1 PA14 sRNA N = 8
dcr-1 rescue OP50 sRNA N = 8
dcr-1 rescue PA14 sRNA N = 8
N2 PA14 sRNA minus N2 OP50 sRNA
dcr-1 PA14 sRNA minus dcr-1 OP50 sRNA
dcr-1 rescue PA14 sRNA minus dcr-1 rescue OP50 sRNA

2F

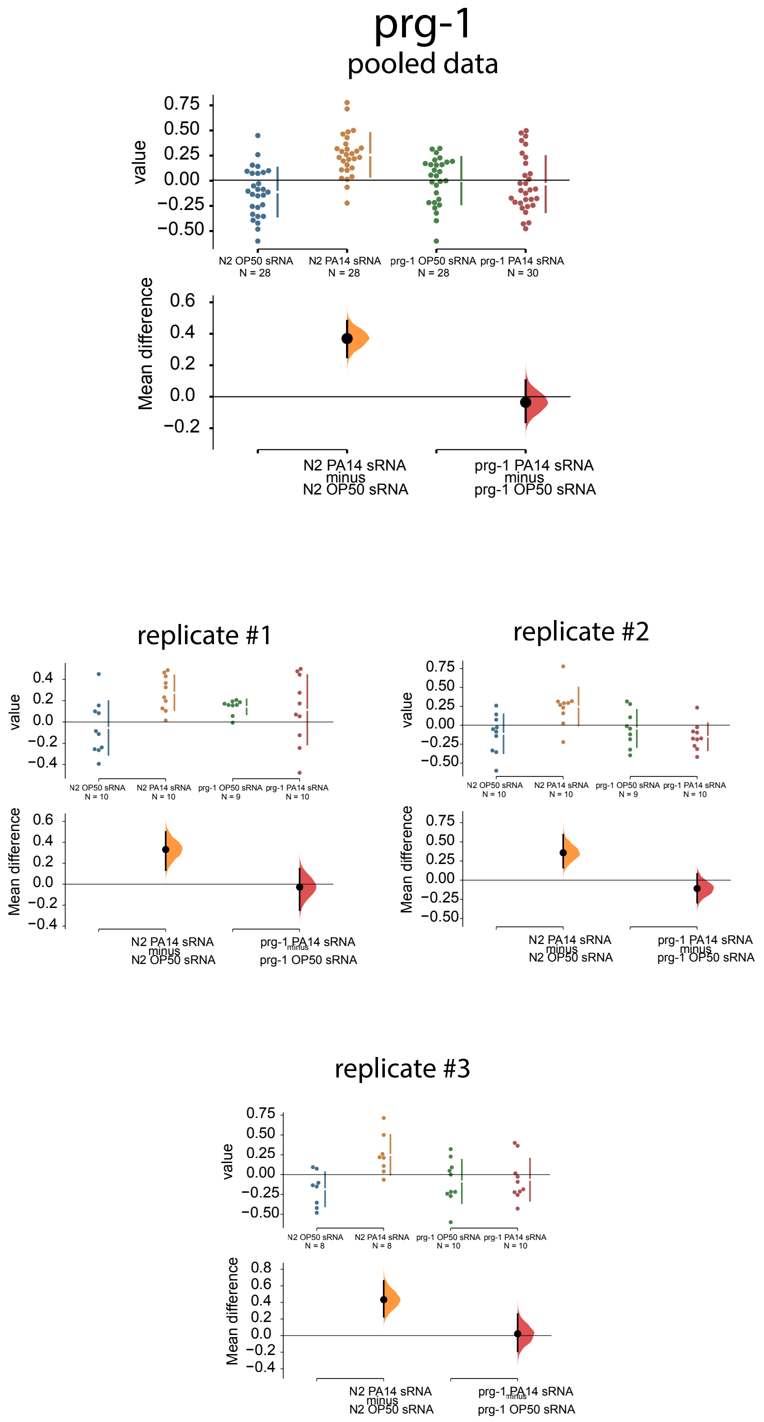
prg-1
pooled data
value
Mean difference
N2 OP50 sRNA
N = 28
N2 PA14 sRNA
N = 28
prg-1 OP50 sRNA
N = 28
prg-1 PA14 sRNA
N = 30
N2 PA14 sRNA
minus
N2 OP50 sRNA
prg-1 PA14 sRNA
minus
prg-1 OP50 sRNA
replicate #1
replicate #2
replicate #3

2G

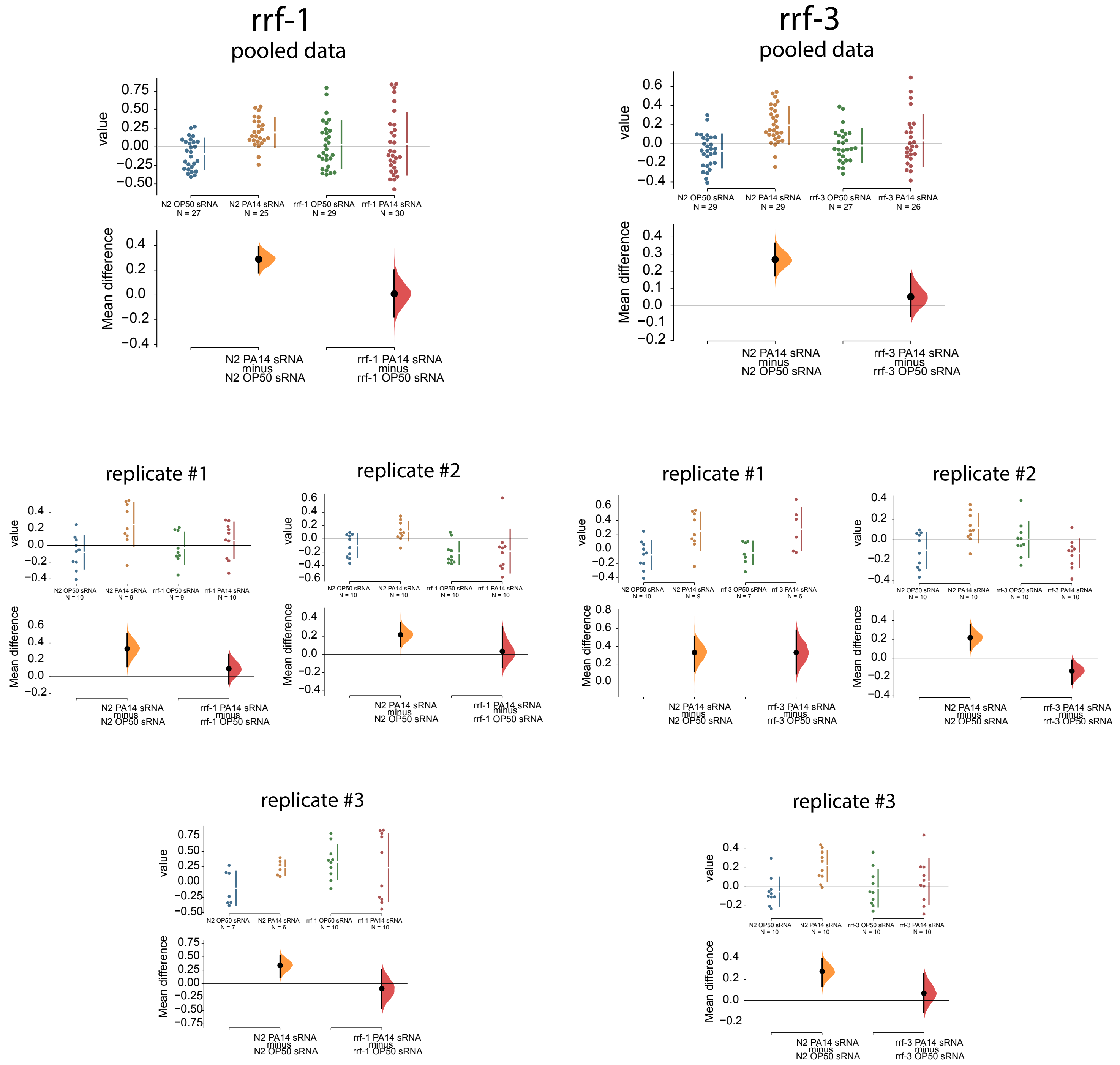
rrf-1
pooled data
rrf-3
pooled data
replicate #1
replicate #2
replicate #3
value
Mean difference
N2 OP50 sRNA
N2 PA14 sRNA
rrf-1 OP50 sRNA
rrf-1 PA14 sRNA
rrf-3 OP50 sRNA
rrf-3 PA14 sRNA
N = 27
N = 25
N = 29
N = 30
N = 26
N = 10
N = 9
N = 7
N = 6
N2 PA14 sRNA minus N2 OP50 sRNA
rrf-1 PA14 sRNA minus rrf-1 OP50 sRNA
rrf-3 PA14 sRNA minus rrf-3 OP50 sRNA

2H

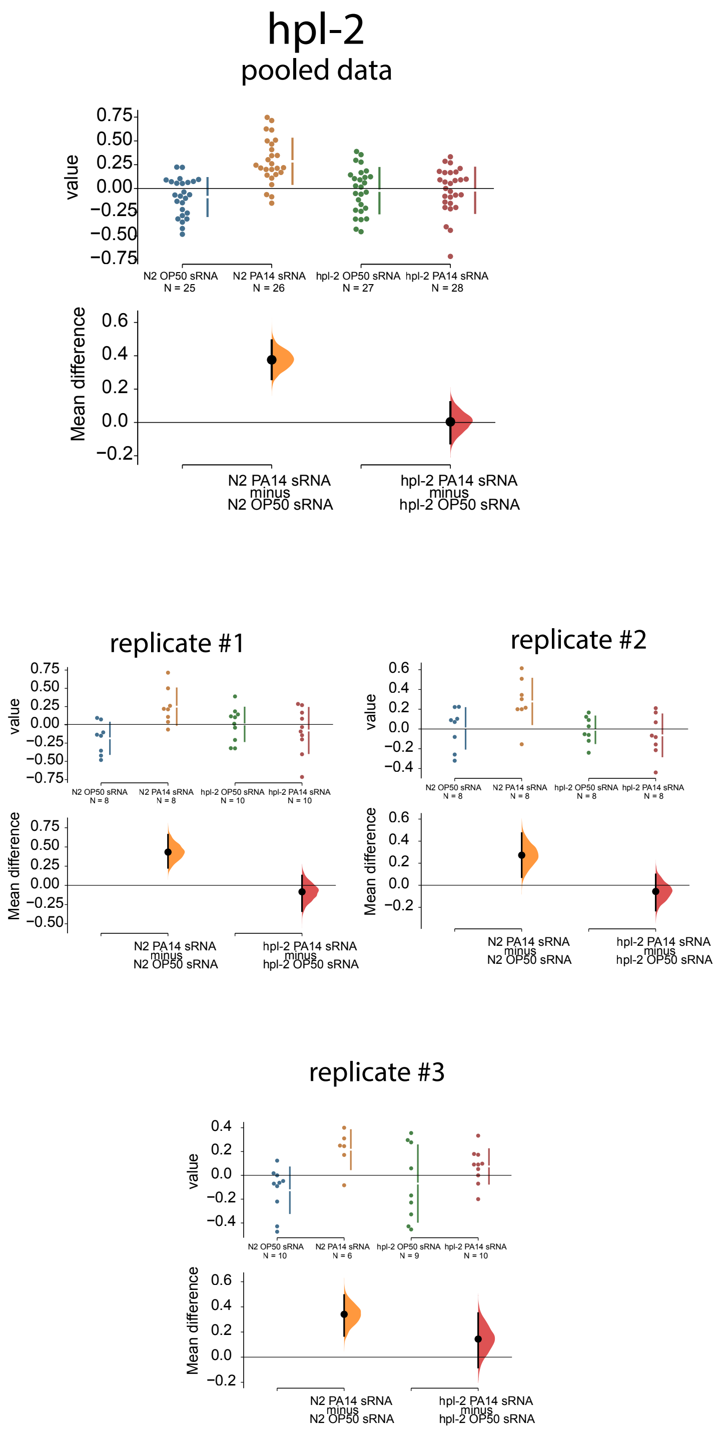

hpl-2
pooled data
value
Mean difference
N2 OP50 sRNA
N = 25
N2 PA14 sRNA
N = 26
hpl-2 OP50 sRNA
N = 27
hpl-2 PA14 sRNA
N = 28
N2 PA14 sRNA minus N2 OP50 sRNA
hpl-2 PA14 sRNA minus hpl-2 OP50 sRNA
replicate #1
N2 OP50 sRNA N = 8
N2 PA14 sRNA N = 8
hpl-2 OP50 sRNA N = 10
hpl-2 PA14 sRNA N = 10
replicate #2
N2 OP50 sRNA N = 8
N2 PA14 sRNA N = 8
hpl-2 OP50 sRNA N = 8
hpl-2 PA14 sRNA N = 8
replicate #3
N2 OP50 sRNA N = 10
N2 PA14 sRNA N = 6
hpl-2 OP50 sRNA N = 9
hpl-2 PA14 sRNA N = 10

2K/L

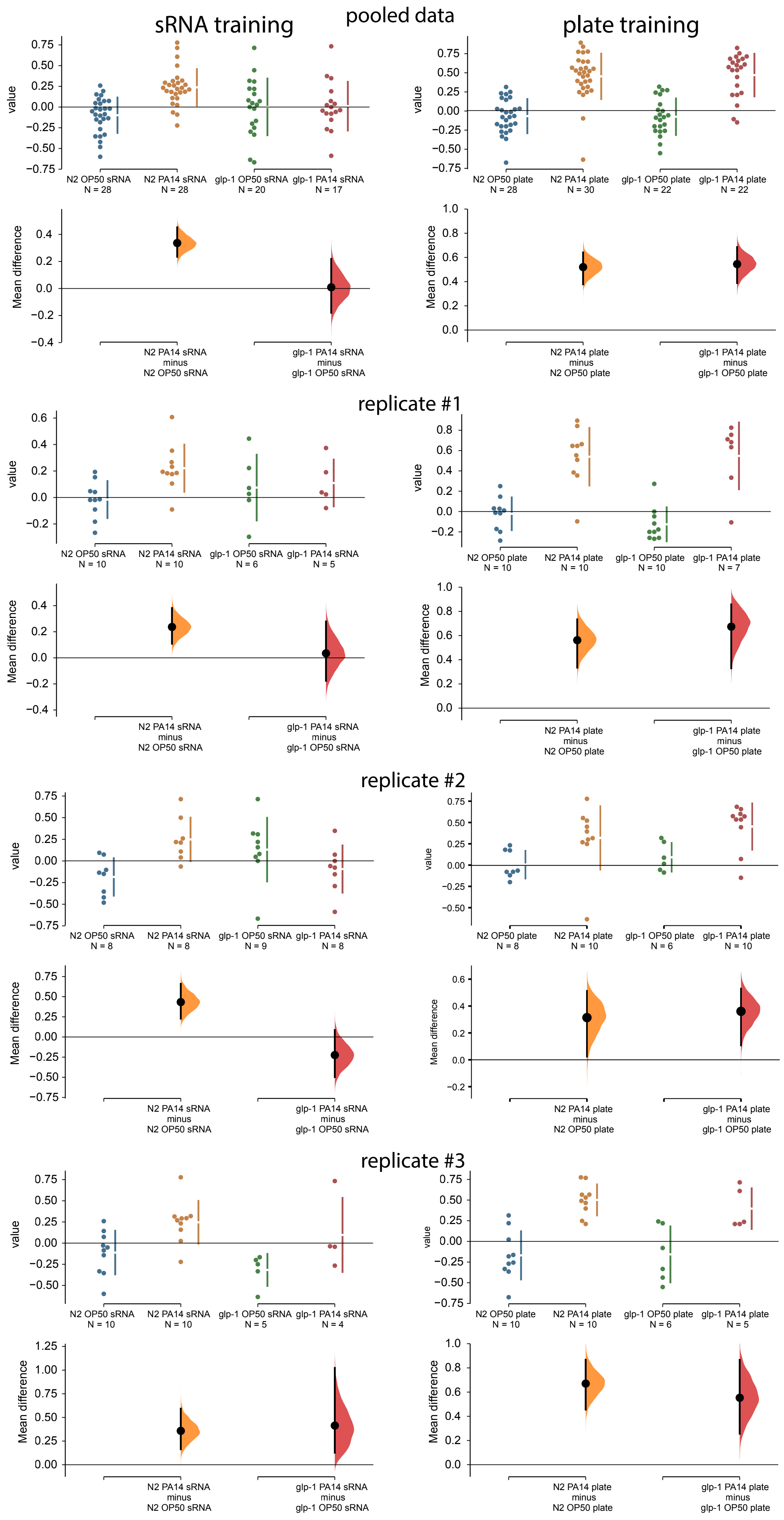

sRNA training
pooled data
plate training
replicate #1
replicate #2
replicate #3
value
Mean difference
N2 OP50 sRNA
N2 PA14 sRNA
glp-1 OP50 sRNA
glp-1 PA14 sRNA
N2 OP50 plate
N2 PA14 plate
glp-1 OP50 plate
glp-1 PA14 plate
N2 PA14 sRNA minus N2 OP50 sRNA
glp-1 PA14 sRNA minus glp-1 OP50 sRNA
N2 PA14 plate minus N2 OP50 plate
glp-1 PA14 plate minus glp-1 OP50 plate
N = 28
N = 28
N = 20
N = 17
N = 28
N = 30
N = 22
N = 22
N = 10
N = 10
N = 6
N = 5
N = 10
N = 10
N = 10
N = 7
N = 8
N = 8
N = 9
N = 8
N = 8
N = 10
N = 6
N = 10
N = 10
N = 10
N = 5
N = 4
N = 10
N = 10
N = 6
N = 5

2O

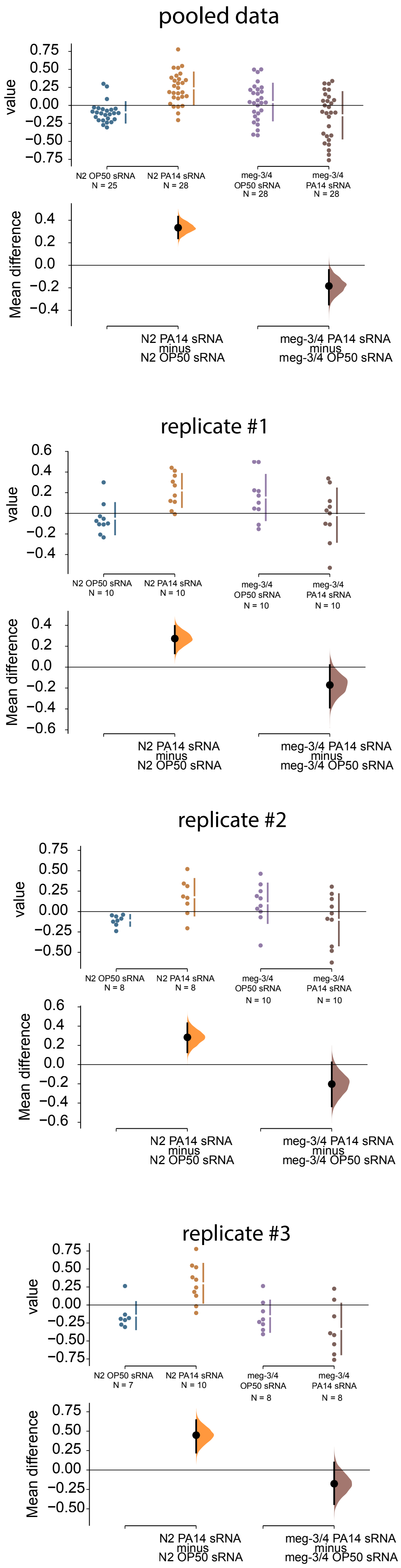
pooled data
value
0.75
0.50
0.25
0.00
−0.25
−0.50
−0.75
N2 OP50 sRNA
N = 25
N2 PA14 sRNA
N = 28
meg-3/4
OP50 sRNA
N = 28
meg-3/4
PA14 sRNA
N = 28
Mean difference
0.4
0.2
0.0
−0.2
−0.4
N2 PA14 sRNA
minus
N2 OP50 sRNA
meg-3/4 PA14 sRNA
minus
meg-3/4 OP50 sRNA
replicate #1
value
0.6
0.4
0.2
0.0
−0.2
−0.4
N2 OP50 sRNA
N = 10
N2 PA14 sRNA
N = 10
meg-3/4
OP50 sRNA
N = 10
meg-3/4
PA14 sRNA
N = 10
Mean difference
0.4
0.2
0.0
−0.2
−0.4
−0.6
N2 PA14 sRNA
minus
N2 OP50 sRNA
meg-3/4 PA14 sRNA
minus
meg-3/4 OP50 sRNA
replicate #2
value
0.75
0.50
0.25
0.00
−0.25
−0.50
N2 OP50 sRNA
N = 8
N2 PA14 sRNA
N = 8
meg-3/4
OP50 sRNA
N = 10
meg-3/4
PA14 sRNA
N = 10
Mean difference
0.6
0.4
0.2
0.0
−0.2
−0.4
−0.6
N2 PA14 sRNA
minus
N2 OP50 sRNA
meg-3/4 PA14 sRNA
minus
meg-3/4 OP50 sRNA
replicate #3
value
0.75
0.50
0.25
0.00
−0.25
−0.50
−0.75
N2 OP50 sRNA
N = 7
N2 PA14 sRNA
N = 10
meg-3/4
OP50 sRNA
N = 8
meg-3/4
PA14 sRNA
N = 8
Mean difference
0.75
0.50
0.25
0.00
−0.25
−0.50
N2 PA14 sRNA
minus
N2 OP50 sRNA
meg-3/4 PA14 sRNA
minus
meg-3/4 OP50 sRNA

2P

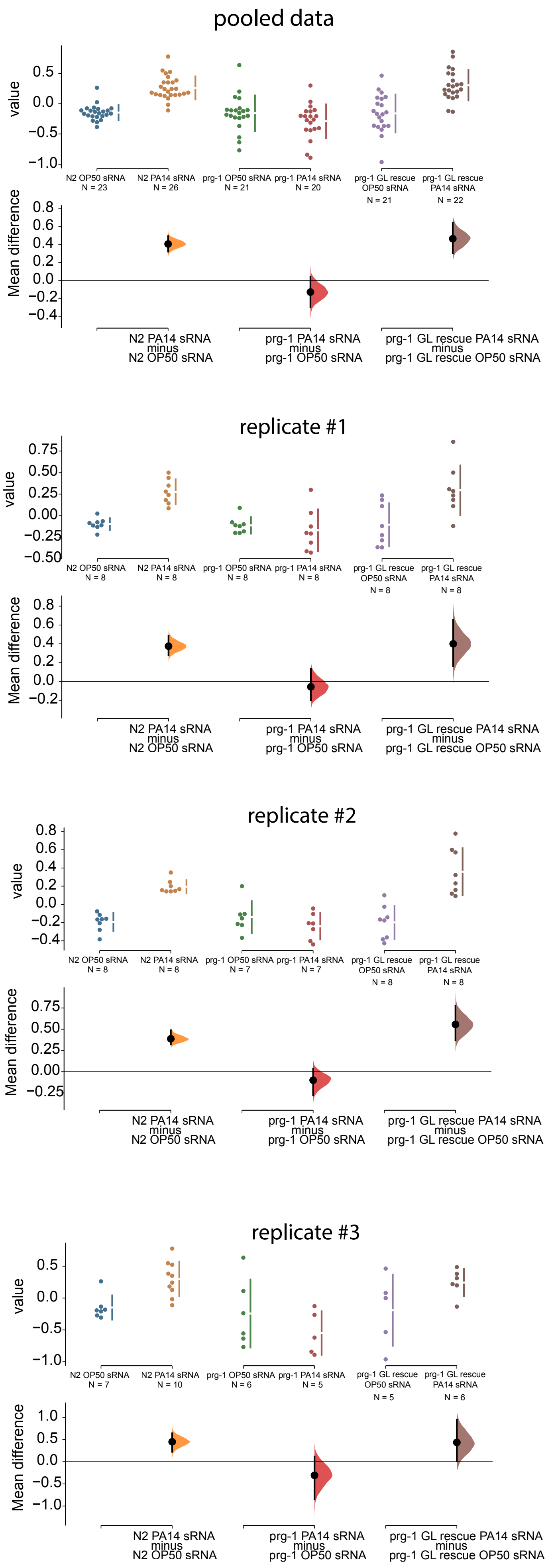

pooled data
value
N2 OP50 sRNA N = 23
N2 PA14 sRNA N = 26
prg-1 OP50 sRNA N = 21
prg-1 PA14 sRNA N = 20
prg-1 GL rescue OP50 sRNA N = 21
prg-1 GL rescue PA14 sRNA N = 22
Mean difference
N2 PA14 sRNA minus N2 OP50 sRNA
prg-1 PA14 sRNA minus prg-1 OP50 sRNA
prg-1 GL rescue PA14 sRNA minus prg-1 GL rescue OP50 sRNA
replicate #1
value
N2 OP50 sRNA N = 8
N2 PA14 sRNA N = 8
prg-1 OP50 sRNA N = 8
prg-1 PA14 sRNA N = 8
prg-1 GL rescue OP50 sRNA N = 8
prg-1 GL rescue PA14 sRNA N = 8
Mean difference
N2 PA14 sRNA minus N2 OP50 sRNA
prg-1 PA14 sRNA minus prg-1 OP50 sRNA
prg-1 GL rescue PA14 sRNA minus prg-1 GL rescue OP50 sRNA
replicate #2
value
N2 OP50 sRNA N = 8
N2 PA14 sRNA N = 8
prg-1 OP50 sRNA N = 7
prg-1 PA14 sRNA N = 7
prg-1 GL rescue OP50 sRNA N = 8
prg-1 GL rescue PA14 sRNA N = 8
Mean difference
N2 PA14 sRNA minus N2 OP50 sRNA
prg-1 PA14 sRNA minus prg-1 OP50 sRNA
prg-1 GL rescue PA14 sRNA minus prg-1 GL rescue OP50 sRNA
replicate #3
value
N2 OP50 sRNA N = 7
N2 PA14 sRNA N = 10
prg-1 OP50 sRNA N = 6
prg-1 PA14 sRNA N = 5
prg-1 GL rescue OP50 sRNA N = 5
prg-1 GL rescue PA14 sRNA N = 6
Mean difference
N2 PA14 sRNA minus N2 OP50 sRNA
prg-1 PA14 sRNA minus prg-1 OP50 sRNA
prg-1 GL rescue PA14 sRNA minus prg-1 GL rescue OP50 sRNA

# 3A/B

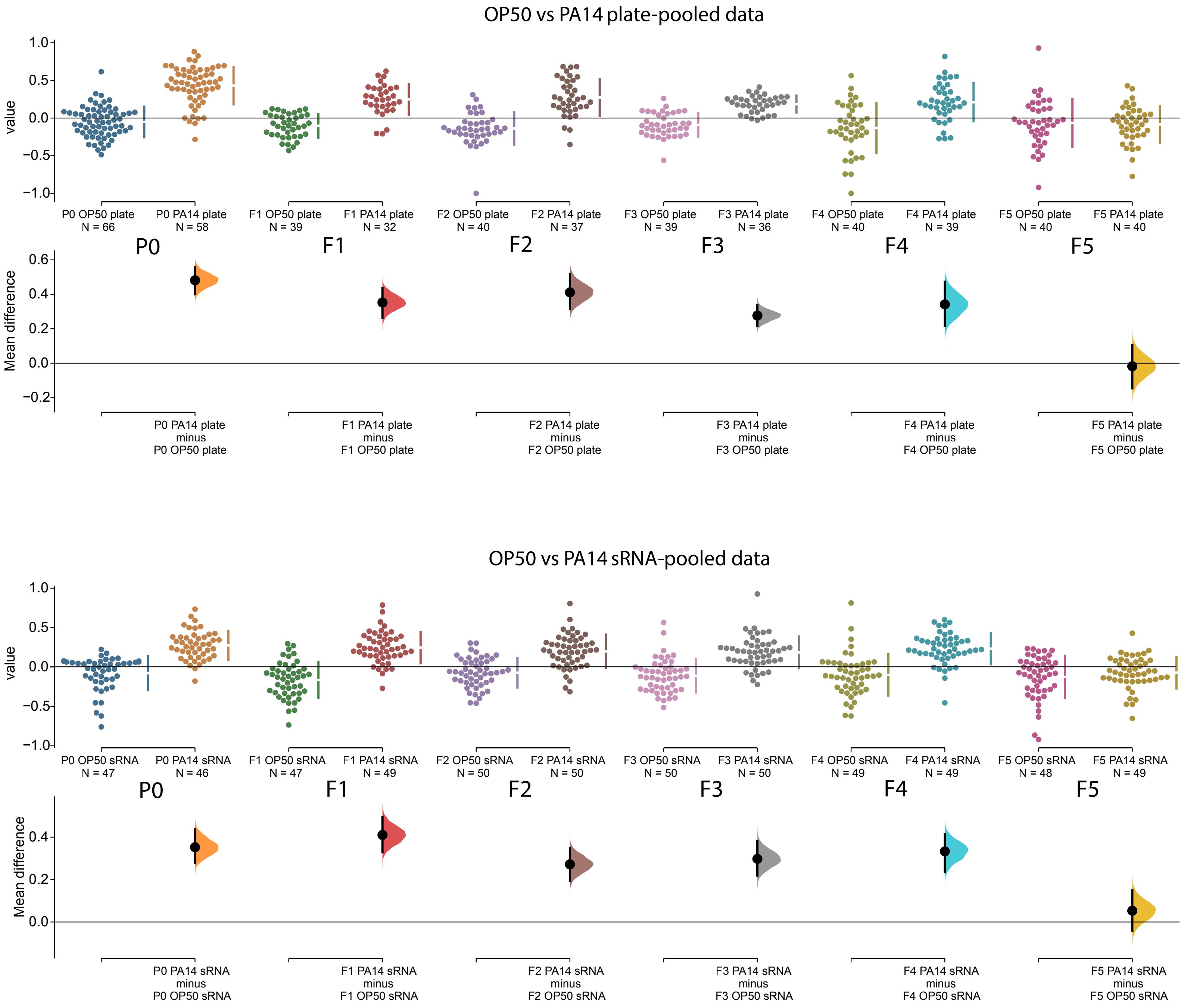
OP50 vs PA14 plate-pooled data
value
Mean difference
P0 OP50 plate N = 66
P0 PA14 plate N = 58
F1 OP50 plate N = 39
F1 PA14 plate N = 32
F2 OP50 plate N = 40
F2 PA14 plate N = 37
F3 OP50 plate N = 39
F3 PA14 plate N = 36
F4 OP50 plate N = 40
F4 PA14 plate N = 39
F5 OP50 plate N = 40
F5 PA14 plate N = 40
P0
F1
F2
F3
F4
F5
P0 PA14 plate minus P0 OP50 plate
F1 PA14 plate minus F1 OP50 plate
F2 PA14 plate minus F2 OP50 plate
F3 PA14 plate minus F3 OP50 plate
F4 PA14 plate minus F4 OP50 plate
F5 PA14 plate minus F5 OP50 plate
OP50 vs PA14 sRNA-pooled data
value
Mean difference
P0 OP50 sRNA N = 47
P0 PA14 sRNA N = 46
F1 OP50 sRNA N = 47
F1 PA14 sRNA N = 49
F2 OP50 sRNA N = 50
F2 PA14 sRNA N = 50
F3 OP50 sRNA N = 50
F3 PA14 sRNA N = 50
F4 OP50 sRNA N = 49
F4 PA14 sRNA N = 49
F5 OP50 sRNA N = 48
F5 PA14 sRNA N = 49
P0
F1
F2
F3
F4
F5
P0 PA14 sRNA minus P0 OP50 sRNA
F1 PA14 sRNA minus F1 OP50 sRNA
F2 PA14 sRNA minus F2 OP50 sRNA
F3 PA14 sRNA minus F3 OP50 sRNA
F4 PA14 sRNA minus F4 OP50 sRNA
F5 PA14 sRNA minus F5 OP50 sRNA

4A

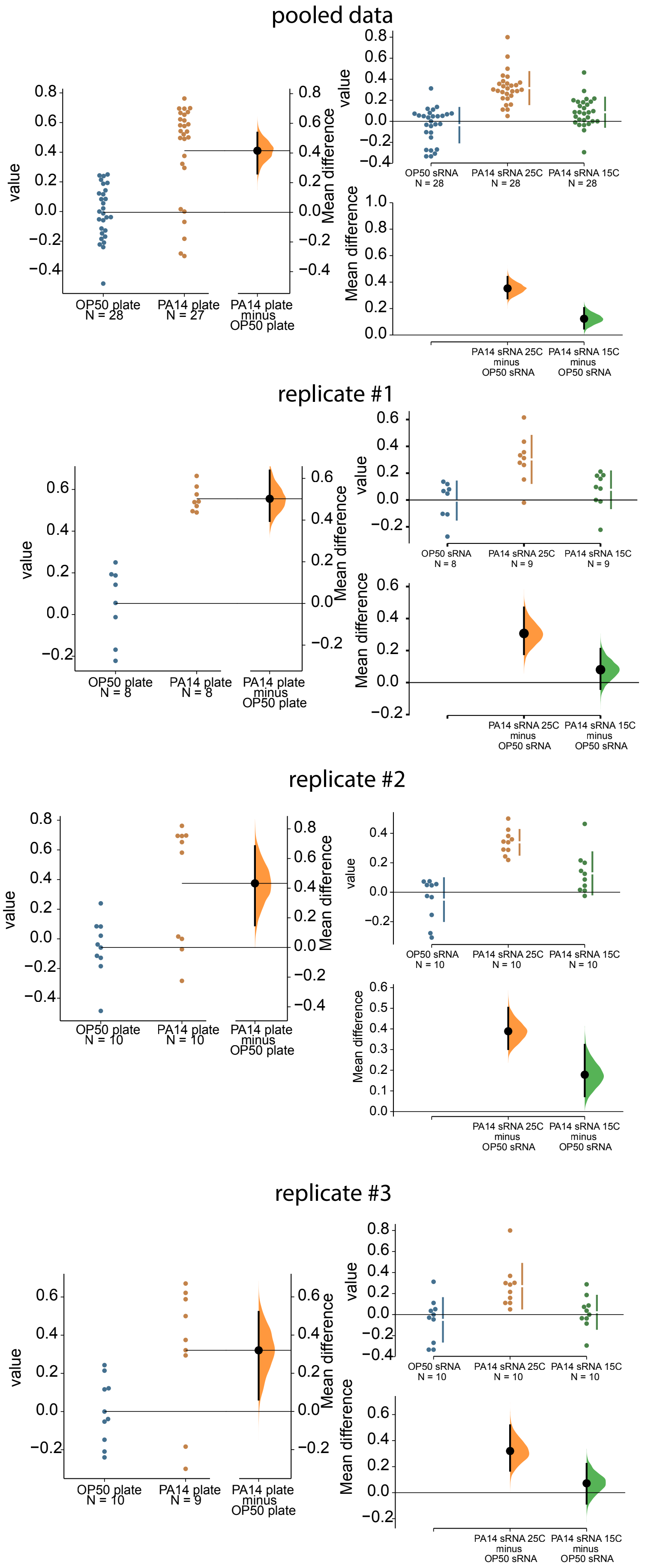
pooled data
value
Mean difference
OP50 plate
N = 28
PA14 plate
N = 27
PA14 plate
minus
OP50 plate
OP50 sRNA
N = 28
PA14 sRNA 25C
N = 28
PA14 sRNA 15C
N = 28
PA14 sRNA 25C
minus
OP50 sRNA
PA14 sRNA 15C
minus
OP50 sRNA
replicate #1
value
Mean difference
OP50 plate
N = 8
PA14 plate
N = 8
PA14 plate
minus
OP50 plate
OP50 sRNA
N = 8
PA14 sRNA 25C
N = 9
PA14 sRNA 15C
N = 9
PA14 sRNA 25C
minus
OP50 sRNA
PA14 sRNA 15C
minus
OP50 sRNA
replicate #2
value
Mean difference
OP50 plate
N = 10
PA14 plate
N = 10
PA14 plate
minus
OP50 plate
OP50 sRNA
N = 10
PA14 sRNA 25C
N = 10
PA14 sRNA 15C
N = 10
PA14 sRNA 25C
minus
OP50 sRNA
PA14 sRNA 15C
minus
OP50 sRNA
replicate #3
value
Mean difference
OP50 plate
N = 10
PA14 plate
N = 9
PA14 plate
minus
OP50 plate
OP50 sRNA
N = 10
PA14 sRNA 25C
N = 10
PA14 sRNA 15C
N = 10
PA14 sRNA 25C
minus
OP50 sRNA
PA14 sRNA 15C
minus
OP50 sRNA

4B

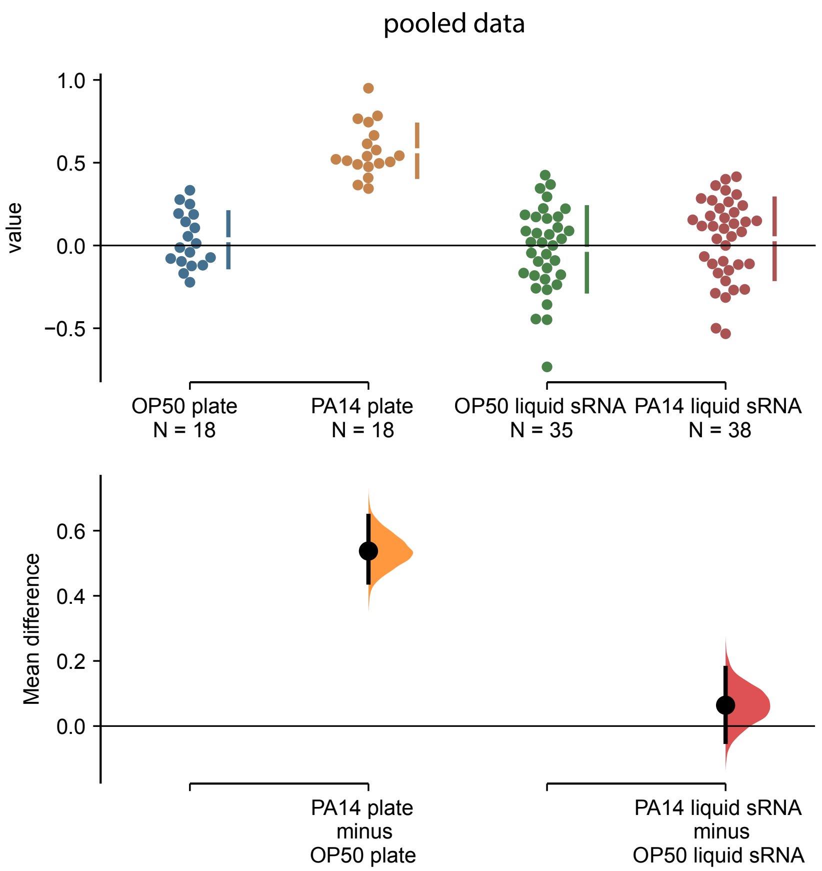
pooled data
value
1.0
0.5
0.0
−0.5
OP50 plate
N = 18
PA14 plate
N = 18
OP50 liquid sRNA
N = 35
PA14 liquid sRNA
N = 38
Mean difference
0.6
0.4
0.2
0.0
PA14 plate
minus
OP50 plate
PA14 liquid sRNA
minus
OP50 liquid sRNA

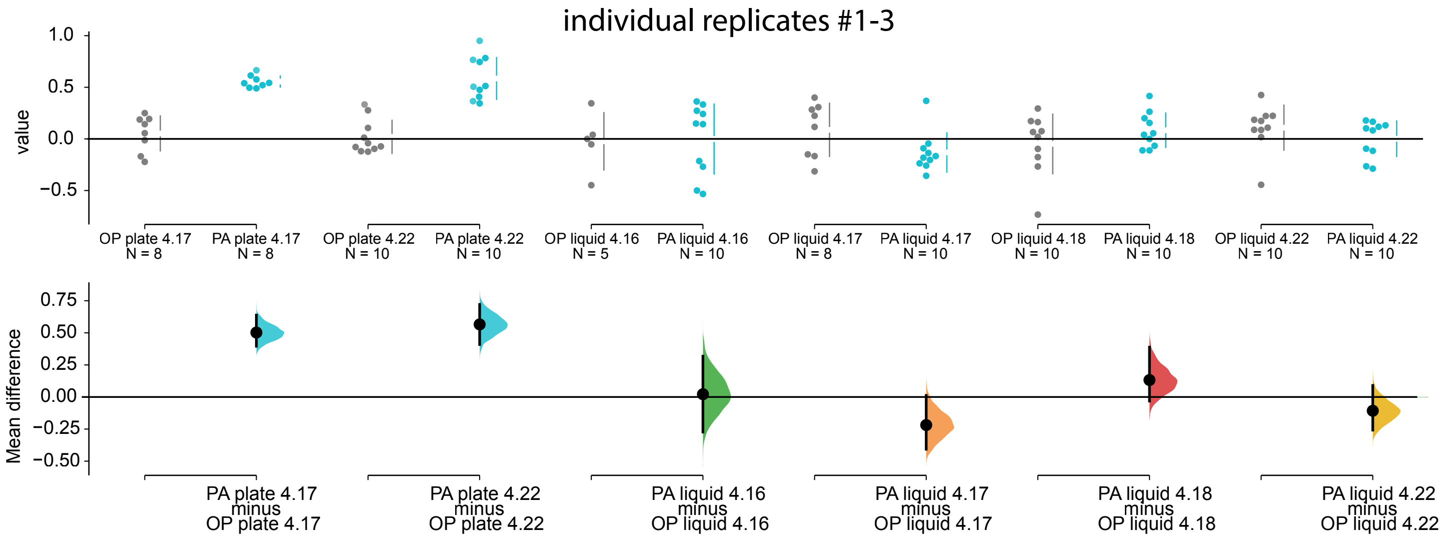
individual replicates #1-3
value
1.0
0.5
0.0
−0.5
OP plate 4.17
N = 8
PA plate 4.17
N = 8
OP plate 4.22
N = 10
PA plate 4.22
N = 10
OP liquid 4.16
N = 5
PA liquid 4.16
N = 10
OP liquid 4.17
N = 8
PA liquid 4.17
N = 10
OP liquid 4.18
N = 10
PA liquid 4.18
N = 10
OP liquid 4.22
N = 10
PA liquid 4.22
N = 10
Mean difference
0.75
0.50
0.25
0.00
−0.25
−0.50
PA plate 4.17
minus
OP plate 4.17
PA plate 4.22
minus
OP plate 4.22
PA liquid 4.16
minus
OP liquid 4.16
PA liquid 4.17
minus
OP liquid 4.17
PA liquid 4.18
minus
OP liquid 4.18
PA liquid 4.22
minus
OP liquid 4.22

4F

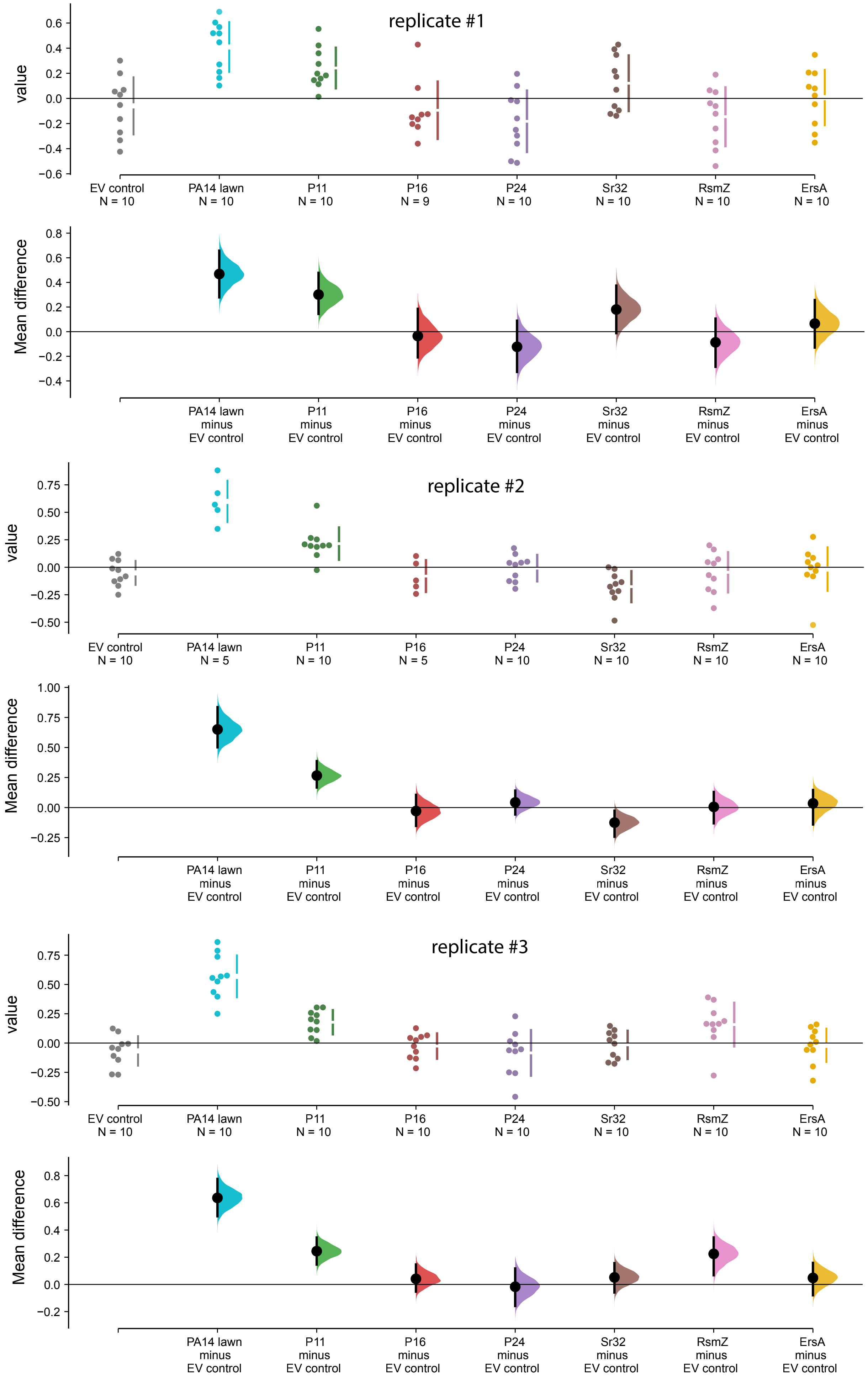
replicate #1
value
EV control
N = 10
PA14 lawn
N = 10
P11
N = 10
P16
N = 9
P24
N = 10
Sr32
N = 10
RsmZ
N = 10
ErsA
N = 10
Mean difference
PA14 lawn
minus
EV control
P11
minus
EV control
P16
minus
EV control
P24
minus
EV control
Sr32
minus
EV control
RsmZ
minus
EV control
ErsA
minus
EV control
replicate #2
value
EV control
N = 10
PA14 lawn
N = 5
P11
N = 10
P16
N = 5
P24
N = 10
Sr32
N = 10
RsmZ
N = 10
ErsA
N = 10
Mean difference
PA14 lawn
minus
EV control
P11
minus
EV control
P16
minus
EV control
P24
minus
EV control
Sr32
minus
EV control
RsmZ
minus
EV control
ErsA
minus
EV control
replicate #3
value
EV control
N = 10
PA14 lawn
N = 10
P11
N = 10
P16
N = 10
P24
N = 10
Sr32
N = 10
RsmZ
N = 10
ErsA
N = 10
Mean difference
PA14 lawn
minus
EV control
P11
minus
EV control
P16
minus
EV control
P24
minus
EV control
Sr32
minus
EV control
RsmZ
minus
EV control
ErsA
minus
EV control

4H

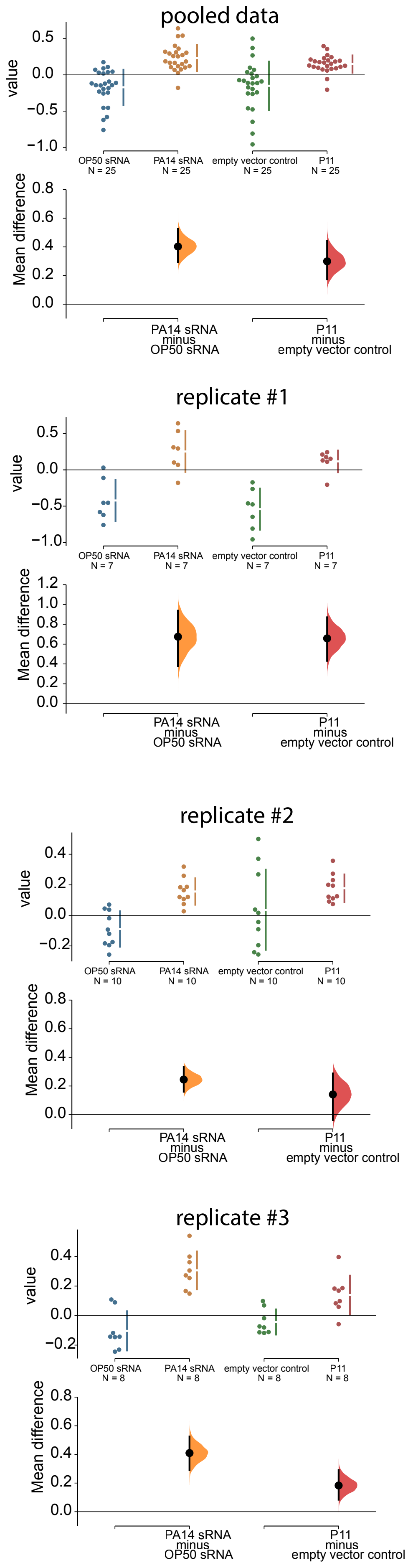
pooled data
value
OP50 sRNA N = 25
PA14 sRNA N = 25
empty vector control N = 25
P11 N = 25
Mean difference
PA14 sRNA minus OP50 sRNA
P11 minus empty vector control
replicate #1
OP50 sRNA N = 7
PA14 sRNA N = 7
empty vector control N = 7
P11 N = 7
replicate #2
OP50 sRNA N = 10
PA14 sRNA N = 10
empty vector control N = 10
P11 N = 10
replicate #3
OP50 sRNA N = 8
PA14 sRNA N = 8
empty vector control N = 8
P11 N = 8

4J

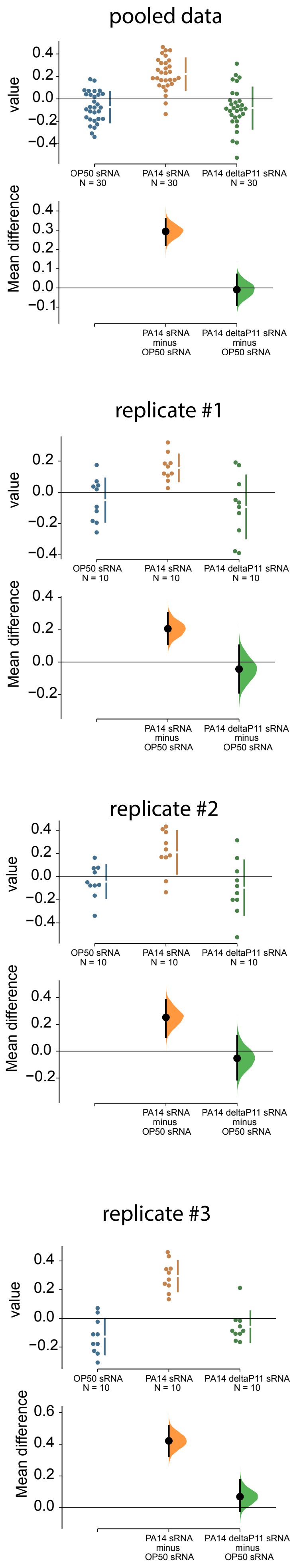
pooled data
value
Mean difference
OP50 sRNA
N = 30
PA14 sRNA
N = 30
PA14 deltaP11 sRNA
N = 30
PA14 sRNA
minus
OP50 sRNA
PA14 deltaP11 sRNA
minus
OP50 sRNA
replicate #1
value
Mean difference
OP50 sRNA
N = 10
PA14 sRNA
N = 10
PA14 deltaP11 sRNA
N = 10
PA14 sRNA
minus
OP50 sRNA
PA14 deltaP11 sRNA
minus
OP50 sRNA
replicate #2
value
Mean difference
OP50 sRNA
N = 10
PA14 sRNA
N = 10
PA14 deltaP11 sRNA
N = 10
PA14 sRNA
minus
OP50 sRNA
PA14 deltaP11 sRNA
minus
OP50 sRNA
replicate #3
value
Mean difference
OP50 sRNA
N = 10
PA14 sRNA
N = 10
PA14 deltaP11 sRNA
N = 10
PA14 sRNA
minus
OP50 sRNA
PA14 deltaP11 sRNA
minus
OP50 sRNA

# 5A/B

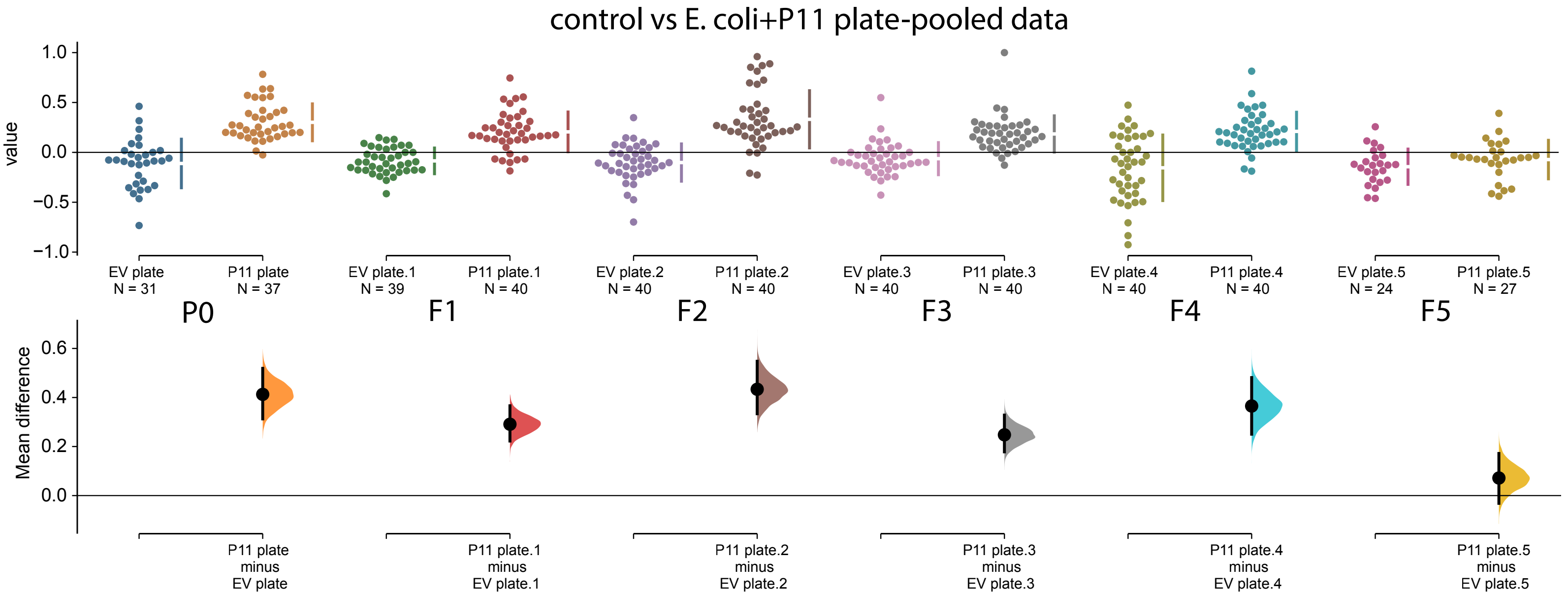

control vs E. coli+P11 plate-pooled data
value
Mean difference
EV plate N = 31
P11 plate N = 37
EV plate.1 N = 39
P11 plate.1 N = 40
EV plate.2 N = 40
P11 plate.2 N = 40
EV plate.3 N = 40
P11 plate.3 N = 40
EV plate.4 N = 40
P11 plate.4 N = 40
EV plate.5 N = 24
P11 plate.5 N = 27
P0
F1
F2
F3
F4
F5
P11 plate minus EV plate
P11 plate.1 minus EV plate.1
P11 plate.2 minus EV plate.2
P11 plate.3 minus EV plate.3
P11 plate.4 minus EV plate.4
P11 plate.5 minus EV plate.5


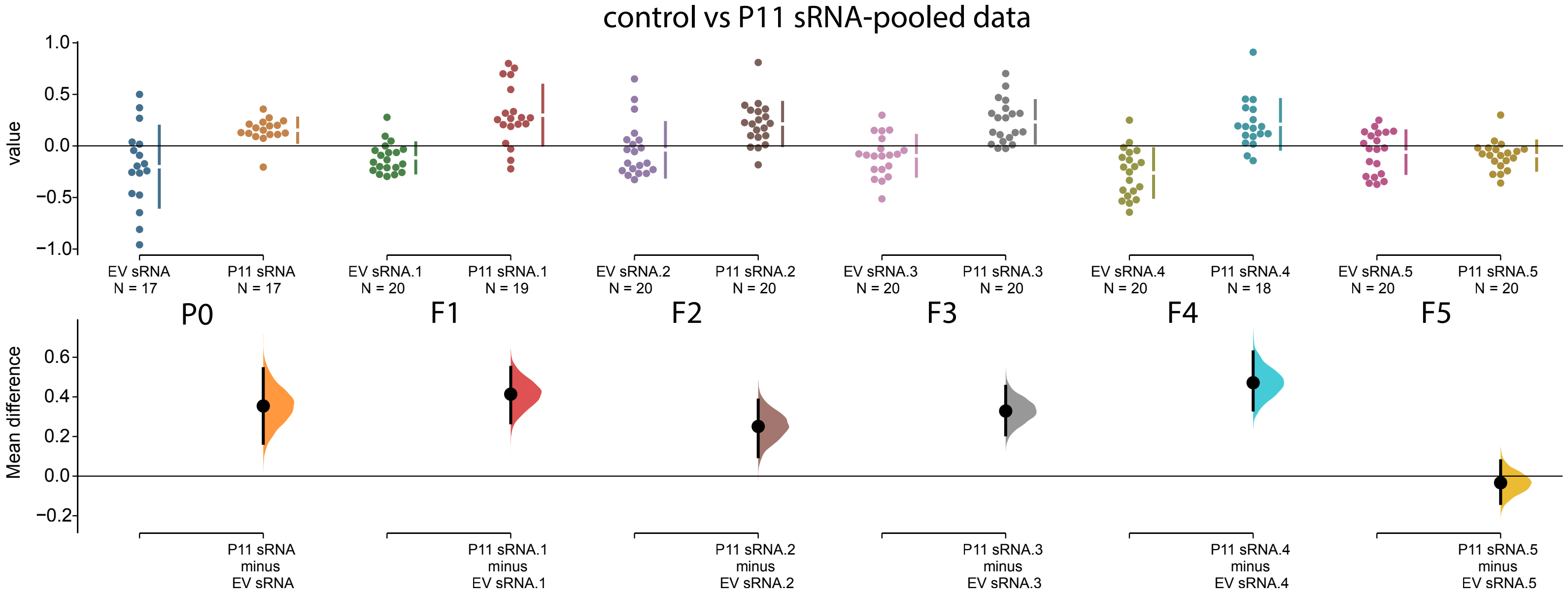

control vs P11 sRNA-pooled data
value
Mean difference
EV sRNA N = 17
P11 sRNA N = 17
EV sRNA.1 N = 20
P11 sRNA.1 N = 19
EV sRNA.2 N = 20
P11 sRNA.2 N = 20
EV sRNA.3 N = 20
P11 sRNA.3 N = 20
EV sRNA.4 N = 20
P11 sRNA.4 N = 18
EV sRNA.5 N = 20
P11 sRNA.5 N = 20
P0
F1
F2
F3
F4
F5
P11 sRNA minus EV sRNA
P11 sRNA.1 minus EV sRNA.1
P11 sRNA.2 minus EV sRNA.2
P11 sRNA.3 minus EV sRNA.3
P11 sRNA.4 minus EV sRNA.4
P11 sRNA.5 minus EV sRNA.5

# 5C

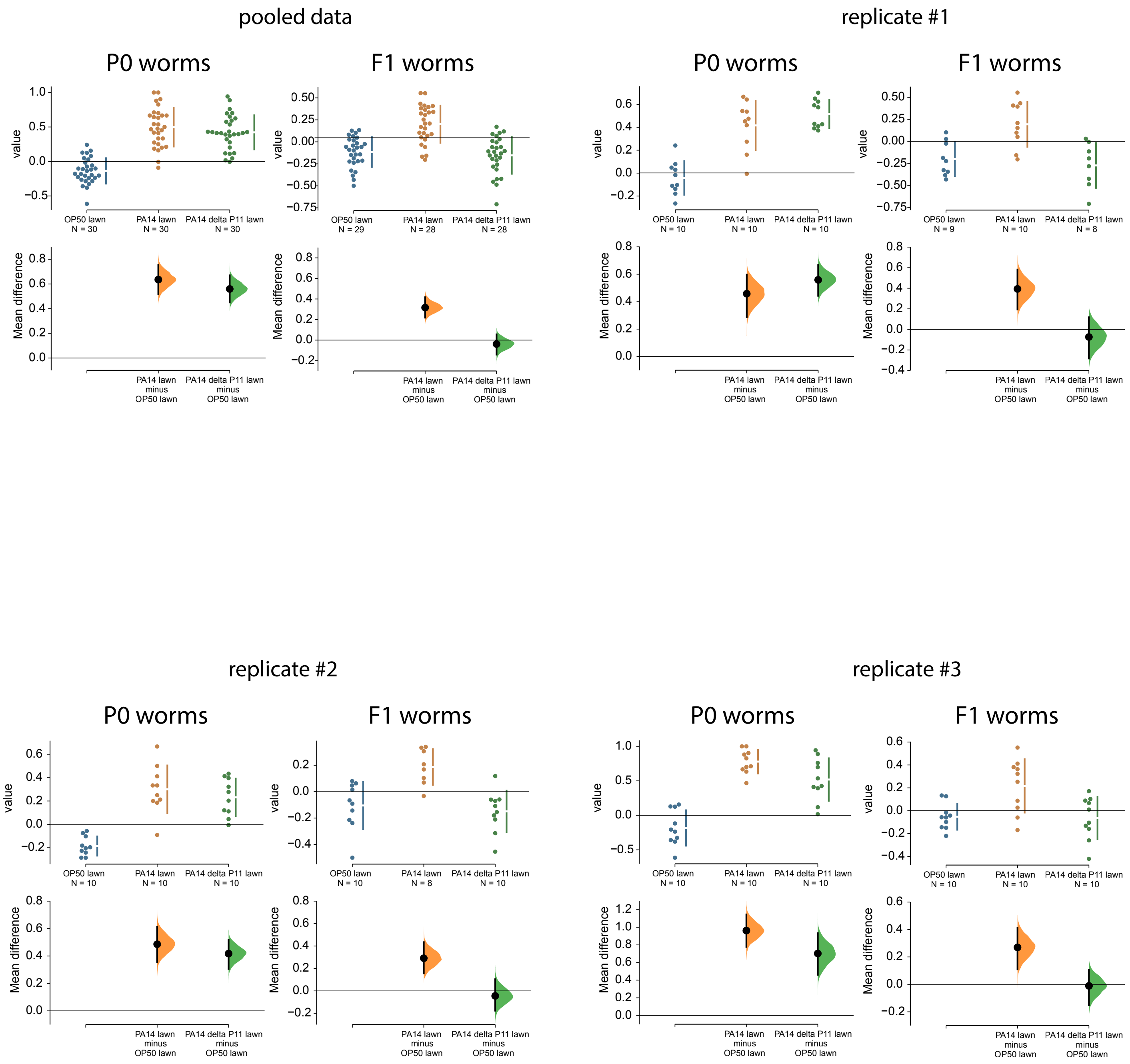

pooled data
replicate #1
replicate #2
replicate #3
P0 worms
F1 worms
value
Mean difference
OP50 lawn
PA14 lawn
PA14 delta P11 lawn
N = 30
N = 29
N = 28
N = 10
N = 9
N = 8
PA14 lawn minus OP50 lawn
PA14 delta P11 lawn minus OP50 lawn

6B

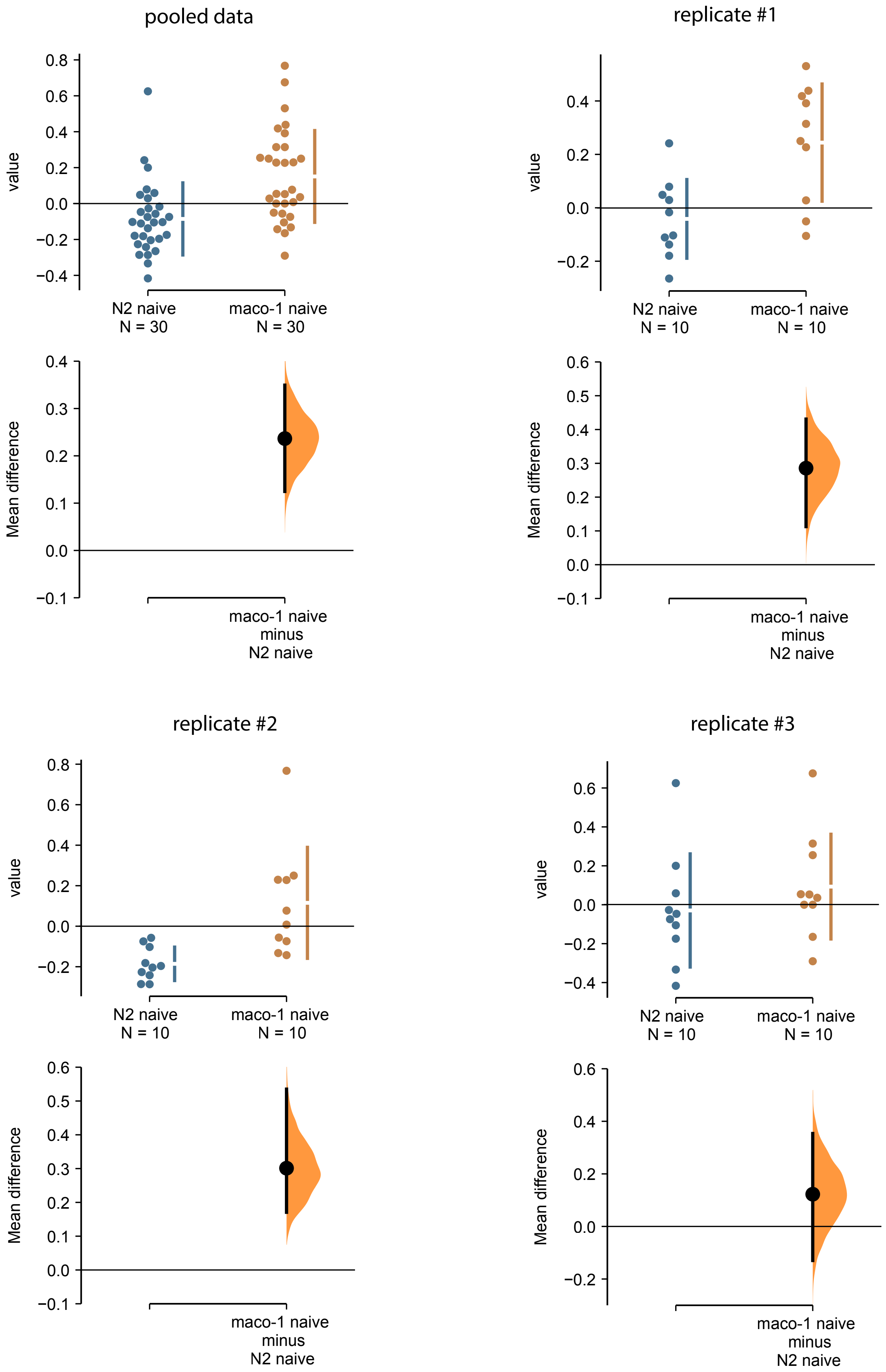

pooled data
replicate #1
replicate #2
replicate #3
value
Mean difference
N2 naive
N = 30
maco-1 naive
N = 30
N = 10
maco-1 naive
minus
N2 naive
0.8
0.6
0.4
0.2
0.0
−0.2
−0.4
0.5
0.3
0.1
−0.1

6C

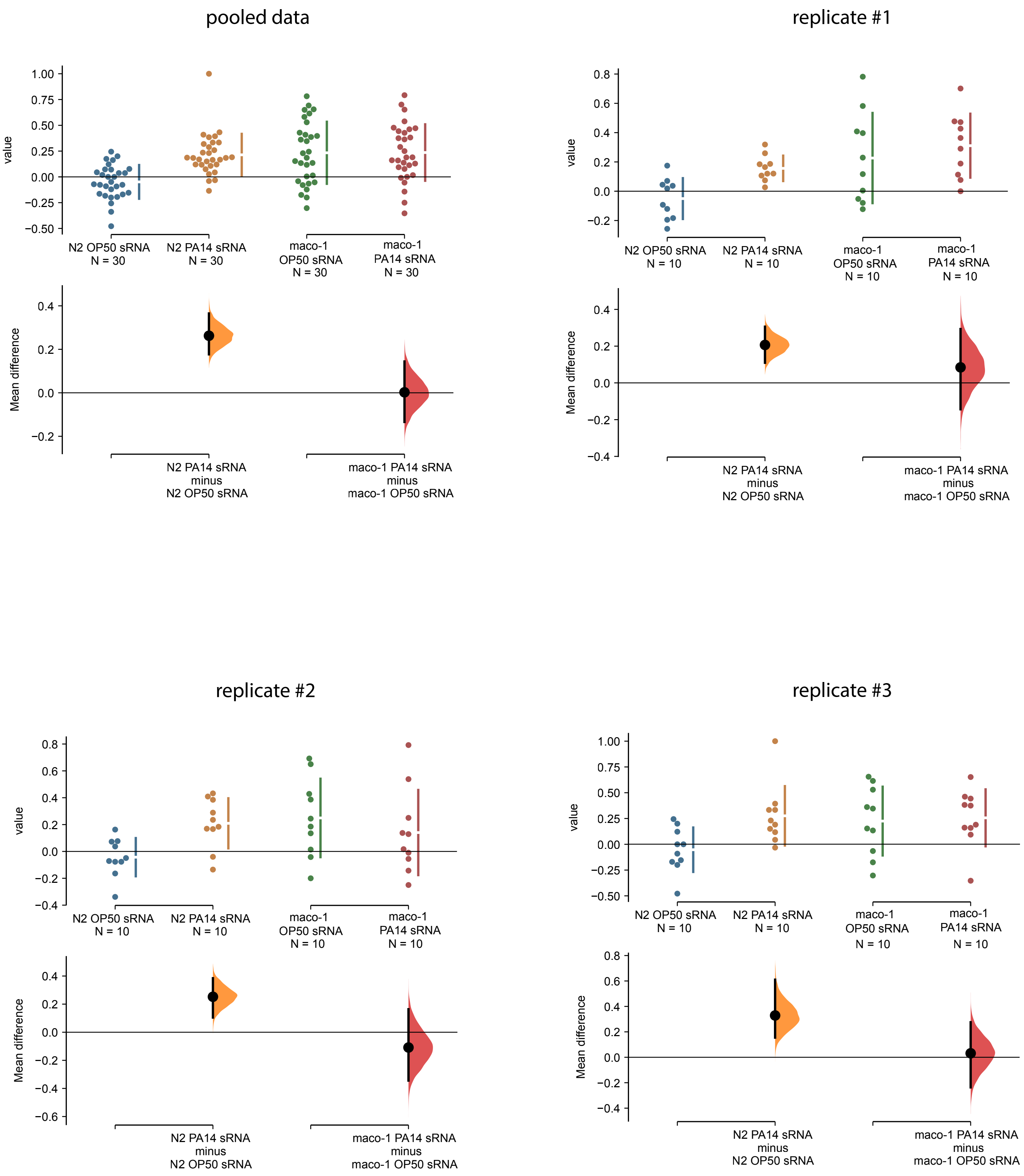
pooled data
value
Mean difference
N2 OP50 sRNA
N = 30
N2 PA14 sRNA
N = 30
maco-1
OP50 sRNA
N = 30
maco-1
PA14 sRNA
N = 30
N2 PA14 sRNA
minus
N2 OP50 sRNA
maco-1 PA14 sRNA
minus
maco-1 OP50 sRNA
replicate #1
value
Mean difference
N2 OP50 sRNA
N = 10
N2 PA14 sRNA
N = 10
maco-1
OP50 sRNA
N = 10
maco-1
PA14 sRNA
N = 10
N2 PA14 sRNA
minus
N2 OP50 sRNA
maco-1 PA14 sRNA
minus
maco-1 OP50 sRNA
replicate #2
value
Mean difference
N2 OP50 sRNA
N = 10
N2 PA14 sRNA
N = 10
maco-1
OP50 sRNA
N = 10
maco-1
PA14 sRNA
N = 10
N2 PA14 sRNA
minus
N2 OP50 sRNA
maco-1 PA14 sRNA
minus
maco-1 OP50 sRNA
replicate #3
value
Mean difference
N2 OP50 sRNA
N = 10
N2 PA14 sRNA
N = 10
maco-1
OP50 sRNA
N = 10
maco-1
PA14 sRNA
N = 10
N2 PA14 sRNA
minus
N2 OP50 sRNA
maco-1 PA14 sRNA
minus
maco-1 OP50 sRNA

6E

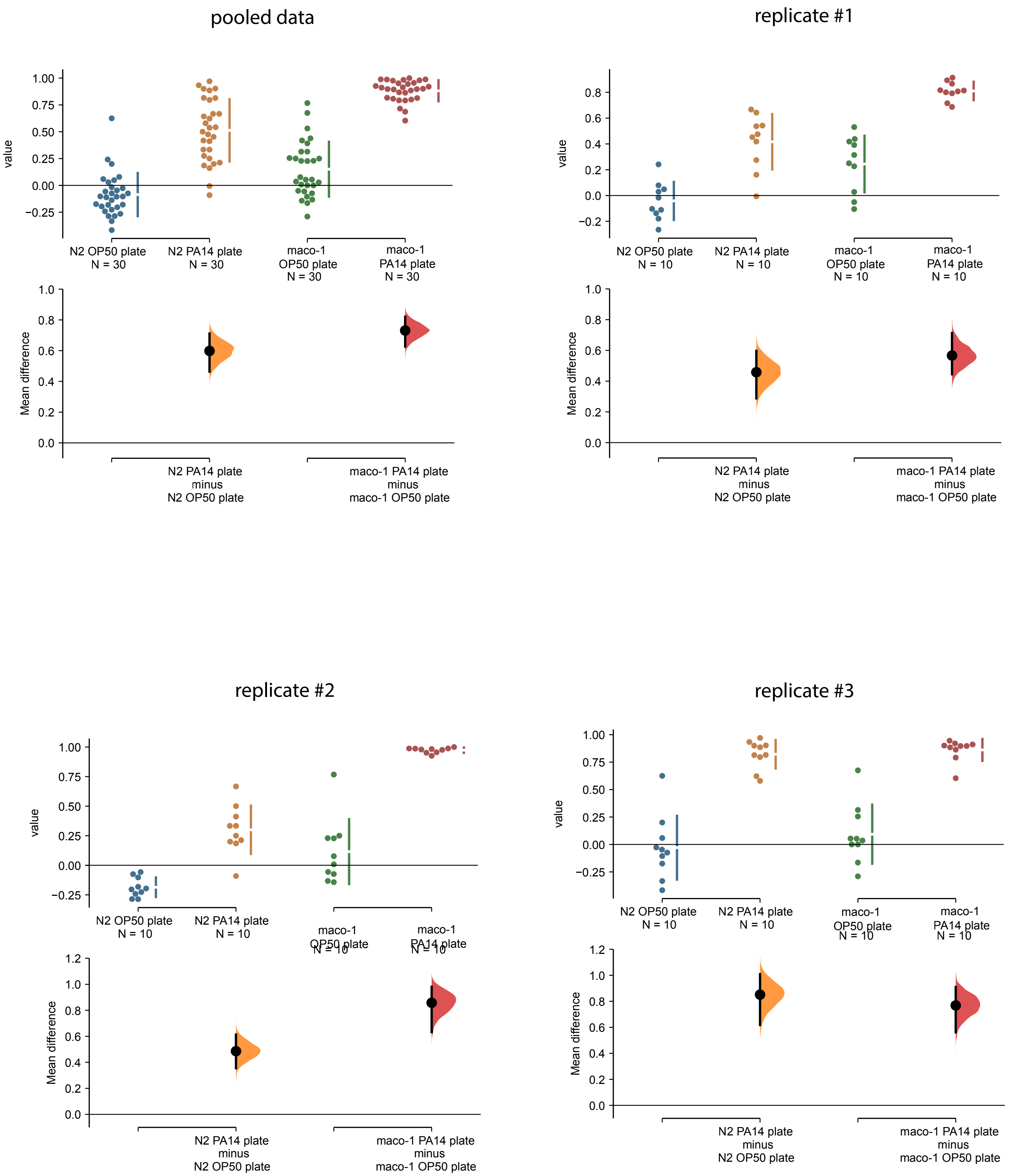

pooled data
value
1.00
0.75
0.50
0.25
0.00
−0.25
N2 OP50 plate
N = 30
N2 PA14 plate
N = 30
maco-1
OP50 plate
N = 30
maco-1
PA14 plate
N = 30
Mean difference
1.0
0.8
0.6
0.4
0.2
0.0
N2 PA14 plate
minus
N2 OP50 plate
maco-1 PA14 plate
minus
maco-1 OP50 plate
replicate #1
value
0.8
0.6
0.4
0.2
0.0
−0.2
N2 OP50 plate
N = 10
N2 PA14 plate
N = 10
maco-1
OP50 plate
N = 10
maco-1
PA14 plate
N = 10
Mean difference
1.0
0.8
0.6
0.4
0.2
0.0
N2 PA14 plate
minus
N2 OP50 plate
maco-1 PA14 plate
minus
maco-1 OP50 plate
replicate #2
value
1.00
0.75
0.50
0.25
0.00
−0.25
N2 OP50 plate
N = 10
N2 PA14 plate
N = 10
maco-1
OP50 plate
N = 10
maco-1
PA14 plate
N = 10
Mean difference
1.2
1.0
0.8
0.6
0.4
0.2
0.0
N2 PA14 plate
minus
N2 OP50 plate
maco-1 PA14 plate
minus
maco-1 OP50 plate
replicate #3
value
1.00
0.75
0.50
0.25
0.00
−0.25
N2 OP50 plate
N = 10
N2 PA14 plate
N = 10
maco-1
OP50 plate
N = 10
maco-1
PA14 plate
N = 10
Mean difference
1.2
1.0
0.8
0.6
0.4
0.2
0.0
N2 PA14 plate
minus
N2 OP50 plate
maco-1 PA14 plate
minus
maco-1 OP50 plate

7A

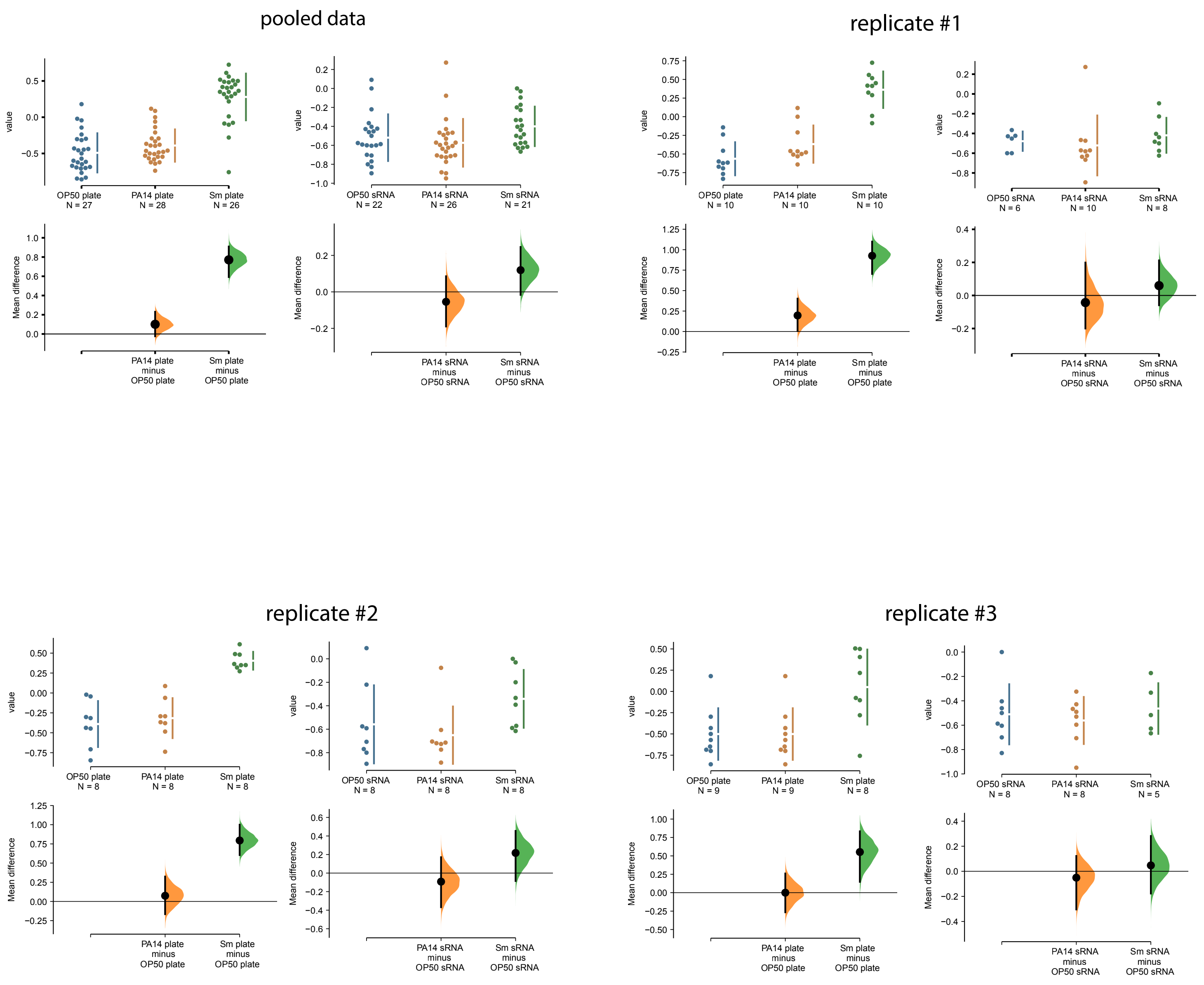
pooled data
replicate #1
replicate #2
replicate #3
value
Mean difference
OP50 plate
PA14 plate
Sm plate
OP50 sRNA
PA14 sRNA
Sm sRNA
PA14 plate minus OP50 plate
Sm plate minus OP50 plate
PA14 sRNA minus OP50 sRNA
Sm sRNA minus OP50 sRNA

# 7B

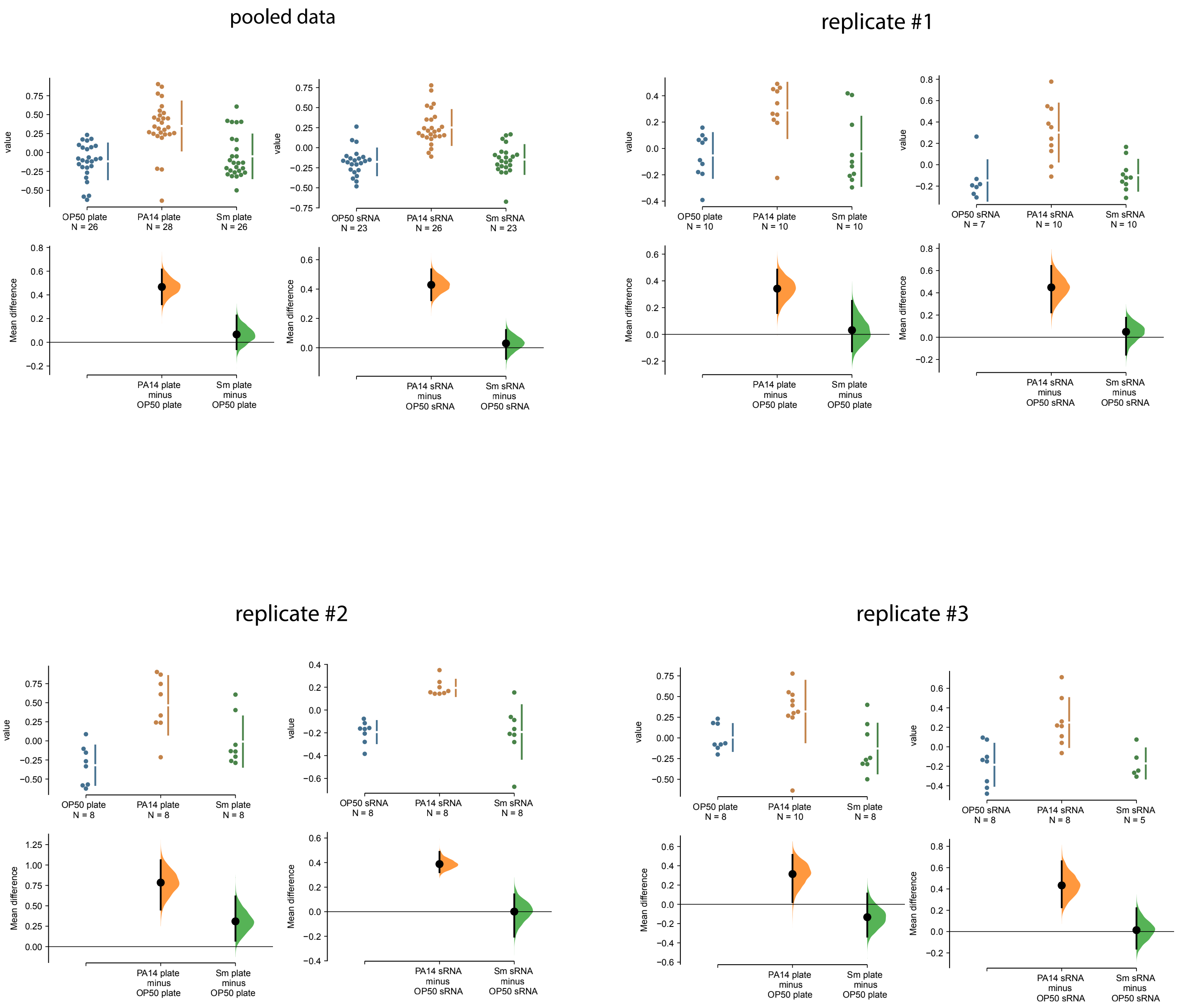
pooled data
replicate #1
replicate #2
replicate #3
value
Mean difference
OP50 plate N = 26
PA14 plate N = 28
Sm plate N = 26
OP50 sRNA N = 23
PA14 sRNA N = 26
Sm sRNA N = 23
OP50 plate N = 10
PA14 plate N = 10
Sm plate N = 10
OP50 sRNA N = 7
PA14 sRNA N = 10
Sm sRNA N = 10
OP50 plate N = 8
PA14 plate N = 8
Sm plate N = 8
OP50 sRNA N = 8
PA14 sRNA N = 8
Sm sRNA N = 8
OP50 plate N = 8
PA14 plate N = 10
Sm plate N = 8
OP50 sRNA N = 8
PA14 sRNA N = 8
Sm sRNA N = 5
PA14 plate minus OP50 plate
Sm plate minus OP50 plate
PA14 sRNA minus OP50 sRNA
Sm sRNA minus OP50 sRNA

7E

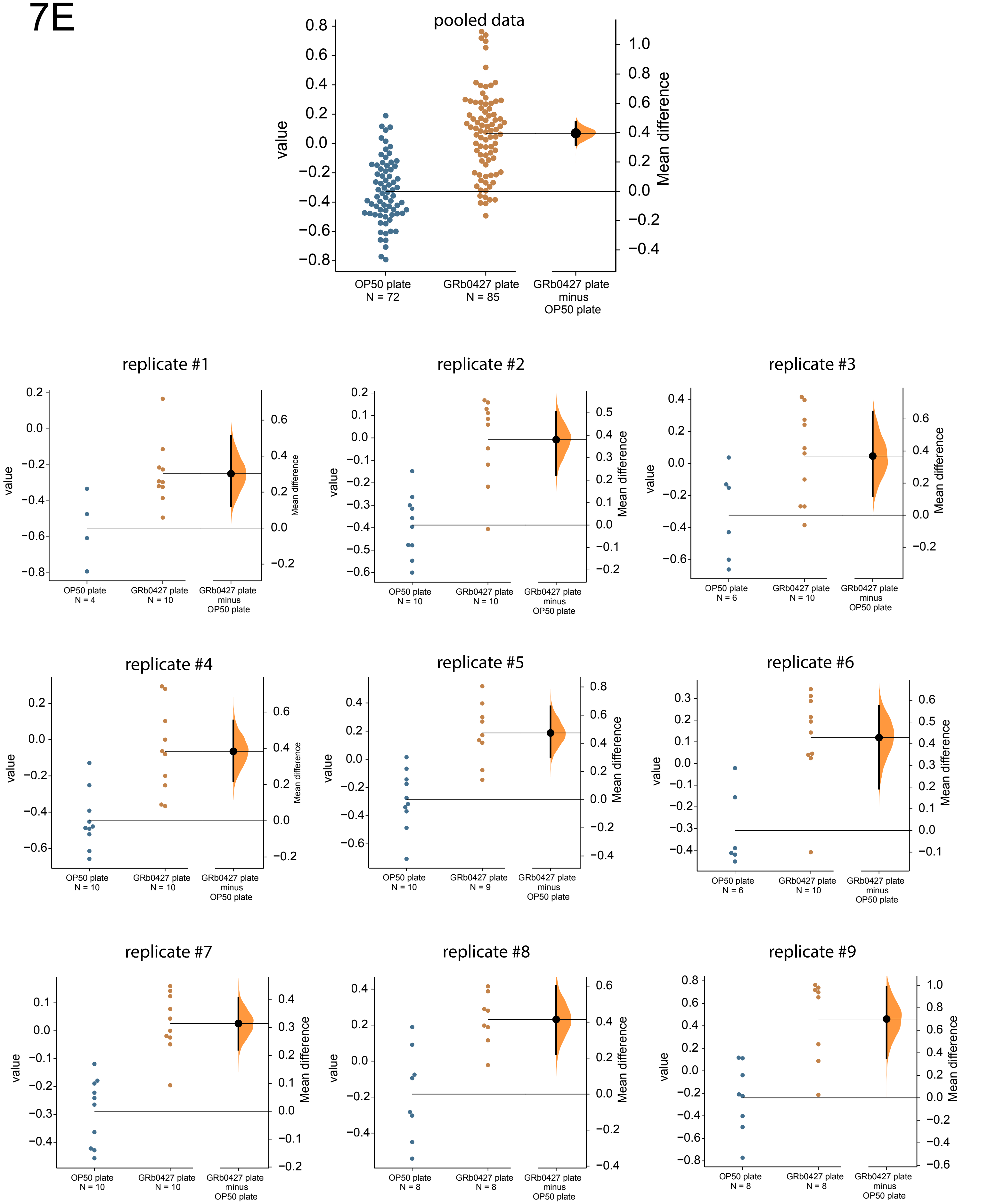

pooled data
value
Mean difference
OP50 plate
N = 72
GRb0427 plate
N = 85
GRb0427 plate
minus
OP50 plate
replicate #1
OP50 plate
N = 4
GRb0427 plate
N = 10
replicate #2
OP50 plate
N = 10
GRb0427 plate
N = 10
replicate #3
OP50 plate
N = 6
GRb0427 plate
N = 10
replicate #4
OP50 plate
N = 10
GRb0427 plate
N = 10
replicate #5
OP50 plate
N = 10
GRb0427 plate
N = 9
replicate #6
OP50 plate
N = 6
GRb0427 plate
N = 10
replicate #7
OP50 plate
N = 10
GRb0427 plate
N = 10
replicate #8
OP50 plate
N = 8
GRb0427 plate
N = 8
replicate #9
OP50 plate
N = 8
GRb0427 plate
N = 8

7F

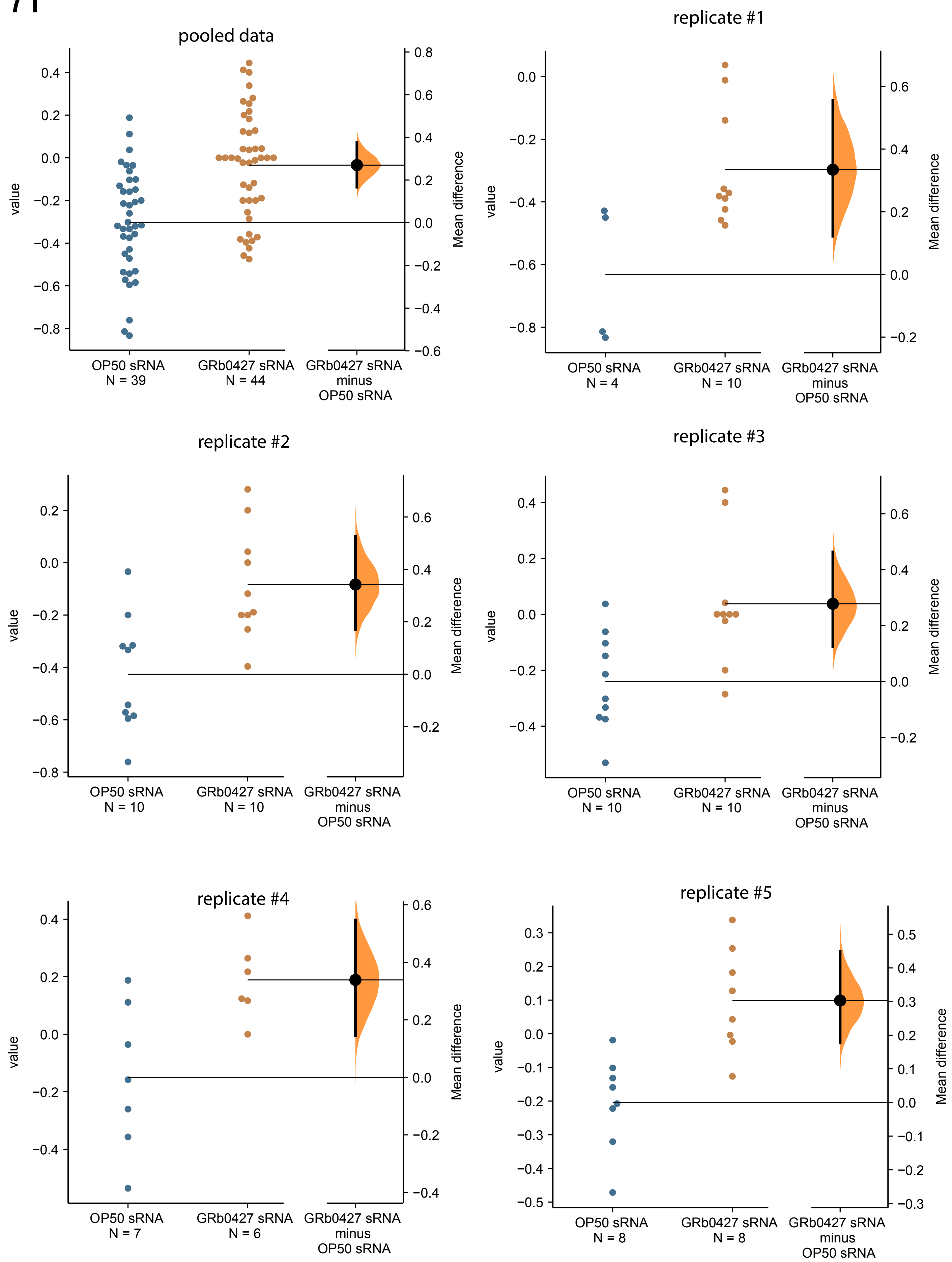
pooled data
value
Mean difference
OP50 sRNA
N = 39
GRb0427 sRNA
N = 44
GRb0427 sRNA
minus
OP50 sRNA
replicate #1
OP50 sRNA
N = 4
GRb0427 sRNA
N = 10
replicate #2
OP50 sRNA
N = 10
GRb0427 sRNA
N = 10
replicate #3
OP50 sRNA
N = 10
GRb0427 sRNA
N = 10
replicate #4
OP50 sRNA
N = 7
GRb0427 sRNA
N = 6
replicate #5
OP50 sRNA
N = 8
GRb0427 sRNA
N = 8

7G

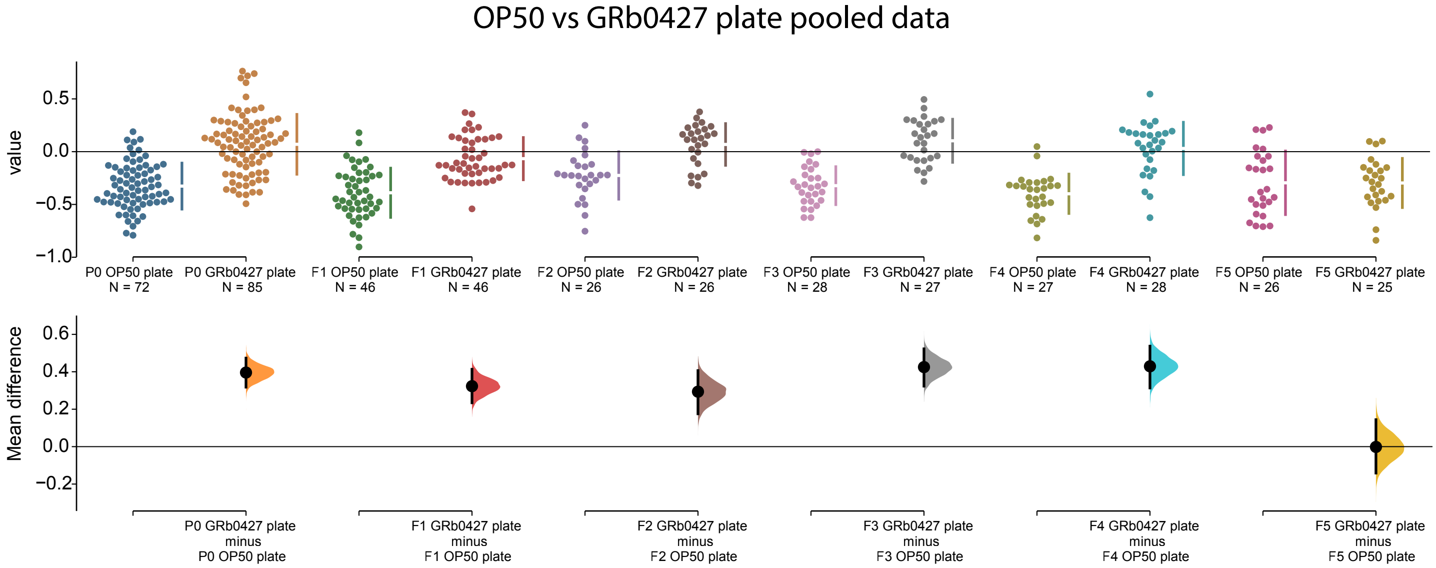
OP50 vs GRb0427 plate pooled data
value
Mean difference
P0 OP50 plate N = 72
P0 GRb0427 plate N = 85
F1 OP50 plate N = 46
F1 GRb0427 plate N = 46
F2 OP50 plate N = 26
F2 GRb0427 plate N = 26
F3 OP50 plate N = 28
F3 GRb0427 plate N = 27
F4 OP50 plate N = 27
F4 GRb0427 plate N = 28
F5 OP50 plate N = 26
F5 GRb0427 plate N = 25
P0 GRb0427 plate minus P0 OP50 plate
F1 GRb0427 plate minus F1 OP50 plate
F2 GRb0427 plate minus F2 OP50 plate
F3 GRb0427 plate minus F3 OP50 plate
F4 GRb0427 plate minus F4 OP50 plate
F5 GRb0427 plate minus F5 OP50 plate

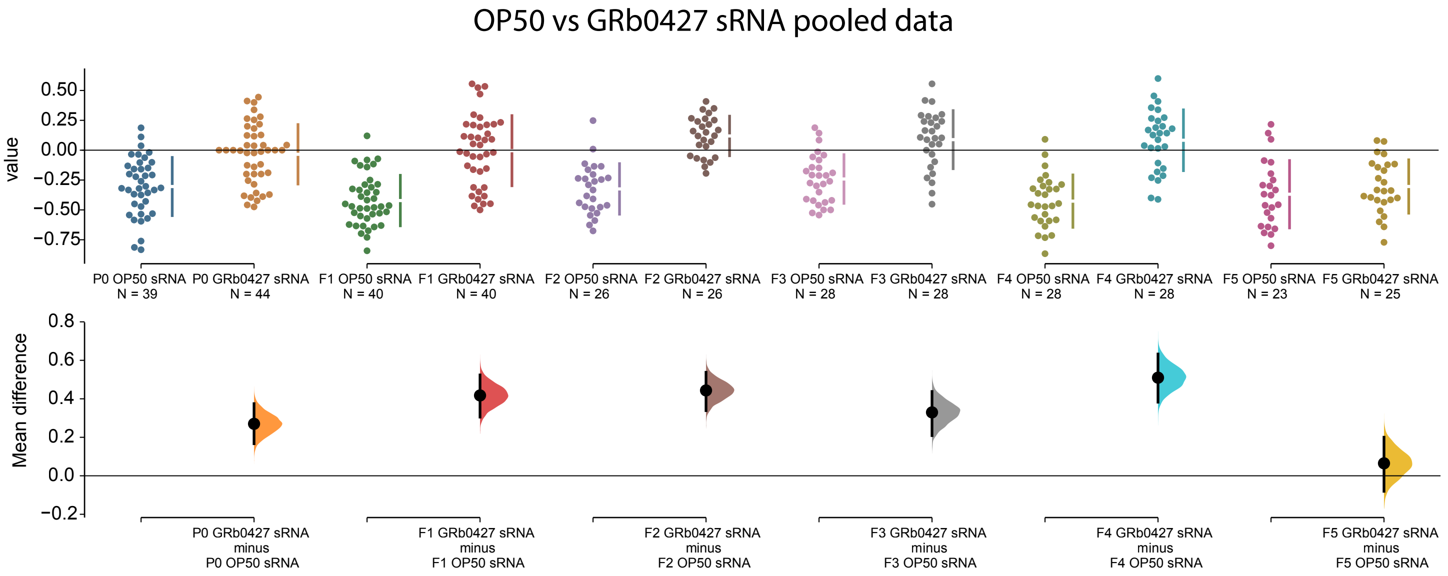
OP50 vs GRb0427 sRNA pooled data
value
Mean difference
P0 OP50 sRNA N = 39
P0 GRb0427 sRNA N = 44
F1 OP50 sRNA N = 40
F1 GRb0427 sRNA N = 40
F2 OP50 sRNA N = 26
F2 GRb0427 sRNA N = 26
F3 OP50 sRNA N = 28
F3 GRb0427 sRNA N = 28
F4 OP50 sRNA N = 28
F4 GRb0427 sRNA N = 28
F5 OP50 sRNA N = 23
F5 GRb0427 sRNA N = 25
P0 GRb0427 sRNA minus P0 OP50 sRNA
F1 GRb0427 sRNA minus F1 OP50 sRNA
F2 GRb0427 sRNA minus F2 OP50 sRNA
F3 GRb0427 sRNA minus F3 OP50 sRNA
F4 GRb0427 sRNA minus F4 OP50 sRNA
F5 GRb0427 sRNA minus F5 OP50 sRNA

# 7H

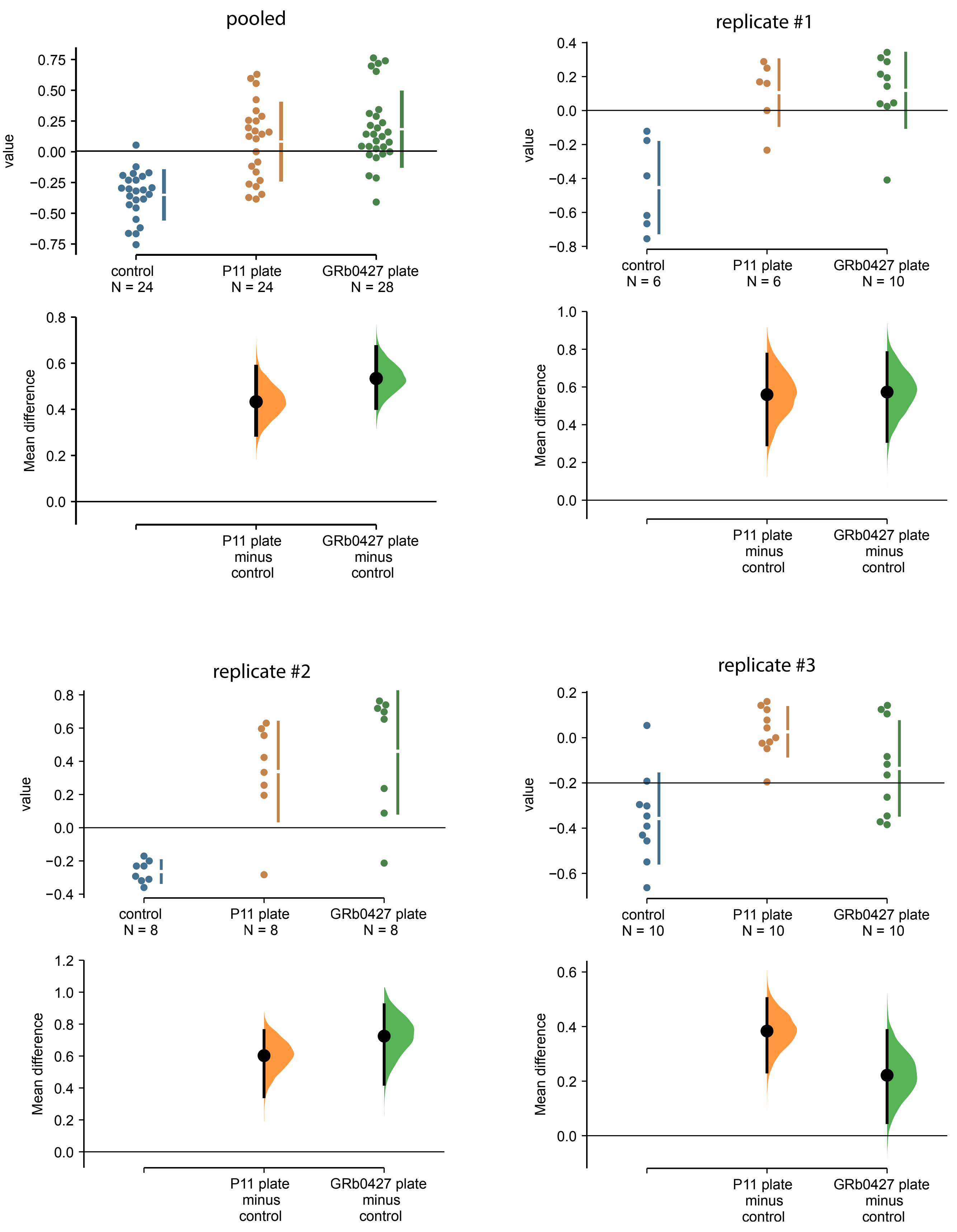

pooled
replicate #1
replicate #2
replicate #3
value
Mean difference
control
N = 24
P11 plate
N = 24
GRb0427 plate
N = 28
control
N = 6
P11 plate
N = 6
GRb0427 plate
N = 10
control
N = 8
P11 plate
N = 8
GRb0427 plate
N = 8
control
N = 10
P11 plate
N = 10
GRb0427 plate
N = 10
P11 plate
minus
control
GRb0427 plate
minus
control

7l

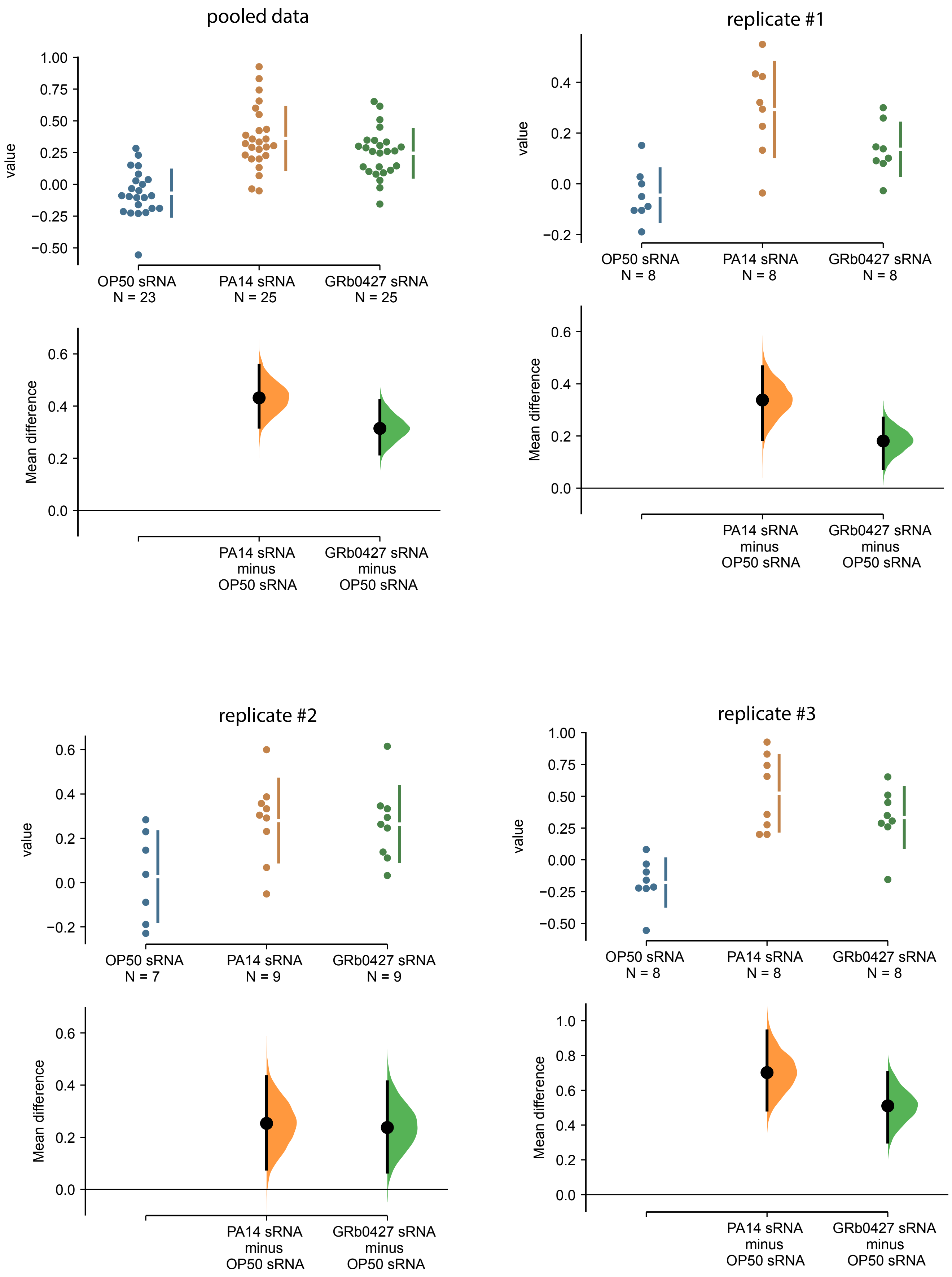
pooled data
value
OP50 sRNA
N = 23
PA14 sRNA
N = 25
GRb0427 sRNA
N = 25
Mean difference
PA14 sRNA
minus
OP50 sRNA
GRb0427 sRNA
minus
OP50 sRNA
replicate #1
N = 8
replicate #2
N = 7
N = 9
replicate #3

7J

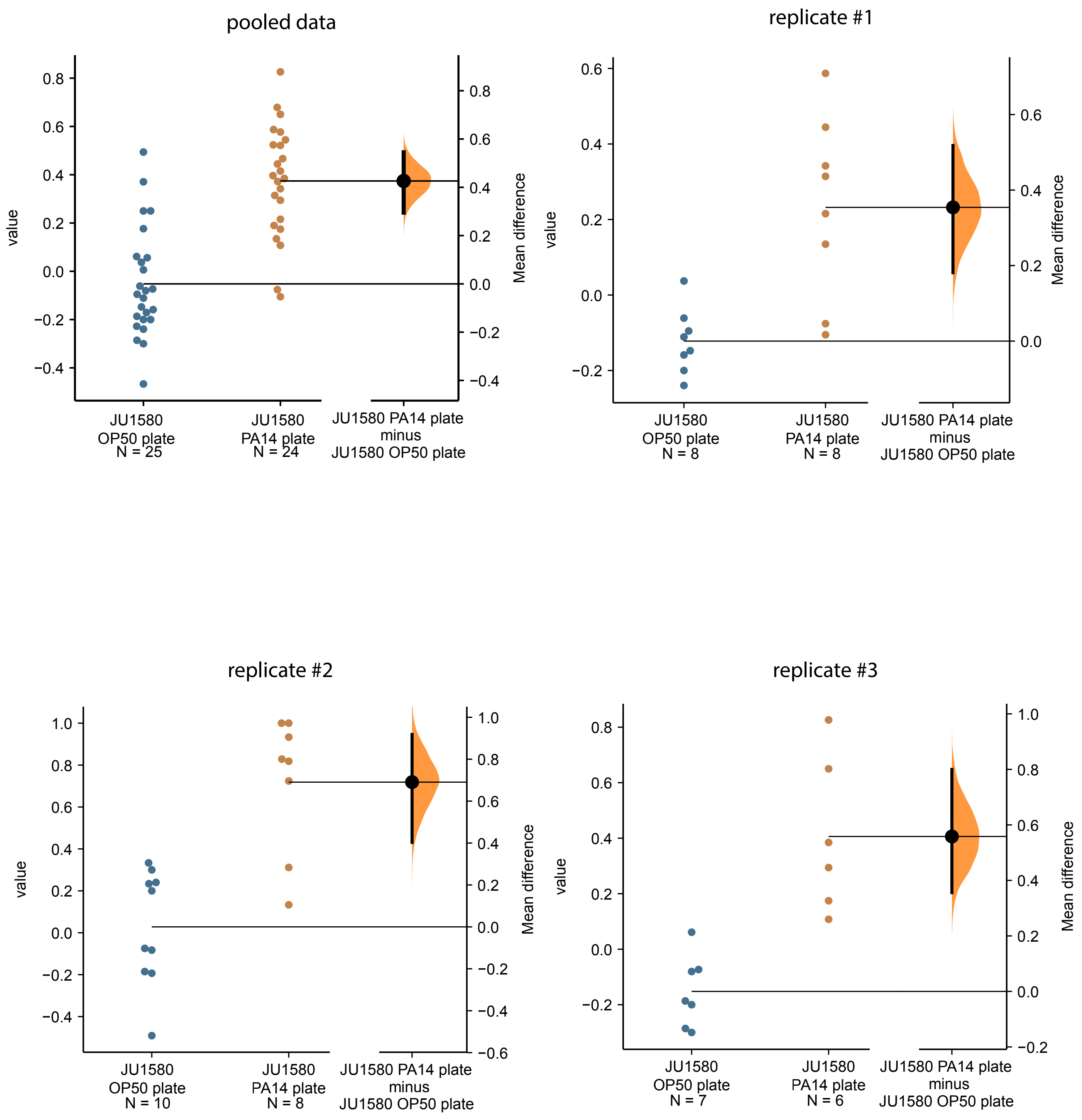
pooled data
value
Mean difference
JU1580 OP50 plate N = 25
JU1580 PA14 plate N = 24
JU1580 PA14 plate minus JU1580 OP50 plate
replicate #1
value
Mean difference
JU1580 OP50 plate N = 8
JU1580 PA14 plate N = 8
JU1580 PA14 plate minus JU1580 OP50 plate
replicate #2
value
Mean difference
JU1580 OP50 plate N = 10
JU1580 PA14 plate N = 8
JU1580 PA14 plate minus JU1580 OP50 plate
replicate #3
value
Mean difference
JU1580 OP50 plate N = 7
JU1580 PA14 plate N = 6
JU1580 PA14 plate minus JU1580 OP50 plate

7K

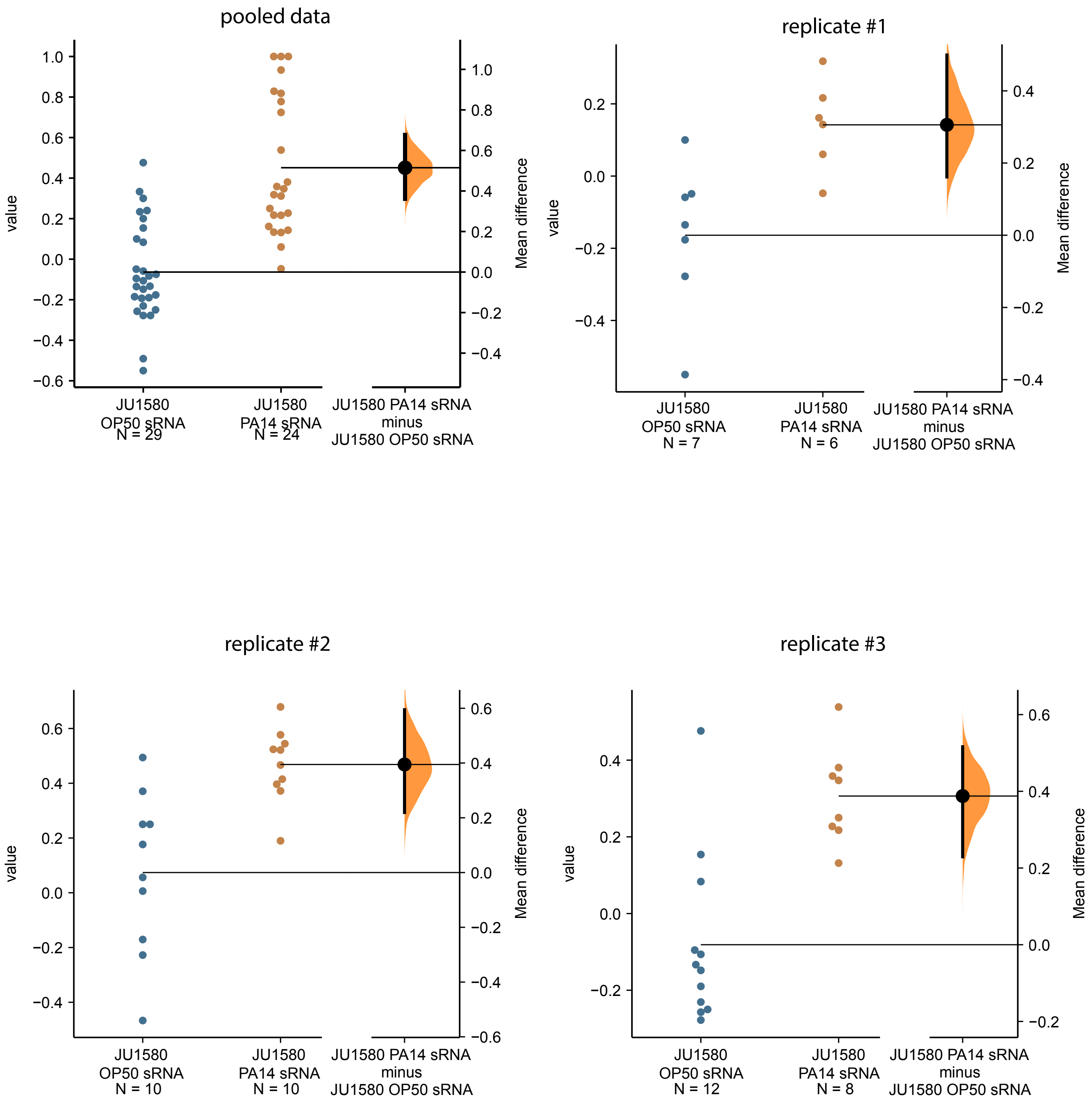
pooled data
replicate #1
replicate #2
replicate #3
value
Mean difference
JU1580 OP50 sRNA N = 29
JU1580 PA14 sRNA N = 24
JU1580 PA14 sRNA minus JU1580 OP50 sRNA
JU1580 OP50 sRNA N = 7
JU1580 PA14 sRNA N = 6
JU1580 OP50 sRNA N = 10
JU1580 PA14 sRNA N = 10
JU1580 OP50 sRNA N = 12
JU1580 PA14 sRNA N = 8

7L

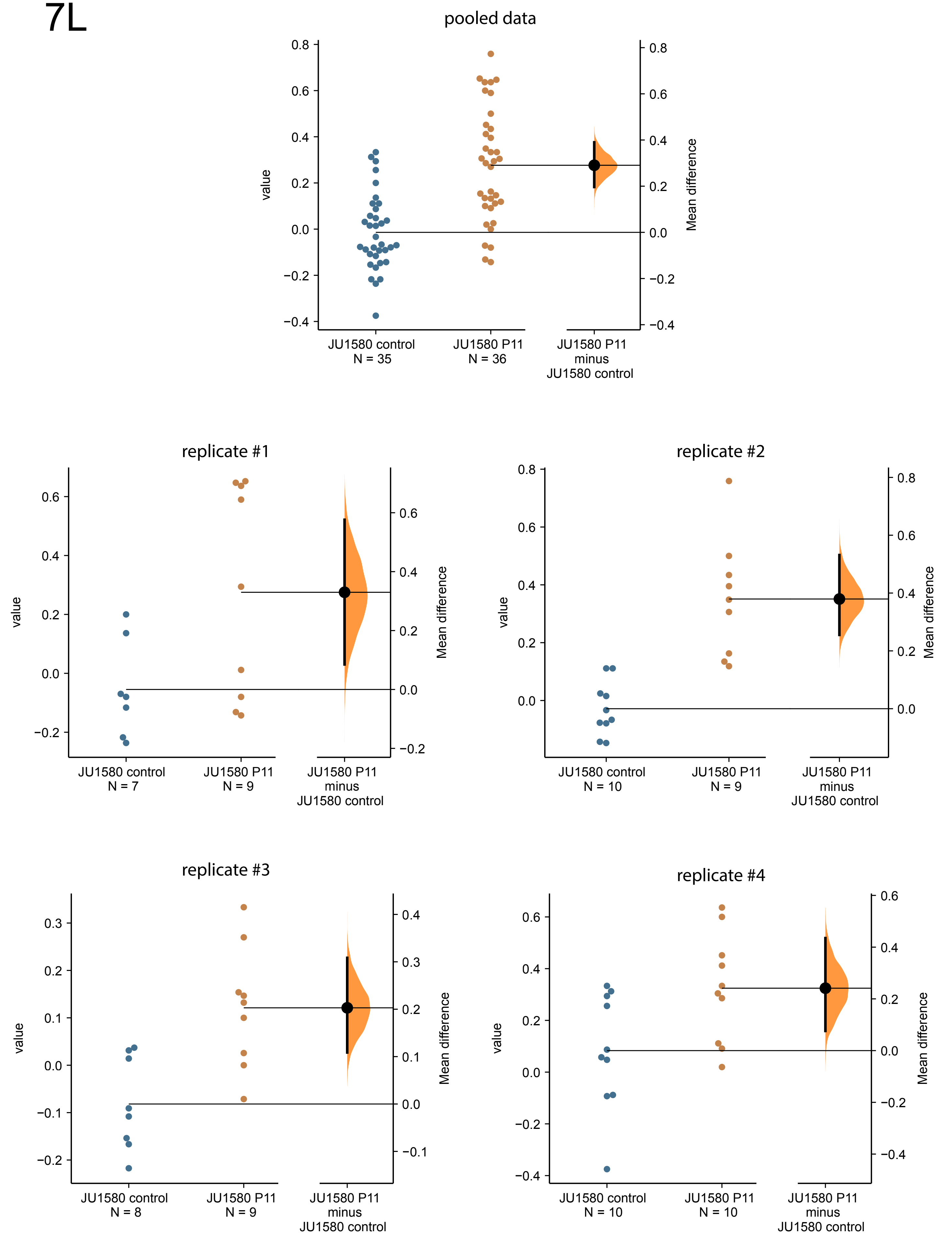
pooled data
value
Mean difference
JU1580 control
N = 35
JU1580 P11
N = 36
JU1580 P11
minus
JU1580 control
replicate #1
value
Mean difference
JU1580 control
N = 7
JU1580 P11
N = 9
JU1580 P11
minus
JU1580 control
replicate #2
value
Mean difference
JU1580 control
N = 10
JU1580 P11
N = 9
JU1580 P11
minus
JU1580 control
replicate #3
value
Mean difference
JU1580 control
N = 8
JU1580 P11
N = 9
JU1580 P11
minus
JU1580 control
replicate #4
value
Mean difference
JU1580 control
N = 10
JU1580 P11
N = 10
JU1580 P11
minus
JU1580 control

# 7M

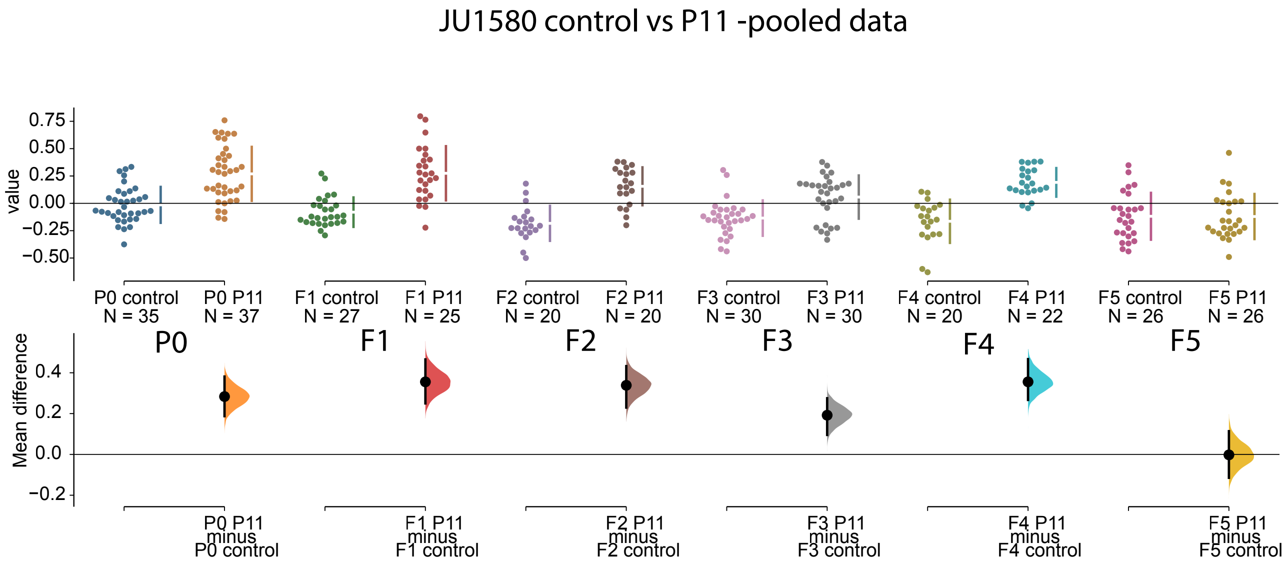

JU1580 control vs P11 -pooled data
value
0.75
0.50
0.25
0.00
−0.25
−0.50
P0 control
N = 35
P0 P11
N = 37
F1 control
N = 27
F1 P11
N = 25
F2 control
N = 20
F2 P11
N = 20
F3 control
N = 30
F3 P11
N = 30
F4 control
N = 20
F4 P11
N = 22
F5 control
N = 26
F5 P11
N = 26
P0
F1
F2
F3
F4
F5
Mean difference
0.4
0.2
0.0
−0.2
P0 P11
minus
P0 control
F1 P11
minus
F1 control
F2 P11
minus
F2 control
F3 P11
minus
F3 control
F4 P11
minus
F4 control
F5 P11
minus
F5 control

7N

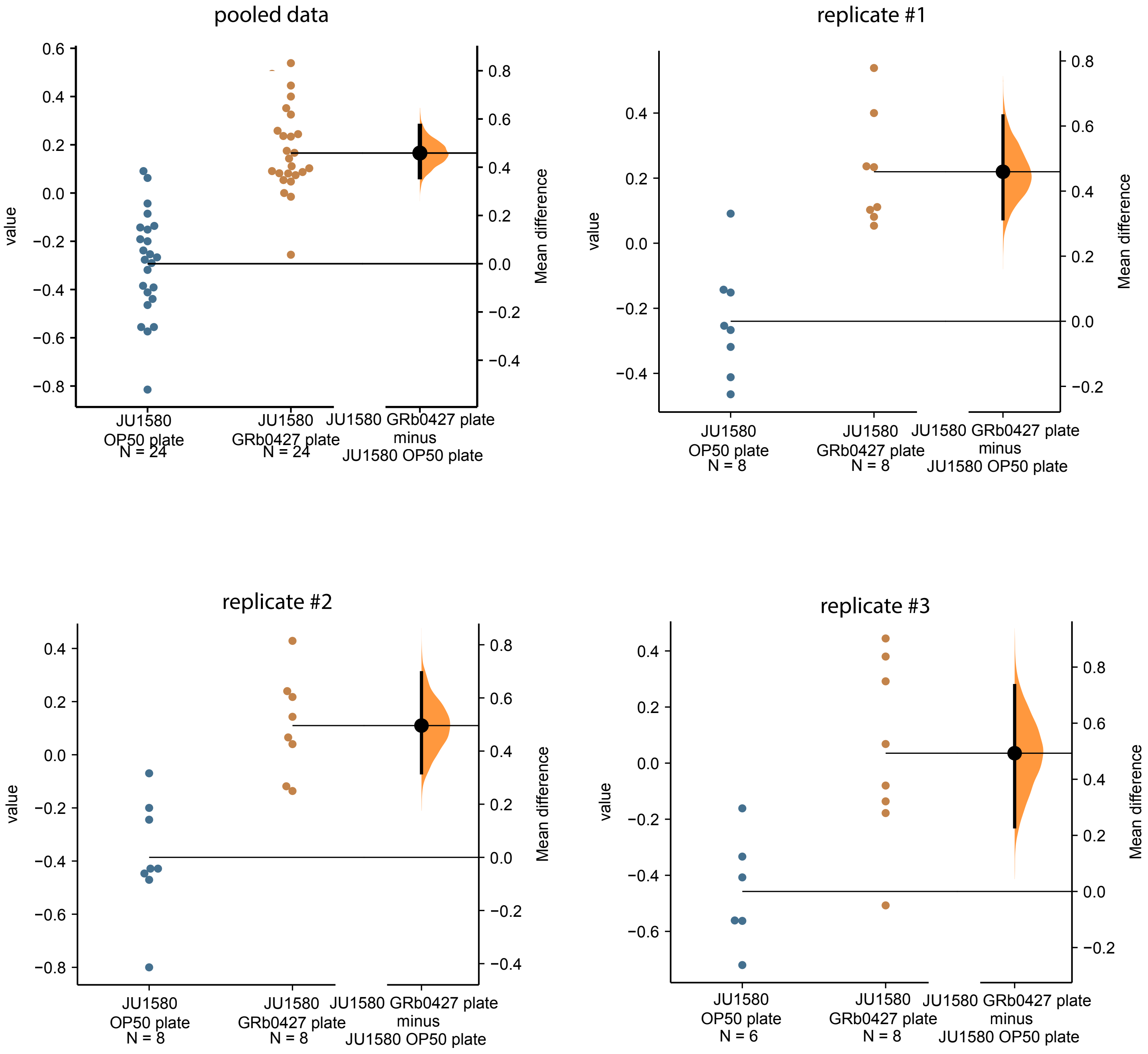
pooled data
replicate #1
replicate #2
replicate #3
value
Mean difference
JU1580 OP50 plate N = 24
JU1580 GRb0427 plate N = 24
JU1580 GRb0427 plate minus JU1580 OP50 plate
JU1580 OP50 plate N = 8
JU1580 GRb0427 plate N = 8
JU1580 OP50 plate N = 6

7O

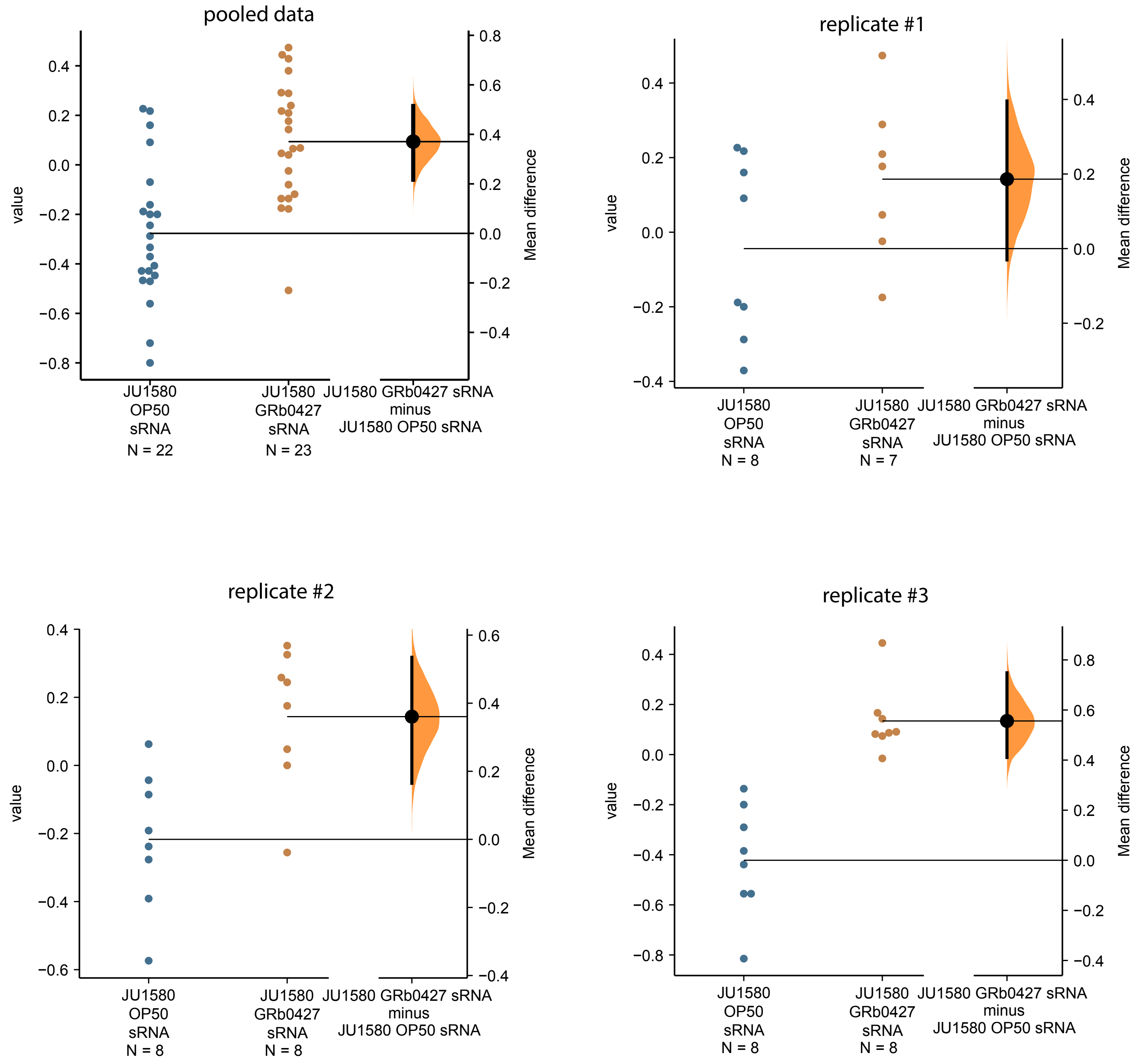
pooled data
value
Mean difference
JU1580 OP50 sRNA N = 22
JU1580 GRb0427 sRNA N = 23
JU1580 GRb0427 sRNA minus JU1580 OP50 sRNA
replicate #1
JU1580 OP50 sRNA N = 8
JU1580 GRb0427 sRNA N = 7
replicate #2
JU1580 OP50 sRNA N = 8
JU1580 GRb0427 sRNA N = 8
replicate #3
JU1580 OP50 sRNA N = 8
JU1580 GRb0427 sRNA N = 8
